# The unique neuronal structure and neuropeptide repertoire in the ctenophore *Mnemiopsis leidyi* shed light on the evolution of animal nervous systems

**DOI:** 10.1101/2021.03.31.437758

**Authors:** Maria Y Sachkova, Eva-Lena Nordmann, Joan J Soto-Àngel, Yasmin Meeda, Bartłomiej Górski, Benjamin Naumann, Daniel Dondorp, Marios Chatzigeorgiou, Maike Kittelmann, Pawel Burkhardt

## Abstract

The ctenophore nerve net represents one of the earliest evolved nervous system of animals. Due to the uncertainties of their phylogenetic placement of ctenophores and the absence of several key bilaterian neuronal genes, it has been hypothesized that their neurons have evolved independently. Whether this is indeed the case remains unclear, and thus the evolutionary history of neurons is still contentious. Here, we have characterized the neuropeptide repertoire of the ctenophore *Mnemiopsis leidyi*. Using the machine learning NeuroPID tool^1^ 129 new putative neuropeptide precursors were predicted. Sixteen of them are detected in the subepithelial nerve net (SNN), aboral organ (AO) and epithelial sensory cells (ESC) of early cydippid-stage *M. leidyi* by in situ hybridization (ISH) and immunohistochemistry (IHC). Four of these neuropeptides increase the animals’ swimming velocity in a behavioral essay. The new neuropeptides were used as markers to identify neuronal cell types in single cell transcriptomic data^2^. To unravel the neuronal architecture, we 3D reconstructed the SNN underlying the comb plates using serial block-face scanning electron microscopy (SBF-SEM). For the first time, we confirm a more than 100 years old hypothesis about anastomoses between neurites of the same cell in ctenophores and reveal that they occur through a continuous membrane. Our findings reveal the unique neuronal structure and neuropeptide repertoire of ctenophores and are important for reconstructing the evolutionary origin of animal neurons and nervous systems.

## Introduction

The question how the earliest animals sensed and reacted to their environment has fascinated humans from Greek naturalists like Aristotle^3^ to the evolutionary biologist Charles Darwin^4^. Neuronal cell types organized into a nervous system via synaptic cell contacts appear to be crucial for the evolution of large, structured bodies and complex behaviors as present in many animal lineages. Recently, it was shown that many components of the synaptic tool kit were present in the last common ancestor of animals (Metazoa) and unicellular choanoflagellates^5,6^. Traditionally, it is thought that the metazoan last common ancestor lacked specialized neuronal cell types and a nervous system^7^, a condition present in extant sponges and Placozoans^8^. Later, a first net-like nervous system evolved as present in cnidarians and ctenophores followed by an increased condensation and diversification in the bilaterian lineage^9^. However, this scenario has been challenged by recent uncertainties about the phylogenetic placement of the ctenophores (**Fig. 1A**).

**Figure 1.**
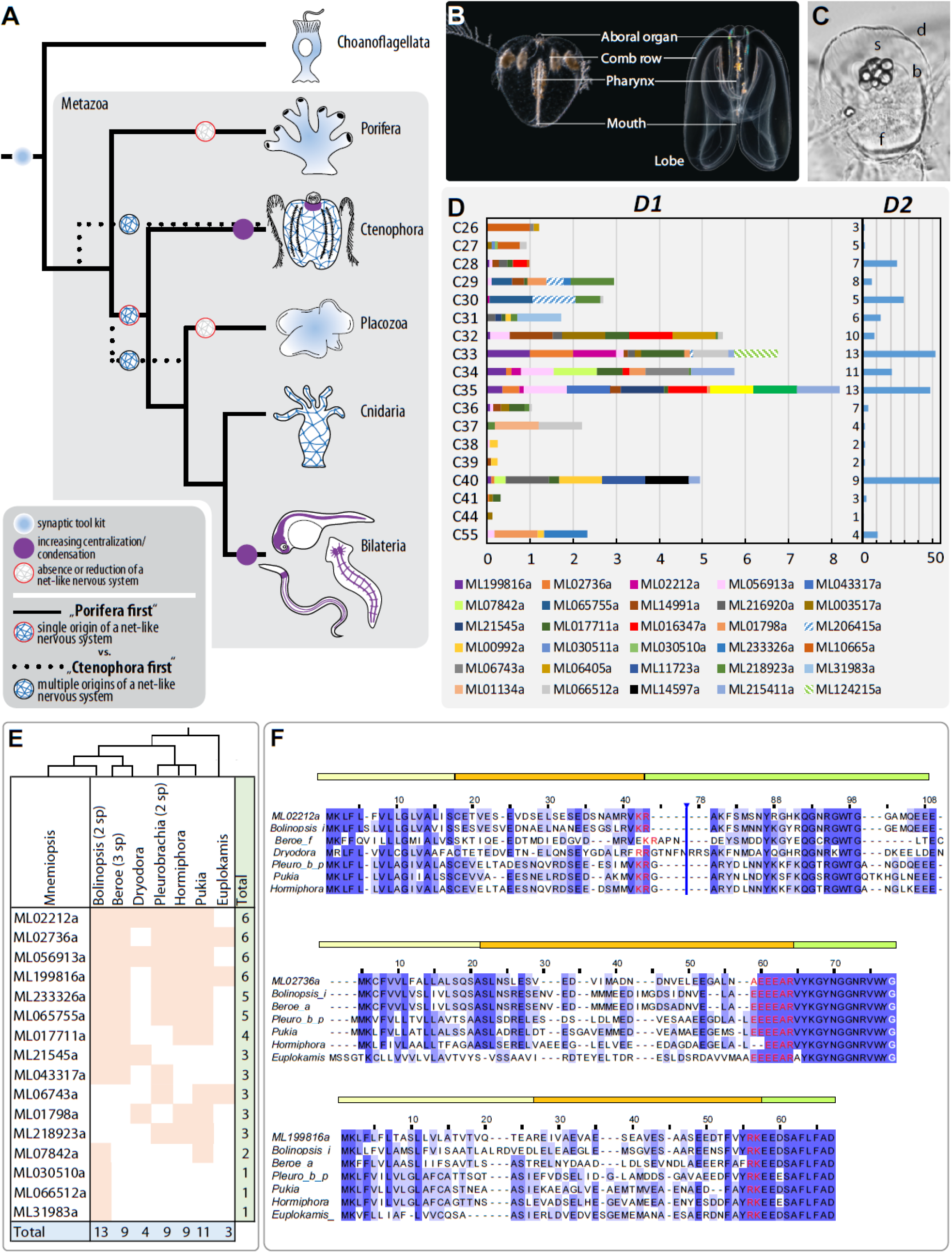
Evolution of the nervous system(s) and identification of putative neuropeptides in the ctenophore M. leidyi. (**A**) Metazoan phylogeny and possible scenarios of nervous system origin. (**B**) Body plan of the ctenophore M. leidyi: cydippid-stage (left) and lobed adult (right); note that the general anatomy is very similar. (**C**) Aboral organ of the cydippid-stage; d – dome cilia, s – statolith, b – balancer cilia, f – floor of the aboral organ. (**D**) Putative neuropeptide precursors enriched in uncharacterised metacells^2^. **D1** – normalised expression levels above 0.05; **D2** – total neuropeptide expression levels in each metacell (summed Umifrac values). (**E**) Distribution of neuropeptide homologs among Ctenophora. (**F**) Alignments of three M. leidyi precursors. Accession numbers are available in the Supplementary Table 3.; in case several sequences were identified in the same genus, only one of them is shown. Vertical blue bars mark hidden columns in alignment gaps in the M. leidyi sequences. Shades of blue represent conservation rate, residues in red correspond to putative processing sites, Gly residues converted into C-terminal amide are in white. Above each alignment the precursor structure is shown: yellow – signal peptide, orange – propeptide, green – mature peptide.

Ctenophores, or comb jellies, are gelatinous marine invertebrates that move by beating unique ciliary combs and catch prey with adhesive colloblasts on their tentacles (**Fig. 1B**). They exhibit a well-developed nervous system consisting of an aboral-sensory organ (AO) (**Fig. 1C**), two adjacent polar fields, potential mesogleal neurons, a polygonal subepithelial nerve net (SNN) and a variety of ciliated and non-ciliated sensory cells all over the epidermis, pharynx, and tentacles^10-13^. For decades, based on morphological features, these animals were grouped with cnidarias in a clade of Coelenterata^11^ or placed as a sister group to bilaterians^14^, while sponges were considered as the earliest branching animal lineage. Yet, sequenced genomes of the ctenophores *Pleurobrachia bachei* and *Mnemiopsis leidyi* placed them as the earliest or one of the earliest branching metazoan lineages^15,16^ (**Fig. 1A**).

The first nervous system might have heavily relied on paracrine signaling using neuropeptide-like molecules as transmitters^15,17,18^. Neuropeptides are present in cnidarians, bilaterians^18,19^ and neuron-less Placozoa^20,21^ but have not been found in sponges so far. In ctenophores, 72 putative prohormones were identified in the genome of *P. bachei*^15^ and immunoreactivity against FMRFamide and vasopressin was reported in the closely related *P. pileus*^12^. However, genes corresponding to these two molecules have not been found in *P. bachei*^15^ and only a single neuropeptide homolog has been found in the genome of *M. leidyi*^22^. The orthologues of genes involved in the multi-step processing pathway of bilaterian neuropeptides^23^ gives further hints for neuropeptide signaling in ctenophores^15,24^. Neuropeptides are fast evolving resulting in many taxon-specific neuropeptides^19,25^ and complicating the identification of putative homologs, especially in highly diverged groups like ctenophores^1,15^. Thus, detailed data on the identity and function of neuropeptides as well as neuronal morphology in the ctenophore nervous system are essential to understanding key events in the origin and diversification of animal nervous system(s).

## Results

### Ctenophores express diverse and unique neuropeptide precursors

Because neuropeptides are short and highly divergent molecules complicating similarity-based searches, we mined predicted protein models^26,27^ of *M. leidyi* de-novo using the NeuroPID tool^1^ (**Supplementary Fig. 1**). We were able to predict 129 putative neuropeptide precursors including both linear and disulphide-rich sequences (**Supplementary Table 1**). Twenty seven out of 67 putative linear precursors were the best hits, satisfying all four machine learning models (**Supplementary Table 2**). Because neurons could not be identified in an earlier single-cell transcriptome analysis of *M. leidyi* we hypothesized that some of the uncharacterised metacell clusters (**Supplementary Fig. 2**, C26-C41, C44)^28^ could correspond to neurons enriched in our newly identified putative neuropeptides. Indeed, mapping these putative neuropeptides onto the metacell clusters we found that 30 of them enriched in one or several clusters (**Fig. 1D, Supplementary Table 1**). From these, C33 to C35 and C40 co-express the highest number of putative neuropeptides (nine up to 13, **Fig. 1D**) and show the highest neuropeptide Umifrac values (21 to 55). However, neuropeptide precursors undergo multi-step processing to become mature functional neuropeptides. We therefore predicted mature neuropeptides from the identified precursors based on specific assumptions for their processing (**Supplementary Fig. 3**)^29,30^. We were able to predict mature peptides for 16 precursors, while for five, the processing was unclear. Blast search and sequence alignment among 11 available ctenophore transcriptomes and genomes resulted in at least one homolog for 16 of the putative neuropeptide precursors (**Fig. 1E, F, Supplementary Fig. 3 and Supplementary Table 3**). Consistent with a previous study^15^, none of these putative neuropeptides could be homologized with neuropeptides of other metazoan groups using Protein Blast.

### Ctenophore-specific neuropeptides are expressed in various cells of the neuro-sensory system

To further characterize the expression profiles and protein functions we selected genes that are expressed between 10 and 20 hours post fertilization correlating with neural development in *M. leidyi* ^31,32^ (**Supplementary Fig. 5**). The expression of two groups of genes (group 1: ML218923a, ML31983a, ML01134a, ML066512a, ML14597a and group 2: ML06743a, ML06405a, ML11723a) does not correlate with estimated neural development.^32^ These genes might be involved in early embryonic development (group 1) or adult homeostasis (group 2). The remaining genes were expressed in cells of various parts of the nervous system.

ISH in three to four days post fertilisation (dpf) cydippids showed that ML199816a, ML02212a, ML02736a and ML056913a are expressed in neuronal cell bodies and neurites of the SNN (**Fig. 2A, Supplementary Fig. 6, 7A-H, 8**), particularly notable at the base of the comb plates (**Fig. 2B**). Additionally, ML199816a-positive neurites are also located inside the tentacles (**Fig. 2D**). ML02212a and ML056913a are expressed in AO sensory cells (**Fig. 2C, Supplementary Fig. 8**). These genes were also detected in ESC and gut cells, tentacle and tentacle bulb cells (**Supplementary Fig. 7, 8**). An antibody raised against ML02212 mature peptide matches the ISH pattern in the AO, tentacle bulbs, outer epithelium (in particular around the comb plates), and in the pharyngeal epithelium (**Fig. 2B-D, Supplementary Fig. 6** and **Supplementary Video 1**). Surprisingly, we found ML199816a-, ML02212a- and ML02736a-positive neurites of the SNN innervating the AO floor (**Supplementary Fig. 6, 7**).

**Figure 2.**
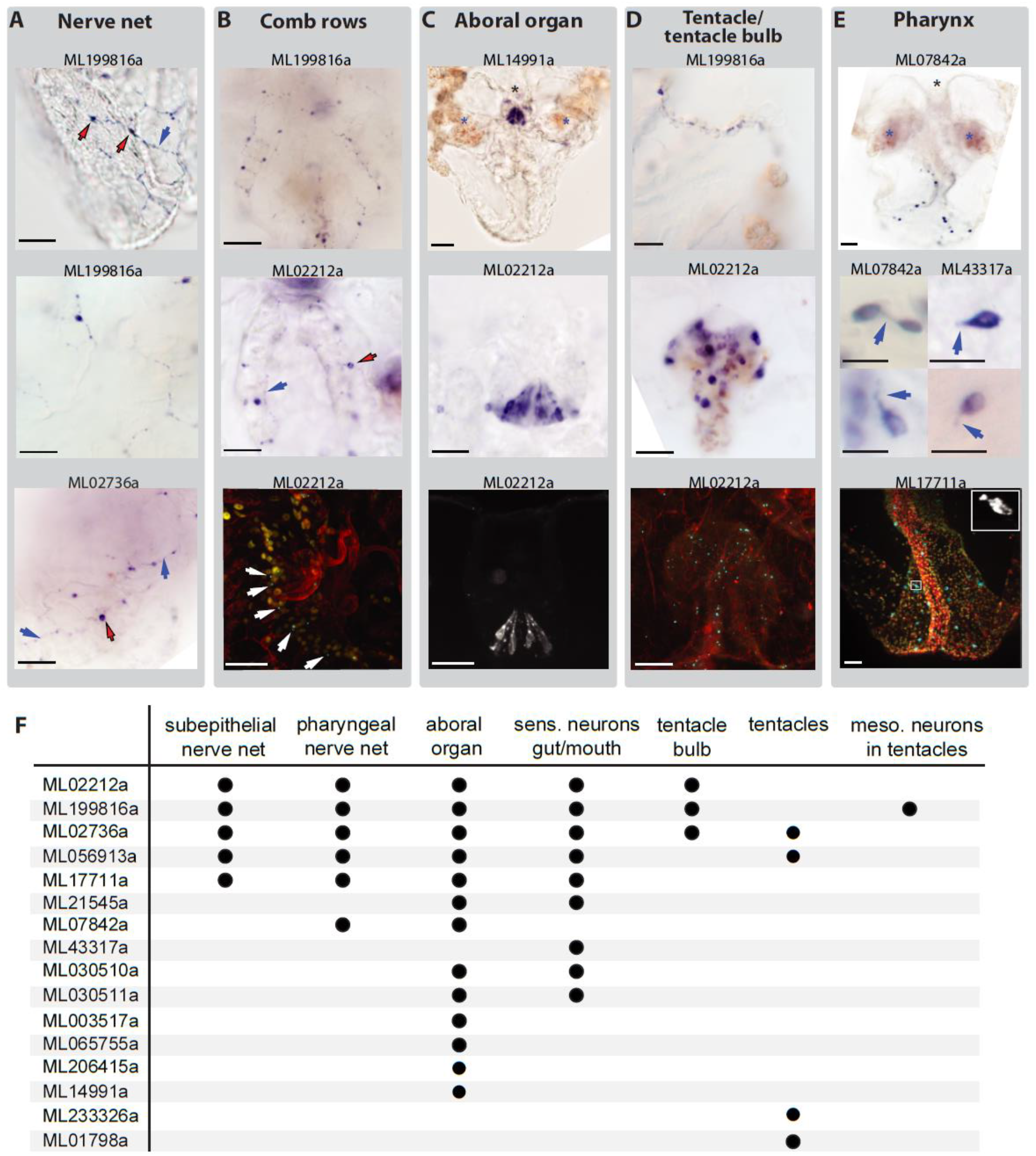
Putative neuropeptides are expressed in various cells of the nervous system of M. leidyi. (**A, B**) Neuropeptide expression in the sub-epithelial nerve net: general view (**A**) and around comb rows (**B**); red arrows – cell bodies, blue arrows-neurites. (**C-E**) Neuropeptide expression in the in cells of the aboral organ (**C**), in tentacles/tentacle bulbs (**D**), and in the pharynx (**E;** for ML07842, arrows indicate cell protrusions; for ML43317a, arrows indicate cilia). Asterisks indicate AO. Staining in blue was done by ISH (**A-E** two upper rows and **A**, bottom row) and fluorescent staining is done by IHC (**B-E**, bottom row), (**F**) Summary of neuropeptide distribution.

ML065755a, ML14991a, ML003517a, and ML206415a are exclusively expressed in cells of the AO that show typical sensory cell morphology^33^ (**Supplementary Fig. 7I-U, 12)**. ML07842a, ML43317a, ML17711a, ML030510a and ML030511a are mostly expressed in cells of or surrounding the mouth opening and pharynx (**Fig. 2E, Supplementary Fig. 9, 10, 11, Supplementary Video 2, 6**). These cells have a variable morphology. ML07842a-positive cells have irregular forms (**Fig. 2E**) indicating that they have one to several protrusions. We confirmed the ISH pattern of this gene with antibodies raised against the mature protein. Additionally, to the ISH pattern the antibodies labeled four groups of cells present in the cydippid and adult AO and the AO rim of adults only (**Supplementary Fig. 10**). ML43317a and ML030510a-positive cells are ciliated cells widely distributed around the mouth opening an within the pharyngeal wall (**Fig. 2E**). Additionally, ML030510a and ML030511a are expressed in the rim of the AO (**Supplementary Fig. 9L, 10**). ML07842a-positive cells were also found in the adult (both ISH and IHC), but not in the cydippid stage AO (**Supplementary Fig. 8 & Supplementary Video 3, 4**). Both ISH and IHC showed that ML17711a and ML21545a are expressed in diverse cell types in the AO, the outer body surface, pharynx and around the mouth (**Supplementary Fig. 11, Supplementary Video 5, 6, 7**). ML233326a and ML01798a are detected in the tentacles only (**Supplementary Fig. 13**).

### Newly identified neuropeptides affect ctenophore swimming behavior

To test if neuropeptides effect the behavior we performed swimming experiments using cydippid-stage *M. leidyi* (4-5 dpf). Animals were placed into an small arena equipped with a camera system and aclimatized for 10 min (instant and median velocity values stabilize after five minutes, **Fig. 3, Supplementary Fig. 13**). We then added the synthetized peptide into the medium and recorded the velocity during two five minute intervalls (interval 1 and 2; **Fig. 3**). The median velocity of control and experimental specimens was then normalized against the median velocity before adding the peptide to account for individual behavioral differences. Indeed, ML056913a, ML065755a, ML43317a, and ML233326a increased the median velocities during interval 2 (p<0.05, Mann-Whitney U, **Fig. 3**). ML02212a showed a non-significant tendency to increase the median velocity during interval 1 (p=0.067, Mann-Whitney U, **Fig. 3D**). ML02736a, ML21545a, ML017711a, and ML01798a showed no effect (**Supplementary Fig. 8**).

**Figure 3.**
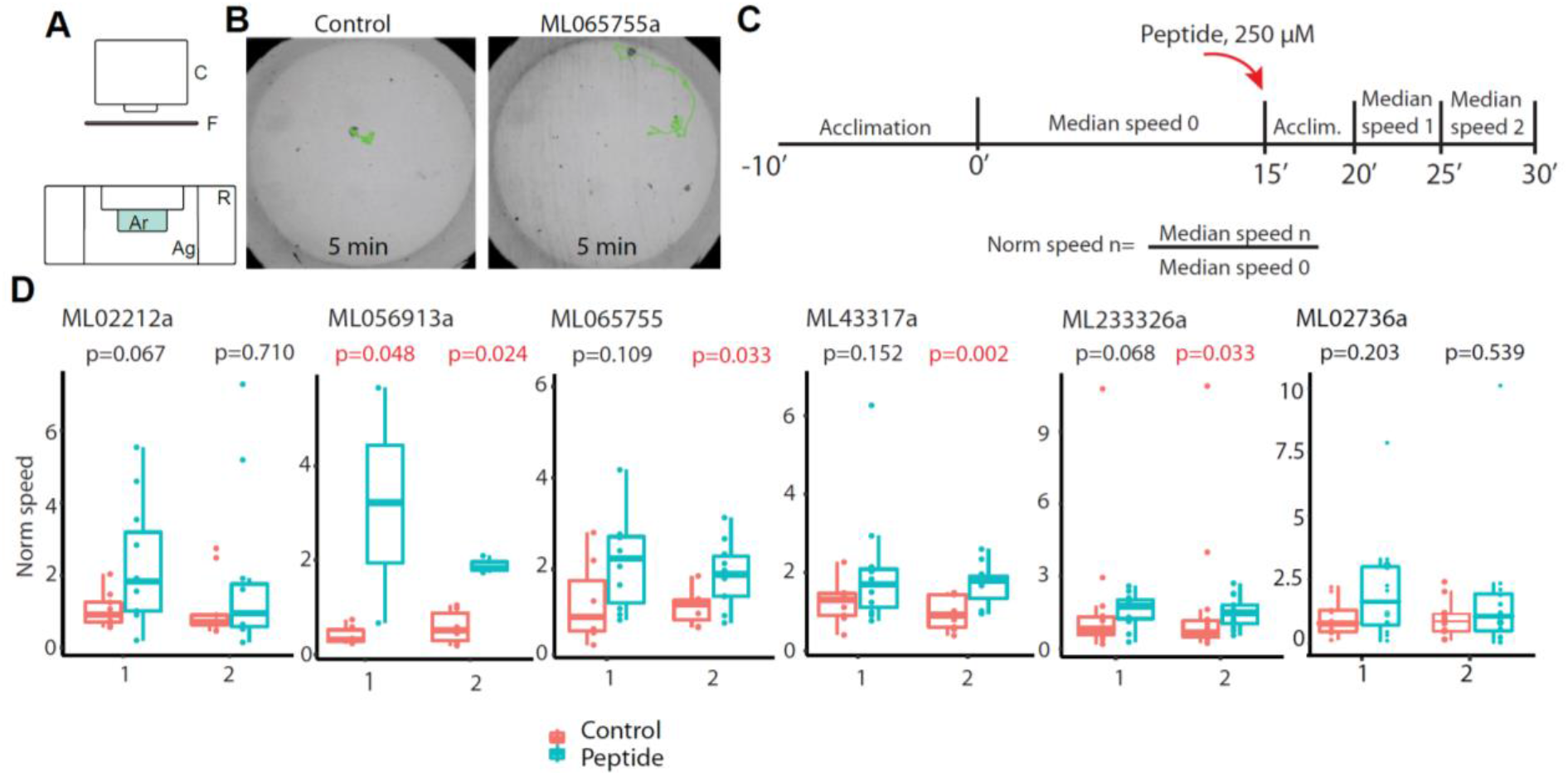
Study of neuropeptide effects on swimming in M. leidyi. (**A**) Behavioral set up. C-camera, F-infrared light filter, Ar – arena, Ag-agarose, R - PLA ring holding arena. (**B**) Examples of tracked 5 min recordings in control and treated samples; videos correspond to the interval 2 in (**C**). (**C**) Schematic of the behavioral assay. (**D**) Plots showing median speeds of the ctenophores after incubation with peptides or control. P values lower than 0.05 are shown in red (Mann-Whitney U).

### Neuropeptide expression patterns reveal the neuronal identity of cell types in the ctenophore single-cell transcriptome

Comparison of ISH and IHC patterns with scRNAseq^28^ profiles allowed us to annotate most of the previously uncharacterized *M. leidyi* metacells (**Fig. 4, Supplementary Table 4;** see also **Methods** section). Among the highest expressed transcripts in cells of the SNN (C33 metacell) are neuropeptides (**Fig. 4, Table 1**). Importantly, we revealed that C33 metacell correponds to the SNN neurons, cells with long branching neurites. Additionally, C33 expresses Secretagogin/Calbindin, Ferlins, Synaptogamin-7, Munc18 (STXBP1), Munc13 (UNC13B), Complexin and neurosecretory SNARE proteins Syntaxin 1, VAMP2^34^ and Snap-25 (**Supplementary Table 6**) which are important components of the bilaterian neurosecretory vesicle pathway^35-38^.

**Table 1.** Proteins of M. leidyi nerve net encoded by the C33 metacell transcriptome^2^. Overexpressed proteins^2^ are shown in bold, proteins with highest expression level in C33, but not reported as overexpressed in ^2^ are shown in regular font, proteins present in C33, but not overexpressed are shown in italic. Asterisks mark neuropeptides excluded from our ISH study.

| Function | Annotation | Protein ID |
| --- | --- | --- |
| Signal transmission | Neuropeptides | **ML199816a**, **ML02736a**, **ML02212a**, **ML056913a**, **ML017711a**, **ML215411a\***, **ML124215a\***, *ML01798a*, *ML014991a*, *ML003517a*, *ML206415*, *ML216920\**, *ML066512a\** |
|  | Ionotropic Glu receptor <sup>42,43</sup> | ML05909a |
| Electric excitability | Voltage gated Potassium channel <sup>39</sup> | **ML04675a** |
|  | Voltage gated Sodium channel <sup>40</sup> | *ML358826a*, *ML17059a* |
|  | Voltage gated Calcium channel <sup>41</sup> | *ML190424a* |
|  | Calcium-activated Potassium channel | **ML128229a**, ML04056a |
|  | Two-pore domain Potassium channel | **ML17562a** |
|  | Innexin | **ML036514a**, **ML32831a** |
| Secretion | Ferlin | **ML08307a**, **ML08309a** |
|  | Secretagogen/Calbindin | **ML03617a** |
|  | Synaptotagmin-7 | *ML16217a* |
|  | VAMP2 <sup>34</sup> | *ML02217a*, *ML214317a* |
|  | SNAP25 | *ML26853a* |
|  | Syntaxin-1 | *ML037014a* |
|  | Munc13 | *ML24335a* |
|  | Munc18 | *ML005115a* |
|  | Complexin | *ML08643a* |
| Neurite growth | Tubulins | **ML021133a**, **ML10376a**, **ML020045a** |
|  | Dematin | **ML334211a** |
|  | ABLIM3 | **ML435828a** |
|  | Contactin-5 | **ML087115a** |

**Figure 4.**
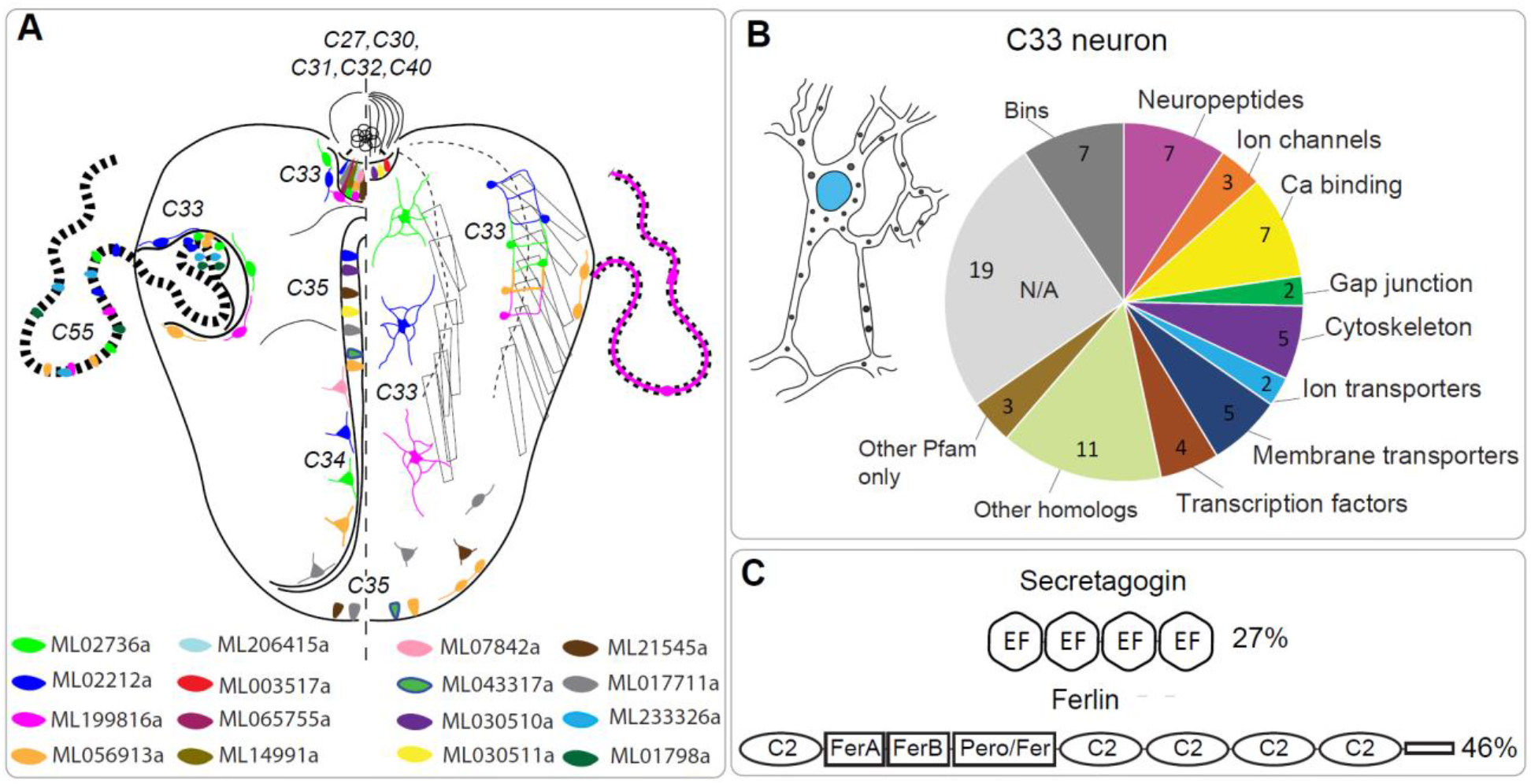
Neuropeptide expression patterns reveal identity of M. leidyi neurons. **(A)** Schematic representation of neuropeptide expression patterns revealed by ISH and IHC and corresponding metacells^2^. **(B)** Multipolar subepithelial neuron corresponding to the C33 metacell: schematic morphology and functional annotation of overexpressed genes^2^. **(C)** Domain structure of overexpressed calcium sensors involved in secretion is conserved; % shows the identity level to the human homolog.

We also found 35 homologs of voltage gated potassium channels^39^, two voltage gated sodium channels^40^ and the voltage gated calcium channel MleCav2^41^ as well as several homologs of iGluRs^42,43^ and other ion channels, Na/Ca exchanger, a sodium/potassium transporting ATPase and two innexins (**Supplementary Table 7**). To test for structural components defining neuron morphology^44-46^ we checked for the expression of structural proteins in the C33 metacell and found three enriched tubulin genes, the F-actin binding protein Dematin, the actin cytoskeleton organizer ABLIM3, Contactin-5 and Contactin protein-like 2 as well as genes described by “actin binding” and “tubulin binding” GO terms (**Supplementary Tables 8, 9**). Thus, the expression of multiple genes important for neuronal functions further confirms that C33 is a neuronal metacell.

### The ctenophore nerve net is composed of neurons with anastomosed neurites

Several of the characterised neuropeptide precursors and predicted structural neuronal components are expressed in the C33 metacell representing neurons of the SNN. To confirm the identity of the SNN neurons on an ultrastructural level, we performed volume electron microscopy of the SNN close to the AO and underlying the comb plates using SBF-SEM (**Fig. 5A-C, Supplementary videos 8-10**). The cell body of the reconstructed neuron of the SNN is located beneath and in contact with comb cells.

**Figure 5.**
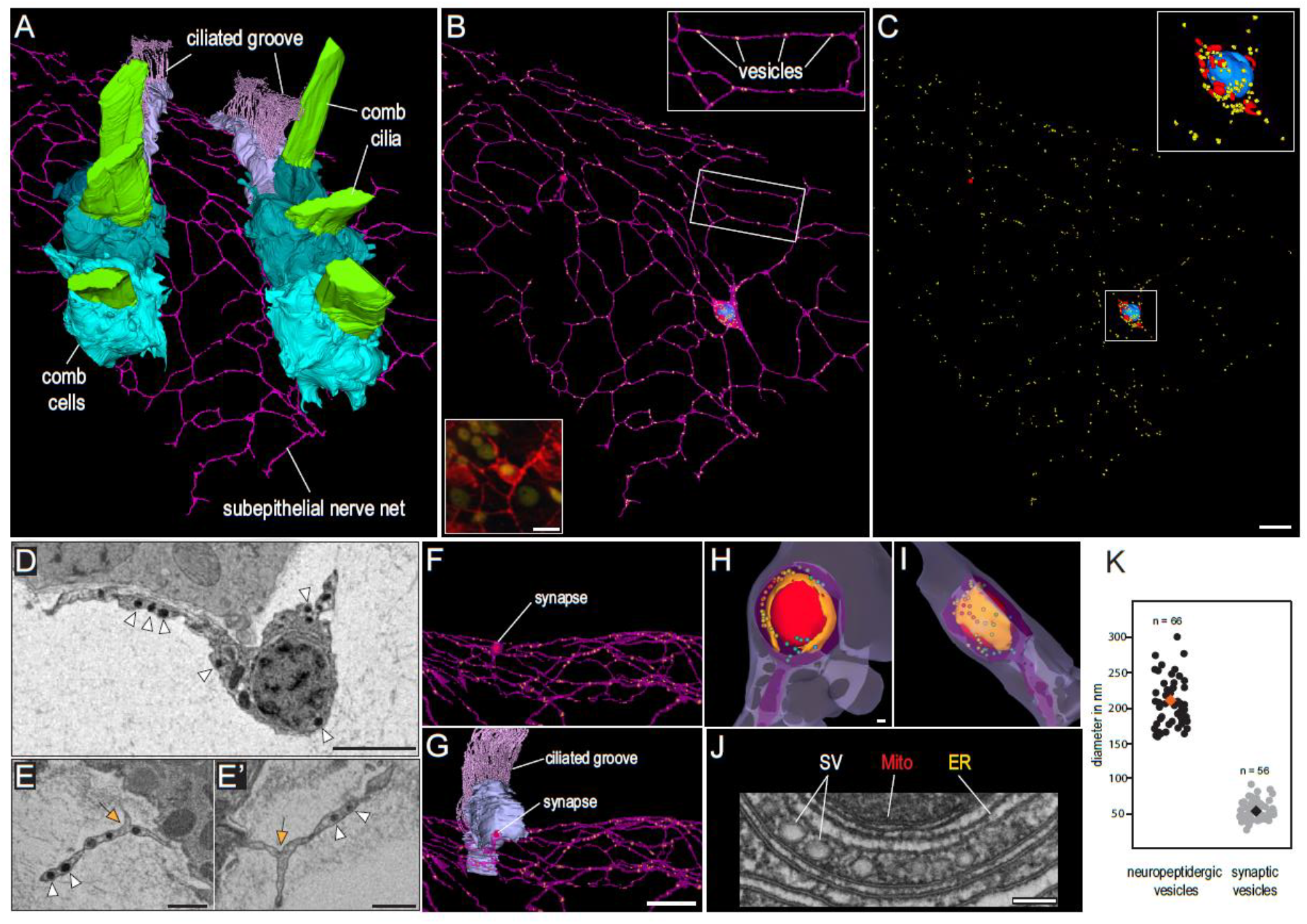
Ultrastructure of the ctenophore nerve net reveals morphology of multipolar neuron with multiple anastomoses between neurites. **(A)** 3D reconstruction of SBF-SEM data showing two comb rows underlaid by the epithelial nerve net. (**B)** Single neuron with several highly branching interconnected neurites (purple). Dense core vesicles probably containing neuropeptides (yellow), nucleus (blue). Bottom inset: neuron stained by IHC with anti-tubulin antibodies (red). (**C**) Localisation of dense-core vesicles (yellow), mitochondria (red) and nucleus (blue) in the same neuron. (**D, E**) SBF-SEM image of the cell body of the neuron with nucleus and two neurites. Note the multiple dense core vesicles (white arrowheads). (**E, E’**) Neurites with merged membranes (formation of anastomoses possibly by self-fusion; orange arrow) and multiple dense core vesicles (white arrowheads). (**F, G**) Synapse between the neuron and a ciliated groove cell (light blue). (**H, I**) 3D reconstruction of the synapse showing the triad: mitochondrion (red), endoplasmic reticulum (orange) and synaptic vesicles (rainbow colours). Loosely packed vesicles are represented in colder colors (lilac, blue, turquoise) while denser packed vesicles are represented in warmer colors (yellow, orange, red). (**J**) Electron micrograph of a synaptic triad (TEM). SV – translucent synaptic vesicles, ER – endoplasmic reticulum, Mito – mitochondrion. (**K**) Plot showing the size difference between the dense core neuropeptidergic vesicles and translucent synaptic vesicles. Scale bars: **A-C** = 4µm; **D-E’** = 1 µm; **F, G** = 4 µm; **H-J** = 100 nm

Five neurites branch off the cell body and extend in various directions giving the neuron a multipolar-morphology (**Fig. 5B-D**). Most intriguingly, many of these neurites are anastomosed to each other through continuous membrane (and not through synapses or gap junctions, **Fig. 5E and E**’). Thus, neurites of one cell form multiple polygons of different sizes and shapes (**Fig. 5B**). Nothing similar is known from neurons of other metazoans.

The presence of a presynaptic triad, specific to ctenophore synapses^10^, contacting a nearby ciliary groove cell further confirms the neural identity of this cell. Numerous electron dense vesicles resembling the neuropeptidergic vesicles of other metazoans in size and structure are present in the cell body and neurites. Interestingly, synaptic vesicles are much smaller than peptidergic vesicles (median diameters: 51 nm (N=56) and 202 nm (N=66), respectively; **Fig 5K**).

## Discussion

### Unique aspects of the ctenophore peptidergic nervous system

Our combined molecular, behavioral and structural approaches enabled us to reveal the uniqueness of ctenophore nervous system in previously unknown detail. The machine learning NeuroPID tool, predicting neuropeptide sequences based on biophysical, chemical and informational-statistical properties rather than sequence similarity^1^, allowed us to predict 129 novel ctenophore-specific neuropeptides precursors. We identified 16 neuropeptide precursors via ISH in the SNN and AO that are potentially post-processed into at least 42 mature peptides. For comparison, around 80 mature neuropeptides are described in the well-characterised *Drosophila*^47^. We also predicted the processing of neuropeptide precursors at positively charged cleavage sites (mostly dibasic). However, it is possible that cleavage occurs at other sites too as reported in cnidarians^48^.

Additionally, it is possible that the ctenophore neuropeptide repertoire is more extensive because we only focused on precursors of linear neuropeptides. Our analysis of the transcriptome of metacell C35 showed enrichment of some predicted disulphide-rich neuropeptides (ML306119a, ML056970a) in line with multiple putative disulfide-rich neuropeptides predicted in another ctenophore^15^. In addition, we considered only genes expressed in cydippid-stage *M. leidyi*, hence adult-specific as well as neuropeptides expressed during early development were not included in our ISH survey. Predicted neuropeptide precursors with relative expression levels higher than 50% in the annotated metacells^28^ were also excluded in this study. It is possible that some of them are regulatory peptides as well (for example, ML030512a is 57% identical to the ML030510a neuropeptide but expressed in the epithelial metacells). While there is no significant sequence similarity to metazoan neuropeptides, many ctenophore-specific neuropeptides are conserved among these animals; one peptide was shown to be restricted to the AO within the taxon^15^. The divergence of neuropeptides between animal groups resulting in many taxon-specific neuropeptides is a general pattern due to the fast evolution of the short protein sequences^19,25^.

Expression of identified neuropeptides is predominantly localized to components of the neuro-sensory system (**Fig. 4A**). The ctenophore AO functions as a hub of reception and signal integration, in the mouth and pharyngeal epithelium chemo- and tactile reception coordinate with muscle movement, and in the tentacle bulb stimuli from the tentacle integrate and spread via the SNN^10,11,49-51^. These organs therefore form a highly connected neuroendocrine system that regulate physiology and behaviour of these animals. It is worth mentioning that we find no peptidergic neurons in endoderm-derived funnel (“stomach”) and the gastral pouches. This is very different to other metazoans showing well-developed ecto- and endoderm-associated nervous components^8^.

To our best knowledge, our study is the first to characterize behavioral effects of ctenophore neuropeptides. We found four peptides that increase the median swimming velocity. ML065755a is expressed in the AO, known to regulate comb beating frequency^51^. ML43317a and ML233326a are expressed in sensory cells of mouth/pharynx and tentacles, respectively. They might therefore be involved in the regulation of hunting behavior as suggested from earlier experiments on tentacle stimulation and comb beating frequency^51^. ML056913a is expressed in multiple peptidergic cells of the AO and SNN probably affecting swimming through several unidentified mechanisms. ML02212a expressed in metacell C33 and the SNN underlying the combs showed only an insignificant tendency to affect median swimming velocity. However, it is possible that ML02212a and the other tested peptides, showing no significant effect, might regulate different physiological processes, different parameters of behavior or act only synergistically with other peptides and/or neuroreactive substances.

Volume EM 3D reconstruction provides first evidence for unique neurons (C33) in the SNN of cydippid-stage *M. leidyi* forming multiple self-anastomoses between neurites (**Fig.4B**). A similar morphology was suggested and depicted in hand drawings by the Haeckel-scholar Richard Hertwig over 100 years ago^11^. However, most observations on SNN structure were made in adult ctenophores in which the nerve net is formed by bundles of parallel neurites with unknown types of anastomosing connection between neurites^11,12^. Our immunostaining and SBF-SEM analysis uncovers that neurites belonging to one neuron in the *M. leidyi* cydippid-stage SNN form an extensive network with continuous membranes at the nodes of the net. While it is still unclear how these anastomoses are formed, self-fusion of neurites is probable scenario^52^. Loops created by neurites of the same neuron are known in mammalian brains however they occur through self-synapses^53^. Fusion of neurite membranes between different neurons has been described in cnidarians^54^ and bilaterians^55^ and is a common mechanism of giant fibre formation^56,57^. Self-fusion in contrast appears to be relatively rare representing a developmental or repair mechanism^52^. Besides an organizational process during development, neurite self-fusion in ctenophores may contribute to more differentiate or faster and amplified signal propagation. How different neurons within the same SNN are connected, either through synapses and/or gap junctions, remains to be investigated.

### Ctenophores and the evolution of animal neurons and nervous system(s)

It has been difficult to precisely define on a sole molecular basis what a “neuron” is ^58,59^. Therefore, a more integrative view combining genes, molecules, physiology, ultrastructure and morphology is needed to define a neuron. Combining all these aspects we were able to annotate metacell C33^28^ as neurons of the SNN. The presence of an extensive repertoire of components of secretory pathways in other *M. leidyi* metacells revealed by scRNAseq^28^ as well as in placozoans, in which almost all cell types are described as secretory^19^, give strong evidence for the evolution of neurons from peptidergic secretory (“gland”) cells^18,19^.

Despite their unique aspects, ctenophore peptidergic neurons show several conserved metazoan features. They co-express several neuropeptide precursors whose mRNA localizes to both neurites and cell bodies as reported for other metazoans^60,61^. Translucent vesicles were exclusively found in the synapse similar to other animals^60^ while dense core vesicles are evenly distributed to cell body and neurites which is in line with the observations in cnidarians and annelids supporting the hypothesis that the first nervous system(s) evolved as nets of peptidergic cells^62^.

If the ctenophore nervous system evolved independently from the nervous system of other metazoans cannot be said until the debate on their phylogenetic position has settled^63-65^. However, our analysis identified ctenophore neural features that are conserved due to functional constraints^66^ (e.g. the molecular secretory apparatus, a general neuronal ultrastructure, multiple voltage gated ion channels) as well as more adaptive components^67^ (e.g. neuropeptide repertoire, nervous system architecture and details of neuron morphology). Thus, regardless of the question if the ctenophore nervous system evolved independently, more detailed studies of the ctenophore nervous system are key to unravel the general principles underlying animal nervous system evolution and diversification.

## Material and Methods

### Prediction of putative neuropeptide precursors

*M. leidyi* protein models ^43,68^ were downloaded from Mnemiopsis Genome Project Portal (https://research.nhgri.nih.gov/mnemiopsis/download/download.cgi?dl=proteome). SignalP ^69^ online tool (http://www.cbs.dtu.dk/services/SignalP/) was used to identify sequences with signal peptides. NeuroPID online tool (http://neuropid.cs.huji.ac.il/) ^70^ was used to identify putative neuropeptide precursors. NeuroPID predictions rely on 4 machine learning models (such as support vector machines and ensemble decision tree classifiers); thus, prediction scorings by each of the four models are included in the output (**Supplementary Table 2**).

For the expression pattern analysis, quantitative *M. leidyi* single cell RNAseq data were used^2^. Expression level (represented as a Umifrac value) of each putative neuropeptide gene in each metacell was retrieved from http://www.wisdom.weizmann.ac.il/~/arnau/Single_cell_datasets/Mnemiopsis/. Because maximum expression levels vary drastically, Umifrac values for each neuropeptide gene were normalized against the maximum value for each gene so that the expression levels are represented by numbers from 0 to 1. A threshold of 0.05 was set and values ≤0.05 were replaced to 0. Putative neuropeptides expressed in metacells corresponding to known cell types at the level 0.5 – 1 were filtered out. Thus, the remaining putative neuropeptides are enriched in the unknown metacells.

### Search for neuropeptide homologs in other Ctenophora

The 30 predicted neuropeptide precursor sequences were used for Blast searches in the available Ctenophora transcriptomes and genomes. Search in the tBlastn mode was performed on the NCBI (https://blast.ncbi.nlm.nih.gov/Blast.cgi) in Ctenophora (excluding *Mnemiopsis*), TSA database resulting in the hits from *Hormiphora californensis* and *Beroe forskalii*, and on NeuroBase (https://neurobase.rc.ufl.edu/pleurobrachia/blast?view=blast) in *Pleurobrachia bachei* filtered gene models, *Beroe abyssicola, Beroe sp* pink, *Bolinopsis infundibulum, Bolinopsis ashleyi, Dryodora glandiformis, Euplokamis dunlapae, Pleurobrachia bachei-pileus, Pukia falcata* RNAseq databases. In both cases, e-value threshold was 10 and for the NeuroBase search “very short query” setting was used. *Mertensiidae sp, Coeloplana astericola* and *Vallicula multiformis* transcriptomes were excluded from the analysis due to reported contamination^71^ and search resulting in sequences identical to other ctenophore species (*B. infundibulum* for *Mertensiidae sp* and *M. leidyi* for *V. multiformis* and *C. astericola*). Retrieved hits were translated and only sequences possessing a signal peptide identified by SignalP were retained for further analysis. Translated sequences were aligned by MUSCLE algorithm implemented in Mega-X^72^ and visualized using JalView^73^. *Pleurobrachia* homologs retrieved from the genome and whole-body transcriptome were used for Blast search in the tissue specific transcriptomes from aboral organ, combs, tentacles, mouth, and stomach (NeuroBase, e-value threshold 10^−4^).

### Animal maintenance

Cydippid-stage animals used in the present study were obtained from reproductive cydippid individuals, 3-5 mm in oral-aboral length, and prior to the lobes development. Premature reproduction in *M. leidyi* was already documented over a century ago^74^, and studied in further detail by^75^. We have taken advantage of this capacity to establish a permanent and reliable culture setup based on cydippid reproduction. Seawater purified by mechanical (10, 5 and 1 µ), chemical (activated charcoal) filters followed by UV irradiation was used during all stages of the culture. First generation of reproductive cydippid individuals was isolated from a pool produced by adults (lobed) specimens kept at 16.5 °C, 27 ppt salinity and pH 7.9 – 8.1 and fed five times a week with 24 h *Artemia*, 48 hours enriched *Artemia*, and *Brachionus* (Rotifera). Adults were set to spawn following previously established protocols^75^. Subsequent generations were obtained from reproductive cydippids kept in 200 mL flat beakers (or crystalizing dish) 95 mm wide and 55 mm high, at a density of 5-10 individuals per beaker, 20-22 °C, 27 ppt and pH 7.9 – 8.1. Reproductive individuals were fed exclusively with living *Brachionus* (Rotifera), which were in turn fed with commercial concentrated algae (*Nannochloropsis* and *Tetraselmis*, ca. 50-60 billion cells per ml). Beakers containing reproductive cydippids were fed once a day, five times a week, with approximately 2000 rotifers per beaker and day, i.e. the equivalent of 200-400 rotifers per individual and day. Individuals were carefully transferred to new beakers containing fresh seawater once a week. In the described conditions, spawning takes place daily, and eggs hatch 22-26 hours after spawning. Therefore, 3-4 days old cydippids were harvested 4 days after water renovation.

### Tissue fixation

For fixation either aboral organs excised from adults (25 days old, ca. 7-8 mm in total length) or cydippids (3 – 4 days post fertilisation, starved for 5 h – overnight) were used. The tissue was fixed in ice cold 3.7% formaldehyde, 0.4% glutaraldehyde in artificial seawater (ASW) for 2 h on ice. The solution was exchanged to PTW buffer (1.8 mM KH2PO4, 10 mM Na2HPO4, 0.137 M NaCl, 2.7 mM KCl, 0.1% Tween-20, pH 7.4) and the cydippids were washed in PTW for 15 min trice at room temperature (RT). The cydippids were transferred to ice cold methanol through 50% methanol/50% PTW step and kept in −20 °C until use.

### In situ hybridisation

ISH protocol was developed based on published jellyfish *Clytia hemisphaerica* ^76^ and sea anemone *Nematostella vectensis* ^77^ protocols. Specific DIG-labelled RNA probes were produced using HiScribe Sp6 and T7 in vitro transcription kits (NEB). The sequences of the probes are in the **Supplementary Table 5**. Fixed *M. leidyi* cydippids were rehydrated trough 60% and 30% methanol in PTW steps, digested by 2.6 µg/ml Proteinase-K in PTW for 2 min and washed in 2 mg/ml glycine in PTW. After treatment with 1% triethanolamine and 0.6% acetic anhydride, the cydippids were refixed in 3.7% formaldehyde, 0.2% glutaraldehyde in PTW. Non-specific binding was blocked by preincubation in hybridisation solution (4 M urea, 50 µg/mL heparin, 1% SDS, 0.075 M sodium citrate, 0.75 M NaCl, pH7) containing 100 µg/mL salmon sperm DNA (Sigma) for 2 h at 60° C. DIG-labelled RNA probes were diluted in the hybridisation solution (0.5 ng/µl) supplemented with salmon sperm DNA and incubated with the cydippids for 36-60 h at 60° C. Non bound probe was washed by the concentration steps (100%, 75%, 50%, 25%, 0%) of hybridisation solution in 2x SSC (0.03 M sodium citrate, 0.3 M NaCl, pH 7) with 0.1% Tween20. Non-specifically bound probe was eliminated by RNAse T1 treatment (2 U/µl in 2x SSC, 40 min at 37 °C) followed by 0.05X SSC with 0.1% Tween20. The RNAse T1 treatment was not used for ML00218a, ML02736a, ML233326a, ML10665a probes. To visualise the probes, cydippids were incubated in blocking buffer (1% Roche Blocking reagent in 0.1 M maleic acid, 0.12 M NaCl, pH 7.5 buffer) 1 hr at room temperature and Roche anti-Dig antibodies coupled to alkaline phosphatase (1:3000 in blocking buffer) at 4 °C overnight. Non bound antibodies were washed with PTW (2 fast washes followed by 8×15min washes), alkaline phosphatase (AP) buffer (100 mM NaCl, 50 mM MgCl2, 0.1% Tween-20, 100 mM Tris, pH 9.5) first without MgCl2 and then with MgCl2. Cydippids were incubated with AP substrate solution (NBT/BCIP 1:50) at RT in dark until the staining developed; for most of the probes it took between 15 min to 4 h, for ML07842 and ML01798 staining was prolonged up to 60 h.

### Immunohistochemistry

Antibodies against four neuropeptides were raised in rabbit or mouse by Covalab (**Table 2**). All antibodies were affinity purified using antigen peptides as baits with the exception of ML017711a where a shorter peptide NH2-CDFR-NH2 was used.

**Table 2.**
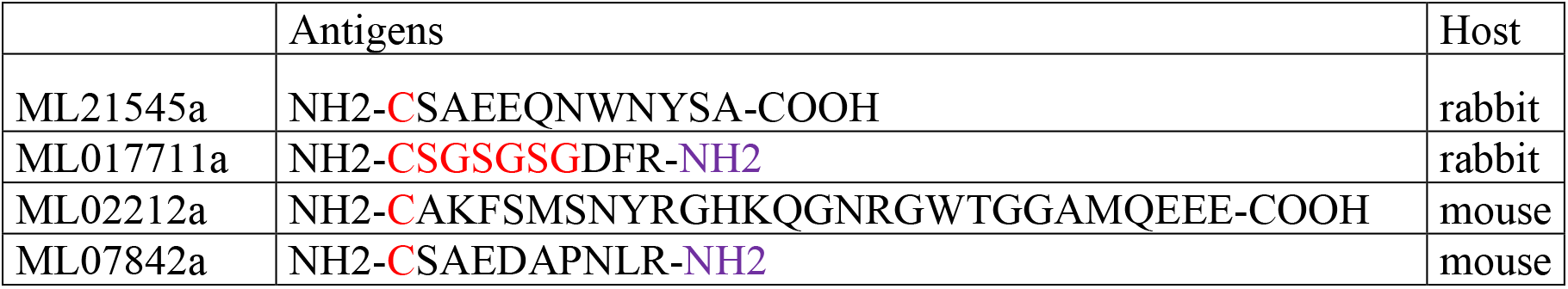
Sequences of the antigens used to produce anti-neuropeptide antibodies. Modifications to the native sequence are in red; amide groups are in purple.

*M. leidyi* cydippids were fixed and rehydrated from methanol as described for ISH. Rehydrated cydippids were permeabilized while washing five times for five minutes in PBTx (0.2% Triton X100 in PBS (137 mM NaCl, 2.68 mM KCl, 10.14 mM Na2HPO4, 1.76 mM KH2PO4, pH 7.4)), before incubating cydippids for 1 hour in Blocking solution (1% Bovine serum albumin in PBTx) at room temperature on a shaker. Neuropeptide antibodies raised in rabbit were combined with mouse E7 beta tubulin antibodies (DSHB). Neuropeptide antibodies raised in mouse were combined with rabbit recombinant anti-alpha tubulin antibody (abcam, ab190471). All primary antibodies were diluted 1:100 in blocking solution, spun down at 16873 rcf for 10 minutes and the supernatant was used for incubation with the samples overnight at 4 °C. The primary antibodies were washed off with five PBTx washes for 15 minutes. The secondary antibodies (anti-mouse/anti-rabbit coupled with Alexa Fluorophores 488 and 647 (abcam, ab150083, ab150117, ab150081, ab150119)) were diluted 1:250 in blocking solution, spun down at 16873 rcf for 10 minutes and only the supernatant was used for immunostaining. The samples were incubated over night at 4 °C, before washing three times in PBTx for 15 minutes and five times in PBS. The samples were mounted on an object glass with Vectashield antifade mounting medium containing DAPI (Vector Laboratories). All image stacks were taken with a 60x objective at either a Leica SP5 confocal microscope or with an Andor Dragonfly 505 confocal spinning disk system. Z-stacks of images were generated in ImageJ using standard deviation. The only adjustments done of the neuropeptide as well as tubulin signals was to increase the brightness if needed, making sure not to lose any information. The DAPI signal minimum was reduced slightly in some cases for better visibility of the two other channels in merged images. The 3D video clips were produced based on maximum intensity projections with Imaris. For best visualization of region of interest other regions were truncated with the clipping plane tools and brightness minima and maxima as well as gamma were adjusted.

### Behavioral test

For behavioral experiments, all cydippids were 3-4 days old, fed in the morning but had empty guts at the time of the experiment. 3 to 17 cydippids were used in each treatment or control. Nine neuropeptides (synthesised by GenScript, USA) were diluted with 27.5 ppt ASW or deionised water according to the solubility test results provided by the manufacturer. Custom made arena moulds were created using 3D printing with a depth of 4 mm and a width of 10 mm. Arenas were created by pouring agarose gel (0.9% in 27.5 ppt ASW) into small dishes and the arena mould was inserted into the liquid agarose gel. We used the custom-written software imMobilize (https://github.com/ChatzigeorgiouGroup/imMobilize) written on Python to record the videos with selected parameters. Arenas were placed under the camera set up (**Fig. 3A**) and were recorded in infrared light at 30 frames/second, 0.0078125 s exposure. Agarose arenas were washed with ASW after each use to avoid contamination between experiments; each cydippid was used only once and was discarded after use.

For recording, a single cydippid was added to the agarose arena and left for 10 min to acclimatise to the arena. Each arena was recorded for 15 min, then either a peptide or a control (ASW or deionised water, depending on the solvent for each peptide) were added, 5 min acclimation was allowed, and 2 x 5 min videos were recorded (**Fig. 3C**).

To track and analyse the videos, a software called Toxtrac^78^ was used with settings optimised for each *M. leidyi* cydippid. To use this programme, all video recordings were inverted by subtracting from a true white frame of equal size in imMobilize. Any background/interference was masked and not taken into account while tracking. Settings for the video tracking were as follows: ID algorithm: 2CTM sel; By shape (TEST); Max. displacement / Frame: 100. Videos with a detection of less than 95% were not used for further analysis.

To further analyse the ToxTrac outputs, we used a custom built software in Python called BehaviourAnalysis (https://github.com/ChatzigeorgiouGroup/BehaviourAnalysis) in order to obtain further parameters which account for the size of the arena and calibrated positions and distances of the animal. Median speed over 30 frames was calculated. Graphs were created using Rstudio software (2019 The R Foundation for Statistical Computing R version 3.6.1). Data was checked for normality using the Shapiro-Wilks test and datasets were found not to be normally distributed and non-parametric tests for Mann-Whitney U were performed on Rstudio.

### Electron microscopy and 3D reconstruction

#### Electron Microscopy sample preparation

For high pressure freezing and freeze substitution freshly hatched *M. leidyi* cydippids were pipetted into a glass dish with 20% BSA in natural seawater, immersed for several seconds and scooped out on a 0.2 mm aluminium planchette for high pressure freezing. The planchette was tapped onto filter paper to remove excess BSA, closed with the flat side of a 0.3 planchette dipped in hexadecane and immediately frozen in a Baltech10 High Pressure Freezer. 0.3 planchettes were removed directly after freezing and samples placed into precooled 100% Acetone and processed as follows in an RMS freeze substitution machine: 20 hrs at −90 °C in 0.1% UA + 1% osmium tetroxide, ramp up to −20 °C over 21 hrs, incubate 24 hrs a −20 °C, ramp up to 4 °C over 5 hrs, leave to warm up to room temperature for 1 hr. Samples were then washed in acetone and incubated in 1% tannic acid in acetone for 2 hrs, washed in acetone and incubated for 90 min in 1% osmium tetroxide, washed and infiltrated with 812 Epoxy resin as described above and embedded in resin blocks.

#### SBF-SEM

For Serial Block Face SEM imaging the cydippids were cut out from resin layer or block and glued onto SBFSEM stubs with conductive Expoxy resin (Chemtronics, Hoofddorp, Netherlands) and hardened for 8 hrs at 70 °C. The sample sides were trimmed with a trimming knife and the entire stub was sputter coated with a layer of gold (20-30 nm) to reduce charging. Settings for imaging the ROI in the Merlin Compact SEM (Zeiss, Cambridge, UK) with the Gatan 3View system and Gatan OnPoint BSD were as follows: Pixel size 5 nm, Dwell time 1µs, 20 nm Aperture, 1.8 kV acceleration voltage in high vacuum with Zeiss FocalCC (Focal Charge Compensation) aimed at the block face with nitrogen gas injection set to 100%, section thickness was 100 nm.

#### 3D Reconstruction & Analysis

For the reconstruction of *M. leidyi* epidermal tissue SBF-SEM sections were imported as z-stacks into the Fiji ^79^ plugin TrakEM2 ^80^ and automatically aligned using default parameters. Alignments were manually curated and adjusted if deemed unsatisfactory. Whole cells (neuron, comb cells, ciliated grove cells) and organelles were manually segmented, and 3D reconstructed by automatically merging traced features. Meshes were then smoothed in TrakEM2. Smoothing of the cell body in TrakEM2 sacrifices fine structures associated with cellular projections and therefore cilia of ciliated grove cells and vesicles of the neuron were not smoothed. All other organelles (nucleus, mitochondria) were subjected to the same smoothing parameters.

For the reconstruction of a ctenophore synapse, digital image stacks of high-resolution TEM sections of single synapses were imported into AMIRA (FEI Visualization Sciences Group) and aligned and vesicles, ER, mitochondria and the plasma membrane manually segmented by tracing these structures along the z-axis. Surface models were rendered from the ER, mitochondria and plasma membrane using the “generated surface”-module and subsequently smoothed by reducing the surface vertices and repeated use of the “smooth surface”-module. A color-coded density visualization of the vesicles was generated by calculating a “density-map” as basis for the “Point Cloud Density”-module. This module measures the minimal distance of the vesicular edge (the vesicle membrane) to the edges of the neighboring vesicles. The minimal distances of one vesicle to the other, neighboring, vesicles were then visualized with a “Point Cloud View”-module and color-coded with the physical look up table. Loosely packed vesicles are represented in colder colors (lilac, blue, turquoise) while denser packed vesicles are represented in warmer colors (yellow, orange, red). Vesicle diameters were measured in ImageJ ^81^ using the line tool and the “ROI” manager to show the lines permanently to prevent multiple measurements of a single vesicle. Mean vesicle diameters were calculated using unsmoothed, unprocessed meshes from 56 measurements from synaptic vesicles and 66 measurements from electron-dense vesicles.

### Annotation of peptidergic metacells

ISH and IHC patterns of the neuropeptides were manually compared with scRNAseq profiles^2^. Both ML065755a and ML206415a neuropeptides have nearly identical expression patterns restricted to AO sensory cells (**Supplementary Fig. 7I-L**) and correspond to C29 and C30 metacells. In cydippids, ML14991a is expressed in several types of AO sensory cells with distinguishable cilia (**Fig. 2C, Supplementary Fig. 7M-O**), which in adult AO appear as five groups of cells (**Supplementary Fig. 7P**) corresponding to C29, C30, C32, C36, C39 metacells. ML003517a (**Supplementary Fig. 7R-U**) is expressed in the C32 metacell, confirming its AO localisation, and the AO rim thus corresponding to C27. The cells of the pharingeal neurons expressing ML07842a (**Supplementary Fig. 9A, B**) correspond to C34 metacell, where this gene has the highest expression. The ML07842a-positive aboral organ cells identified by IHC (**Supplementary Fig. 9D-G**) most probably represent C40 since the expression level in this metacell is much lower than in C34 supported by the fact that we could not detect it by ISH in cydippids. In the adult AO, ML07842a is expressed in three groups of cells probably corresponding to the C40 along with C32 and C36 where it has minor expression (**Supplementary Fig. 9E - G**). The highest expression of ML43317a, ML030510a and ML030511a was detected in C35 metacell (**Supplementary Fig. 9H-J, M, N, 10C)**. Expression of ML030510a and ML030511a in the AO rim corresponds to the C27 metacell identified earlier (**Supplementary Fig. 9L, 10B**. ML233326a and ML01798a expression was observed in tentacles and tentacle bulb and it is enriched in the C55 metacell (**Supplementary Fig. 13**). The highest expression level of ML199816, ML02212, ML02736 in the C33 metacell evidences that it corresponds to the epthelial nerve net (**Fig. 2, Supplementary Fig. 6, 7**). At lower levels these genes are expressed in C34 and C35 that can be cofirmed by ISH and IHC (**Supplementary Fig. 8A, B, F**). The expression in the sensory AO cells of ML02212a can be attributed to the C30 metacell (**Supplementary Fig. 6I, J**) while the aboral expression of ML199816a can be explaied by C28, C32, C36 and C40 metacells (**Supplementary Fig. 8C-E**); the latter also expresses ML02736a. Neurites have been reported to penetrate the aboral organ floor^82^ therefore the aboral expression of ML199816a, ML02212a and ML02736a corresponds to the C33 metacell. ML056913a is mostly expressed in C34 and C35, which corresponds to the multiple cells around the gut and the mouth (**Supplementary Fig. 8K-O**), and in the C33 at the lower level corresponding to the cells with neurites on the body surface, especially next to the comb plates (**Supplementary Fig. 6P-S**). Additionally, ML056913 is found in C55 tentacle metacell (**Supplementary Fig. 8J**), in AO cells confirming that C28, C29, C32, C36 are indeed AO metacells (**Supplementary Fig. 8G-I**). ML21545a is expressed in C35 (**Supplementary Fig. 11D-E**) and C31 indicating that C31 is an AO metacell (**Fig. 5B-C**). ML17711a is expressed in C28, C29, C31, C32, C36, C40 identified as AO metacells earlier (**Supplementary Fig. 11J-L**) as well as in C33, C34 and C35 (**Supplementary Fig. 11H**).

### Functional annotation of the C33 metacell transcriptome

*M. leidyi* protein models were annotated using BlastP search in human, drosophila and yeast Uniprot databases in the earlier study ^2^. To confirm and improve the automated annotation of genes expressed in the C33 metacell (specifically enriched or relevant to nervous system) we ran reciprocal BlastP search in the *M. leidyi* protein model database (https://research.nhgri.nih.gov/mnemiopsis/sequenceserver/) using the human homologs as queries. We also aligned the human and *M. leidyi* sequences (**Supplementary Fig. 16**) using Clustal Omega (https://www.ebi.ac.uk/Tools/msa/clustalo/) and examined their protein family and domain structure with InterProScan (https://www.ebi.ac.uk/interpro/). Ferlin ML08309a and ML08307a protein models appeared truncated therefore we searched for full length sequences in NCBI *M. leidyi* TSA database using tBLASTn.

To identify putative proteins involved in ctenophore secretion, neurite growth and electric transmission, we employed Panther classification system (http://pantherdb.org/) and GO terms SNARE binding (GO:0000149), actin binding (GO:0003779), tubulin binding (GO:0015631), ion channel activity (GO:0005216) using Uniprot IDs of the human homologs as an input. To retrieve the human Uniprot IDs (they were not included in the earlier annotation ^2^), we ran BlastP in the reviewed human Uniprot database with e-value cutoff 10^−5^ with the *M. leidyi* protein models as it was done in the study^2^. For actin and tubulin binding homologs, we also checked if their expression has been detected in human nervous system using the Bgee database (https://bgee.org/). Here we considered genes both reported as overexpressed and whose expression was just detectable (above threshold of zero) (**Table 2, Supplementary Tables 6-9**).

## Acknowledgments

We thank Jeffrey Colgren for valuable comments on the paper, Endy Spriet and Hege Avsnes Dale (Molecular Imaging Center of University of Bergen) for assistance with fluorescent microscopy. Additionally, we acknowledge Kevin Pang for providing animals to start our culture. This work was supported by the Sars Centre core budget.

## Author contributions

MS, MC, MK and PB designed the study; MS, ELN, JJSA, YM, BG, BN, DD, MK, and PB performed experiments; MS, ELN, YM, BN and PB analyzed data; MS and PB wrote the paper and all authors reviewed, commented on, and edited the manuscript.

