## Supplementary Materials for "The unique neuronal structure and neuropeptide repertoire in the ctenophore *Mnemiopsis leidyi* shed light on the evolution of animal nervous systems"

**Supplementary figure 1.** Workflow for the identification of putative neuropeptide precursors of the ctenophore *M. leidyi*.
**Supplementary figure 2.** ML00218a gene specifically overexpressed in C42.
**Supplementary figure 3.** Processing of *M. leidyi* neuropeptide precursors.
**Supplementary figure 4.** Sequence alignments of diverse ctenophore neuropeptide precursors.
**Supplementary figure 5.** Developmental profiles of putative *M. leidyi* neuropeptides.
**Supplementary figure 6.** Neuropeptides expressed in the subepithelial nerve net.
**Supplementary figure 7.** Neuropeptides expressed in the subepithelial nerve net, in multiple neural cell types and restricted to the aboral organ.
**Supplementary figure 8.** ML02212a, ML199816a and ML056913a expression.
**Supplementary figure 9.** Neuropeptides expressed in the pharynx and mouth area.
**Supplementary figure 10.** ML030511a is expressed in the AO rim and sensory cells in the pharynx.
**Supplementary figure 11.** Neuropeptides expressed in multiple cell types.
**Supplementary figure 12.** ISH with neuropeptide precursors expressed in the AO cells.
**Supplementary figure 13.** ISH of neuropeptide precursor genes expressed in the tentacles.
**Supplementary figure 14.** ML10665a is expressed in the meridional canals and pharynx.
**Supplementary figure 15.** Behavioural experiments with *M. leidyi.*
**Supplementary figure 16.** Alignments of *M. leidyi* and *H. sapiens* neuronal proteins shown in Table 1.

**Supplementary video 1.** ML02212a localization in the aboral organ of *M. leidyi*.
**Supplementary video 2.** ML07842a localization around the pharynx of *M. leidyi*.
**Supplementary video 3.** ML07842a localization in the aboral organ of *M. leidyi*.
**Supplementary video 4.** ML07842a localization in the aboral organ of an adult *M. leidyi*.
**Supplementary video 5.** ML21545a localization around the pharynx of *M. leidyi*.
**Supplementary video 6.** ML17711a localization around the pharynx of *M.leidyi*.
**Supplementary video 7.** ML17711a localization in the aboral organ of *M. leidyi*.
**Supplementary video 8.** 3D reconstruction of a part of *M. leidyi* epidermis.
**Supplementary video 9.** 3D reconstruction of a part of *M. leidyi* epidermis.
**Supplementary video 10.** 3D reconstruction of a part of *M. leidyi* epidermis.
**Supplementary video 11.** 3D reconstruction of a *M. leidyi* synapse.

**Supplementary table 1.** Neuropeptide precursors predicted by NeuroPID.
**Supplementary table 2.** NeuroPID scoring for the linear neuropeptide precursors.
**Supplementary table 3.** Neuropeptide homologs uncovered in other ctenophore species.
**Supplementary table 4.** Summary of neuropeptide expression patterns and metacell annotation.
**Supplementary table 5.** ISH probes.
**Supplementary table 6.** GO terms identified in the C33 transcriptome: SNARE binding.
**Supplementary table 7.** GO terms identified in the C33 transcriptome: Ion channel activity.
**Supplementary table 8.** GO terms identified in the C33 transcriptome: Actin binding.
**Supplementary table 9.** GO terms identified in the C33 transcriptome: Tubulin binding.

#### Supplementary figures

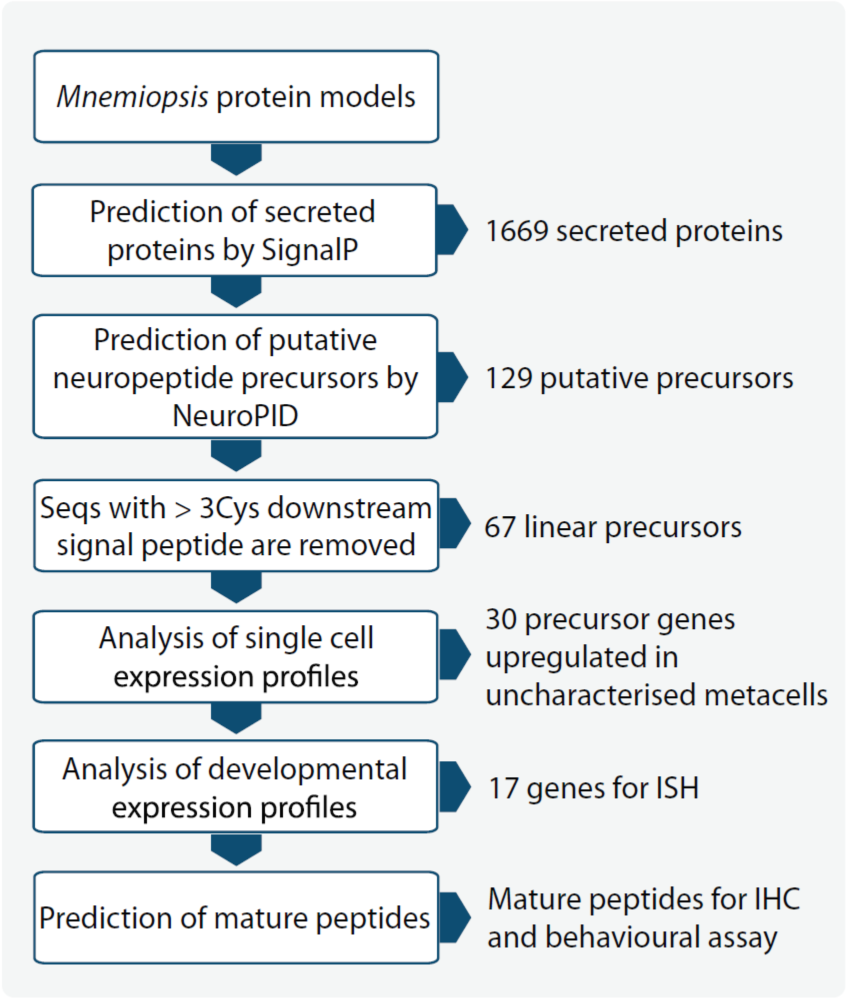


***Supplementary figure 1. Workflow for the identification of putative neuropeptide precursors of the ctenophore M. leidyi*.**

ML00218a

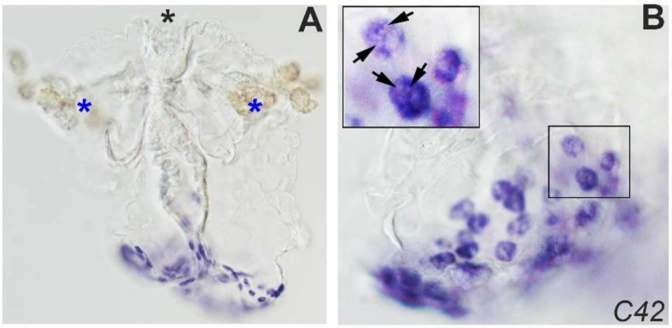


***Supplementary figure 2. ML00218a gene specifically overexpressed in C42 indicates that this metacell represents gland cells around the mouth and in the pharynx. C42 was not annotated in the original study. (A)*** *Whole cydippid.* ***(B)*** *Close up to the mouth area. Black arrows highlight unstained secretory granules.*

**Repetitive sequences**

>ML043317a

*MKTILLISCLLSAVYS*RVLRLNEEDWSRNGAMEDSEDWSRNGAMRDSEDWNRNGGMRDSEDWSRNGRMEDSEDWSRNGAMEDSEDWSRNGAMEDSEEWSRNGGLENSEDWSRNGAMRDSEDWSRNGAIKDSEDWSRNGAMRDSEDWSRNGAMKDSEDWSRNGMEDSEDWSRNGA

>ML07842a

*MMKLTVVCLLFASLLVRAEC*GVVERHEEQLFDAEDEQPNFRAGLENKELEKRSPNFRFGSKRSAEDAPNLRGKRSNEDTPNFRGTKRSAEDSPNFRGAKRSAVEAPNFRGAQKSEEDAPHFRGAKRFAEDAPHLRGVKRSTVDAPNFRGAERLEEDVPNLRGAERSEEGSPNFRGAERAVEDAPQFRG

>ML21545a

*MSKFLFLTLLVGAARA*ASFASDSDNTLADSDECRTGYNMRYDLECRRRRAARAAARLQLQEDGETEGGEELTKRSAEEQNWNYSARRDRGKRSNNEEQDFSGYRRGSGQKRSAEEDDYDIYKRGESEEQDYSGYRGRGRGE

>ML017711a

*MKMFILIGLLITLVLNYVSA*GDLARRSLEEENESGINEGAAEEDDESFRGLRESEDESRGKREVADEELFRGKRDFREKRDFRGKRDFRGKREFRGKREFRGKRDFRGMKSSEEELVKRDLSRGQ

>ML016347a

*MICLNFRVTAVVLFVILLAGFCPA*APIYSTQENDAFLETGEEKGELDIEAPKTPKLRYFYQKRSEEGKDEDPKLQYFYQIRSLEDKEKKPKLRYFYQKREENKDTKPKLRHFYQKREENKDTKPKLRYFFQKREENKDTKPKLRYFYQEREDNKDTKPKLRYFFQKREENKDTKPKLRNFYRKRGEN

>ML11723a (adult specific)

*MKYFMILAVLILTVGS*SPFEKRDAENAELASEKRSEEADPGEEYVLAKRNVDQDAVSLEKRQKYGYKRGE

>ML06405a (adult specific)

*MRKTLCIQLLVLLAIQRFTDG*KAFEESENEVLSLEKRAEDLDDEDFFDKRETEDRENEVEIDLD

**Non-repetitive sequences**

>ML02212a

*MKLFLFVLLGLVALISC*ETVESEVDSELSESEDSNAMRVKRAKFSMSNYRGHKQGNRGWTGGAMQEEE

>ML02736a

*MKCFVVLFALLALSQSAS*LNSLESVEDVIMADNDNVELEEGALNAEEEARVYKGYNGGNRVWYG

>ML199816a

*MKLFLFLTASLLVLATVTVQTEA*REIVAEVAESEAVESAASEEDTFVYRKEEDSAFLFAD

>ML056913a
*MFRLTSILLVLVLAIVTFTRT*IEDSEEIGHGLRMDKNEEIGHGLTLDKRSADSVDTEAMADEEELVGHGIKGSHAWRK

>ML01798a

*MKLLLLTLAVLLACLTA*VPVRQEEVPQEEIRLERSAESSGGETARVEKRAAIDTGSDYPGFEGGKRRWYG

>ML233326a

*MKTTFLVLTLMIICCNYVQS*VPMSLEEDLSDEEHRGLQKRAGTKFNKADYKSVGEGTRKWFG

>ML065755a

*MKLAAGIFLVLACLTVVMIPGEA*ASLGSLDAANSDSNAVLYGADHEILESSLINRKEEAEGALYG

>ML215411a

*MKQRITILLTILGVIVLAQS*KSILVESEDVLLGDNAELEVDSNSVDLDEELIGSDIKLVPGSGGNPWGRK

>ML124215a

*MSRMLFAVTLVIFSVLAITSG*ASLSEEDEELLTGLEDQMENSEETMMYAASDDADNNEAEMAKRGGKARYRRW

**Supplementary figure 3. Processing of *M. leidyi* neuropeptide precursors**. *Signal peptides predicted by SignalP tool are shown in yellow and italics, cleavage sites are in purple, Gly residues converted into C-terminal amides are in blue, mature peptides are underlined, negatively charged residues are in red, positively charged residues are in blue. Mature sequences used for antibody production and behavioural tests are in grey. To predict mature peptides, we were guided by the following assumptions: 1) cleavage occurs at dibasic or monobasic motives; 2) mature peptides are the most conserved parts of the precursors (***Fig 1D***,* ***Suppl fig 3****); 3) propeptides tend to be negatively charged); 4) C-terminal Gly is converted into an amide group*

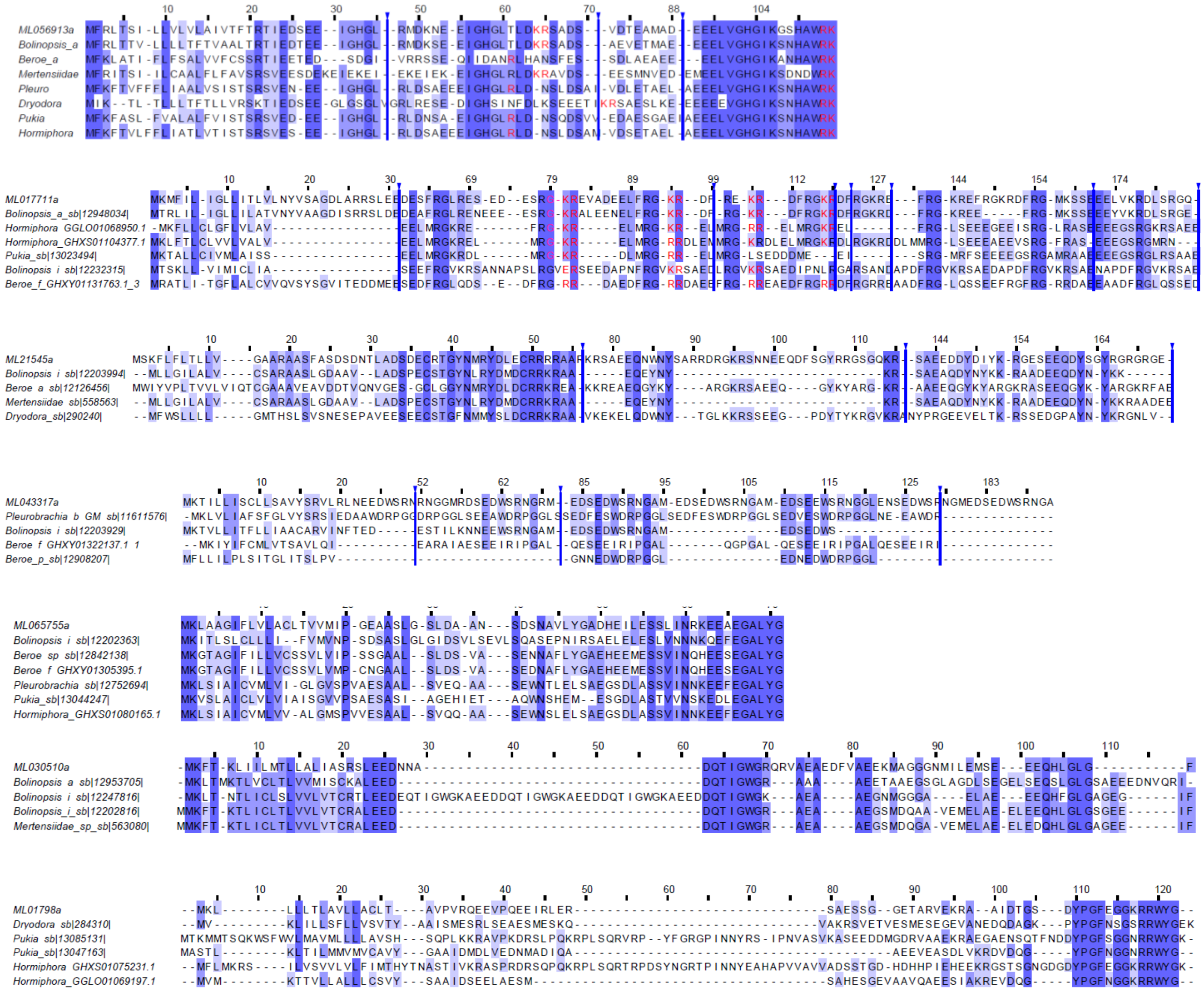
ML056913a
Bolinopsis_a
Beroe_a
Mertensiidae
Pleuro
Dryodora
Pukia
Hormiphora
ML017711a
Bolinopsis_a_sb|12948034|
Hormiphora_GGLO01068950.1
Hormiphora_GHXS01104377.1
Pukia_sb|13023494|
Bolinopsis_i_sb|12232315|
Beroe_f_GHXY01131763.1_3
ML21545a
Bolinopsis_i_sb|12203994|
Beroe_a_sb|12126456|
Mertensiidae_sb|558563|
Dryodora_sb|290240|
ML043317a
Pleurobrachia_b_GM_sb|11611576|
Bolinopsis_i_sb|12203929|
Beroe_f_GHXY01322137.1_1
Beroe_p_sb|12908207|
ML065755a
Bolinopsis_i_sb|12202363|
Beroe_sp_sb|12842138|
Beroe_f_GHXY01305395.1
Pleurobrachia_sb|12752694|
Pukia_sb|13044247|
Hormiphora_GHXS01080165.1
ML030510a
Bolinopsis_a_sb|12953705|
Bolinopsis_i_sb|12247816|
Bolinopsis_i_sb|12202816|
Mertensiidae_sp_sb|563080|
ML01798a
Dryodora_sb|284310|
Pukia_sb|13085131|
Pukia_sb|13047163|
Hormiphora_GHXS01075231.1
Hormiphora_GGLO01069197.1

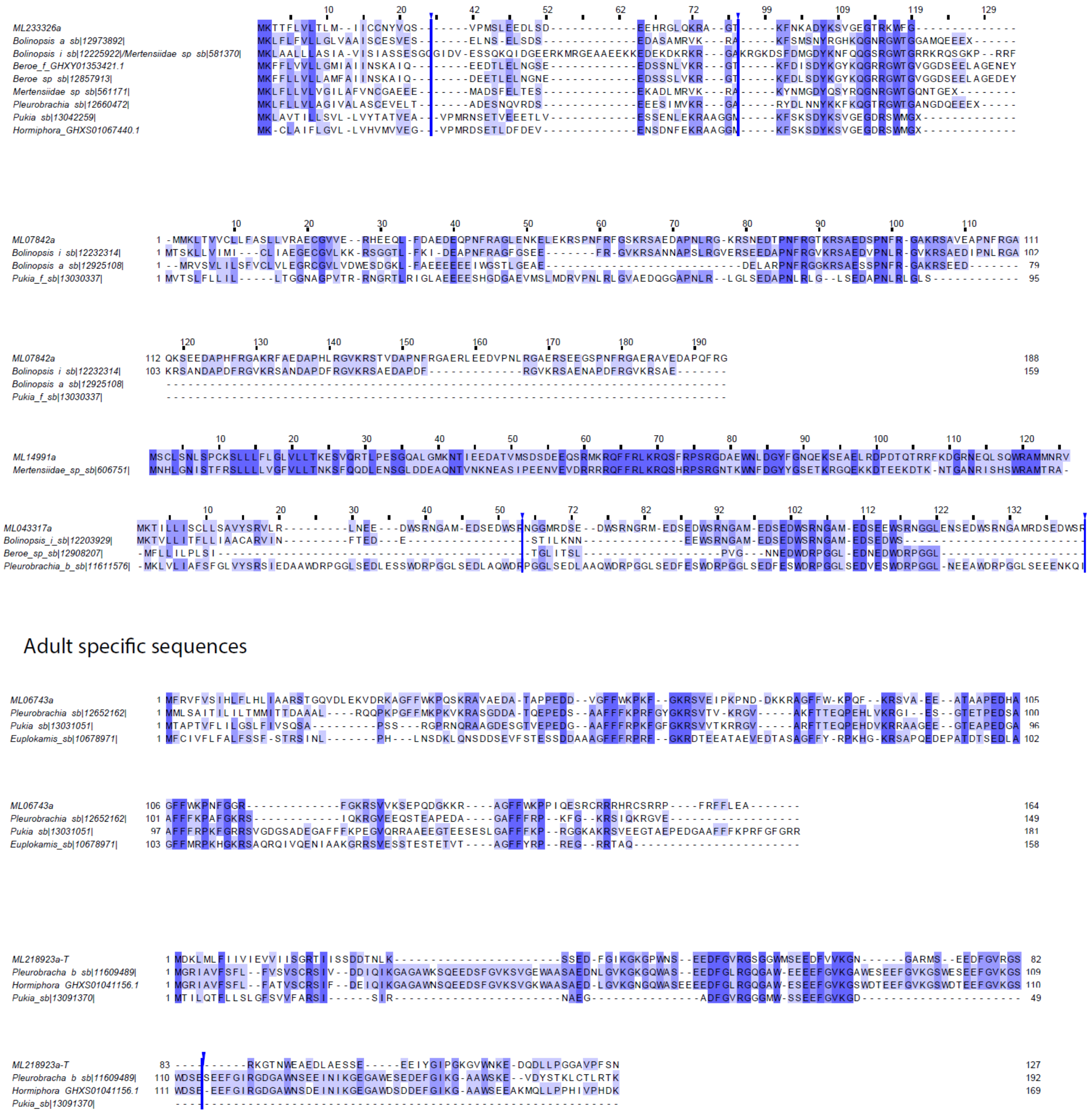


***Supplementary figure 4. Sequence alignments of diverse ctenophore neuropeptide precursors.***

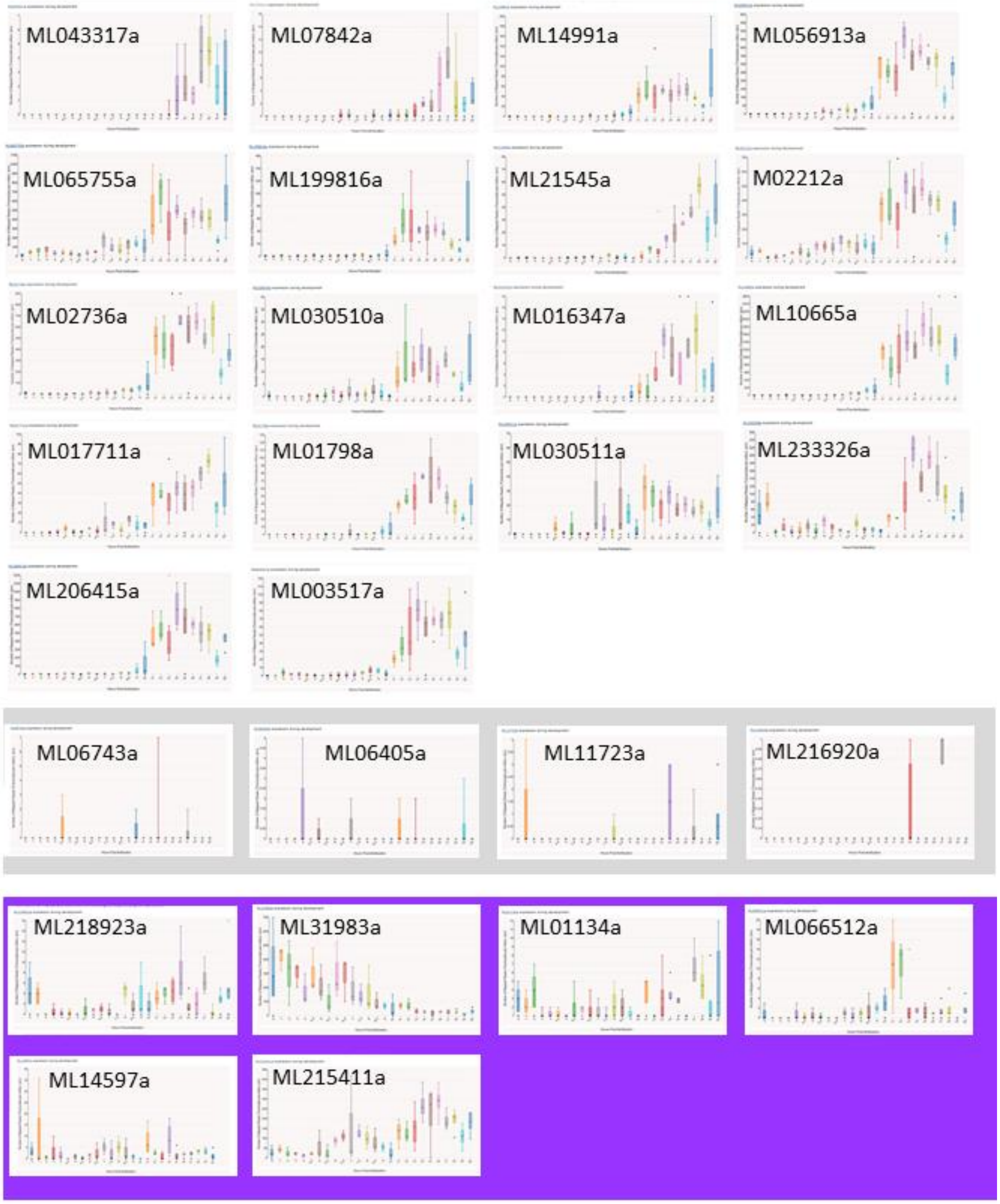


***Supplementary figure 5. Developmental profiles of putative M. leidyi neuropeptides*** *(https://research.nhgri.nih.gov/mnemiopsis/Mlei_expression_timecourse/). Genes that are not expressed in cydippids are labelled in grey, genes that have expression dynamics which do not correlate with the estimated nervous system development dynamics are labelled in purple.*

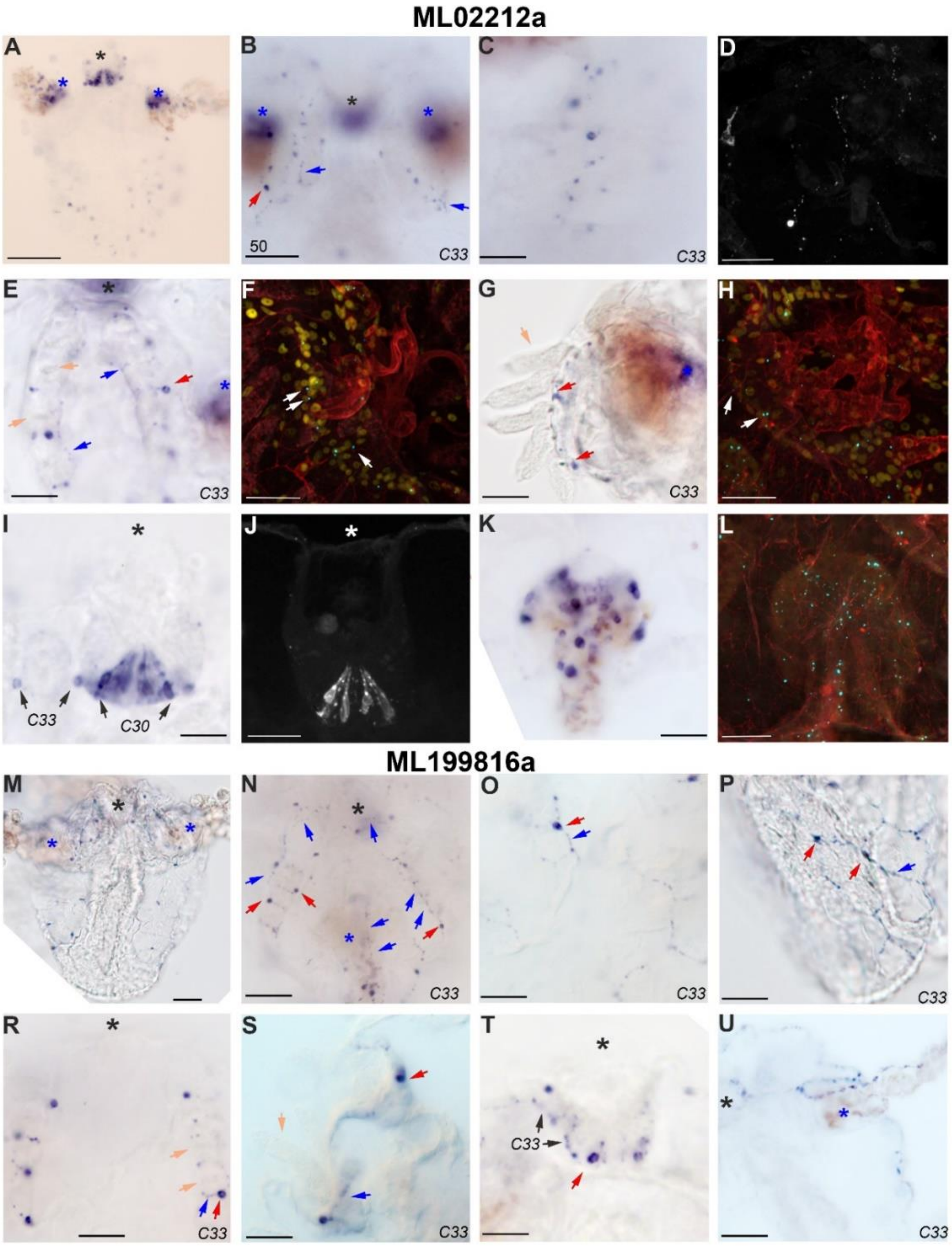


***Supplementary figure 6. Neuropeptides expressed in the subepithelial nerve net.*** *Black star – aboral organ, blue star – tentacle bulb, red arrow – neuron body, blue arrow – neurite, orange arrow – comb plate.* ***(A-L)*** *ML02212a is expressed in the subepithelial nerve net (**B-H, K-L**) and in the AO (**I, J**); (**E-H**) ML02212a expression around the combs.* ***(M-U)*** *ML199816a is expressed in the subepithelial nerve net (**N-T**) and in the mesogleal neurons of tentacles; (**T**) shows the AO and (**R, S**) show the area around the combs.* ***A-E, G, I, K, M-U*** *– staining by ISH**;** **D, F, H, J, L** – staining by IHC, the mature peptide is cyan, tubulin is red, DAPI is yellow.* ***(A, M)*** *Whole cydippid. Scale bars: 50 µm for (**A, M, B, U**), 20 µm for (**C-L**, **N-T**).*

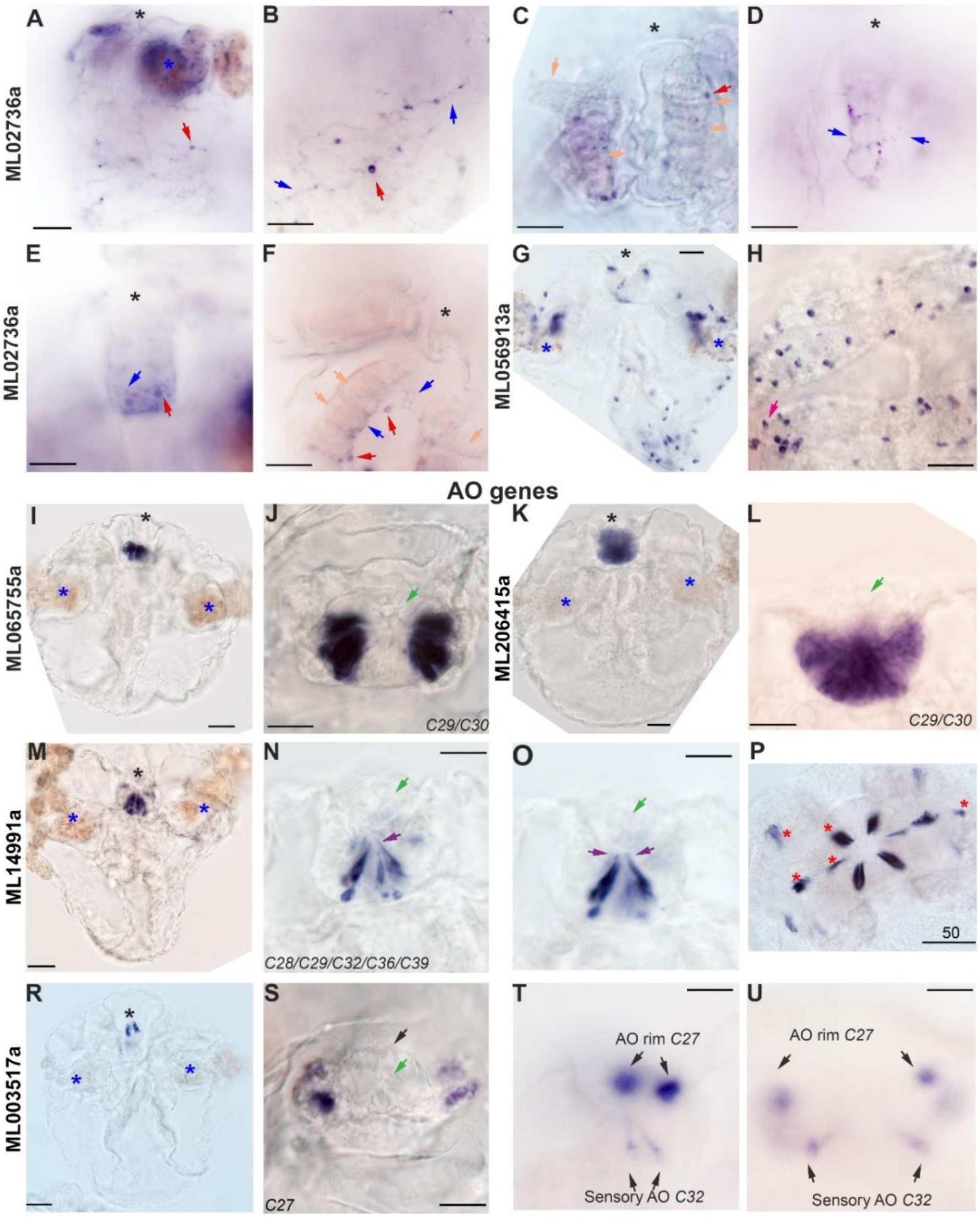


***Supplementary figure 7. Neuropeptides expressed in the subepithelial nerve net (ML02736a, A-F), in multiple neural cell types (ML056913, ML206415a, ML14991a, ML003517a, G-H) and restricted to the aboral organ (ML065755, I-P).*** *Star and arrow labels are the same as at the Fig 3, green arrows – statolith, magenta arrow – sensory cells on the lips, brown arrow - dome. (**A, G, I, K, M, O**) whole cydippid; (**B**) epithelial nerve net, (**C, D, F**) combs, (**E, J, L, P**) aboral organ, (**H**) lips. Scale bars: 50 µm for (**A, G, I, K, M, P, R**), 20 µm for (**B-F, H, J, L, N, O, S-U**).*

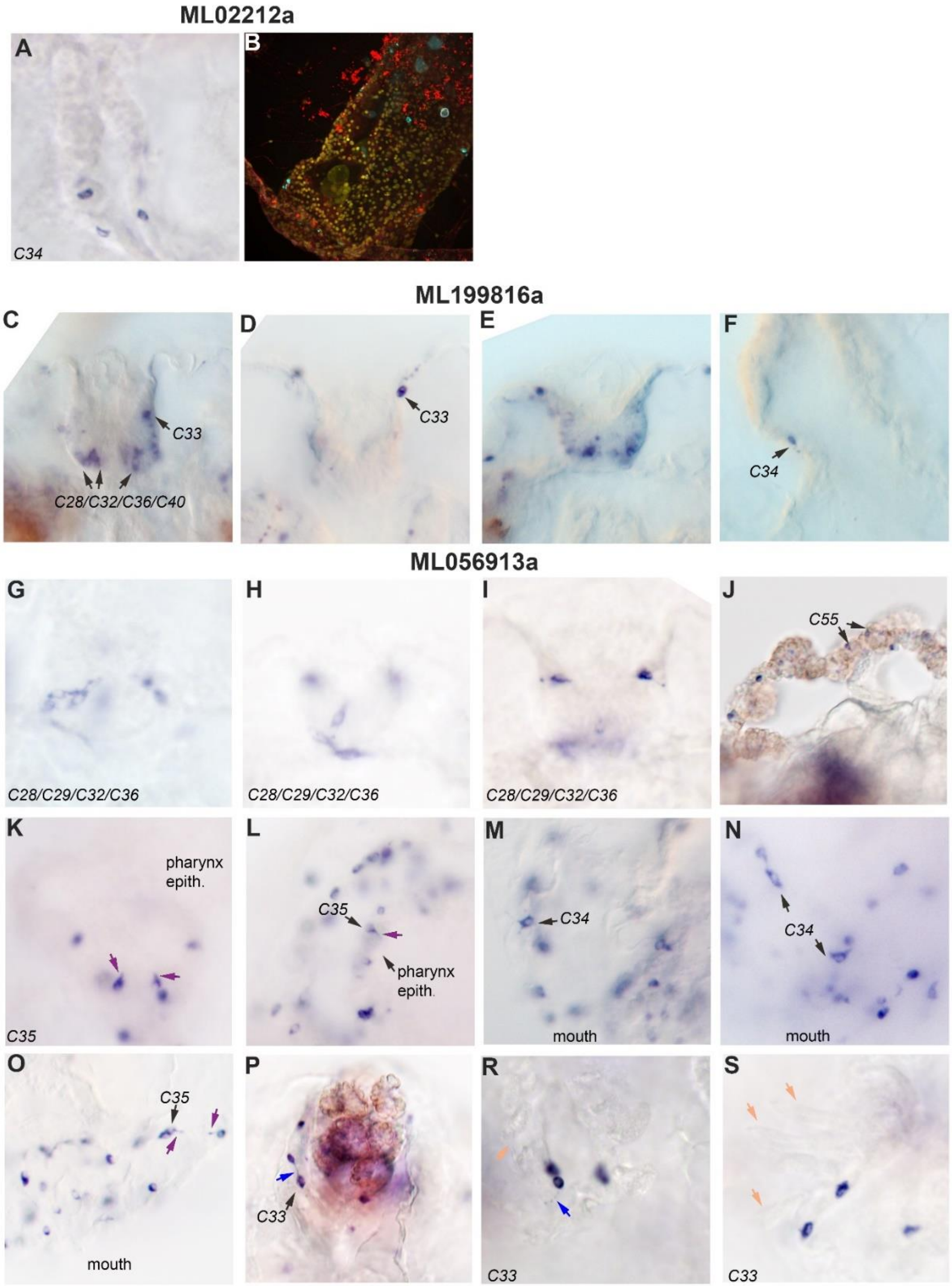


***Supplementary figure 8. ML02212a, ML199816a and ML056913a expression.*** *Blue arrow – neurite, orange arrow – comb plate, black arrows – corresponding metacells.(**A-B**) ML02212a expression around the pharynx revealed by ISH (**A**) and IHC (**B**).(**C-E**) ML199816 expression in different cell types of the AO; (**F**) ML199816 expression in pharynx neurons. (**G-S**) ML056913a is expressed in different cell types of AO (**G-I**), tentacles (**J**), pharynx and mouth (**K - O**) as well as subepithelial neurons on the body surface and under the combs (**P-S**).*

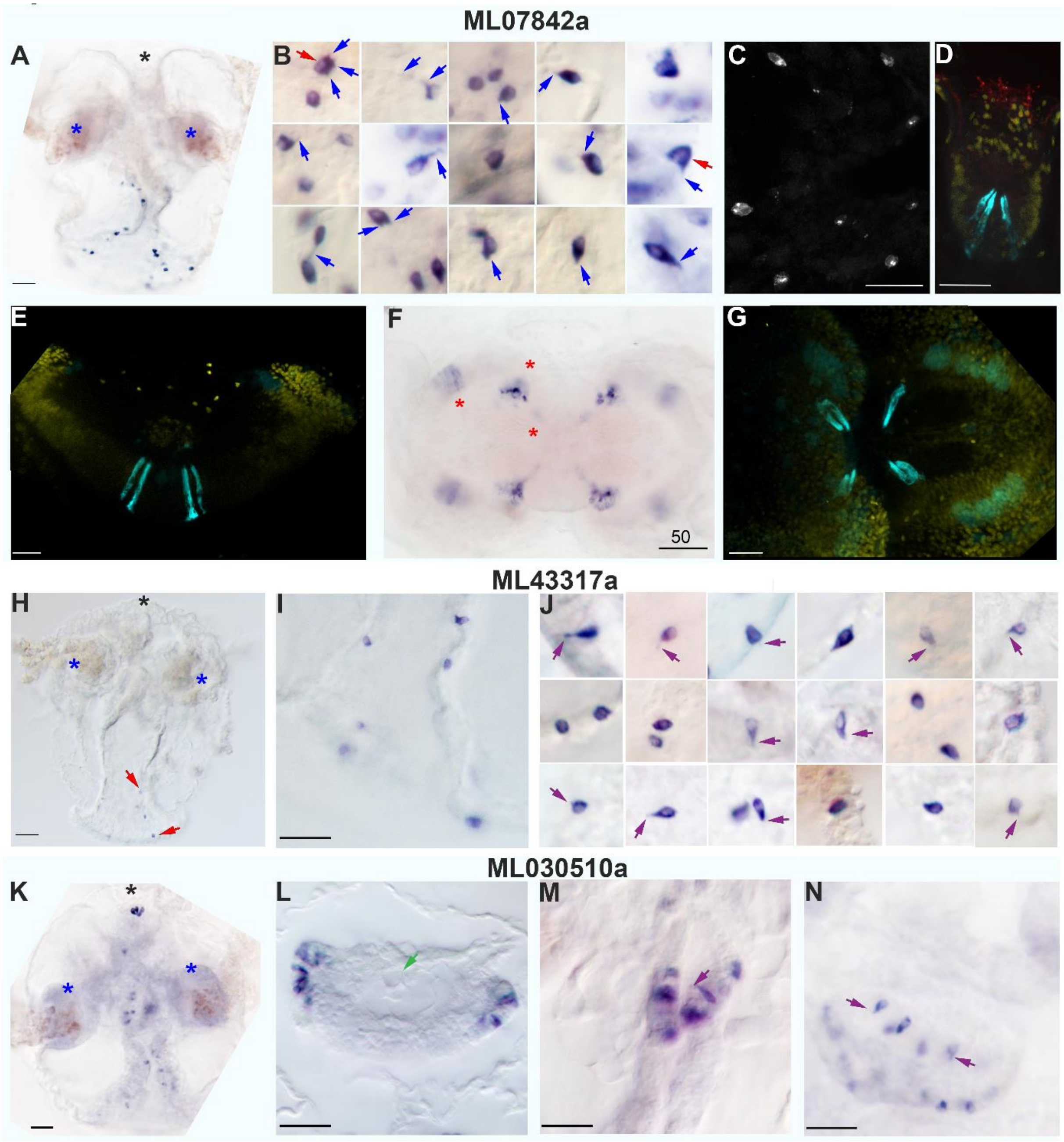


***Supplementary figure 9. Neuropeptides expressed in the pharynx and mouth area (ML07842a, ML43317a, ML030510a).*** *Star and arrow labels are the same as at the Fig 3 and 4, green arrows – statolith, light brown arrows – cilia. (**A**-**D**, **H**-**O)** ISH, (**E**-**G**) IHC. (**A, H, L**) whole cydippid, (**B, I**) close up to individual cells, (**C**) adult aboral organ, (**K, N, O**) pharynx, (**M**) cydippidl aboral organ. Scale bars: 50 µm for (**A, F, H**), 20 µm for (**C**-**E**, **G**, **I**, **L**-**N**).*

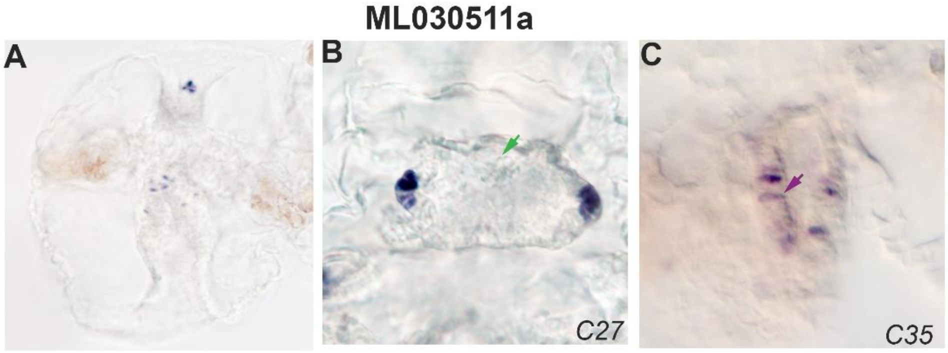


***Supplementary figure 10. ML030511a is expressed in the AO rim and sensory cells in the pharynx.*** *(**A**) whole cydippid; (**B**) AO, green arrow indicates the statholyth; (**C**) pharynx, purple arrow indicates the cilium.*

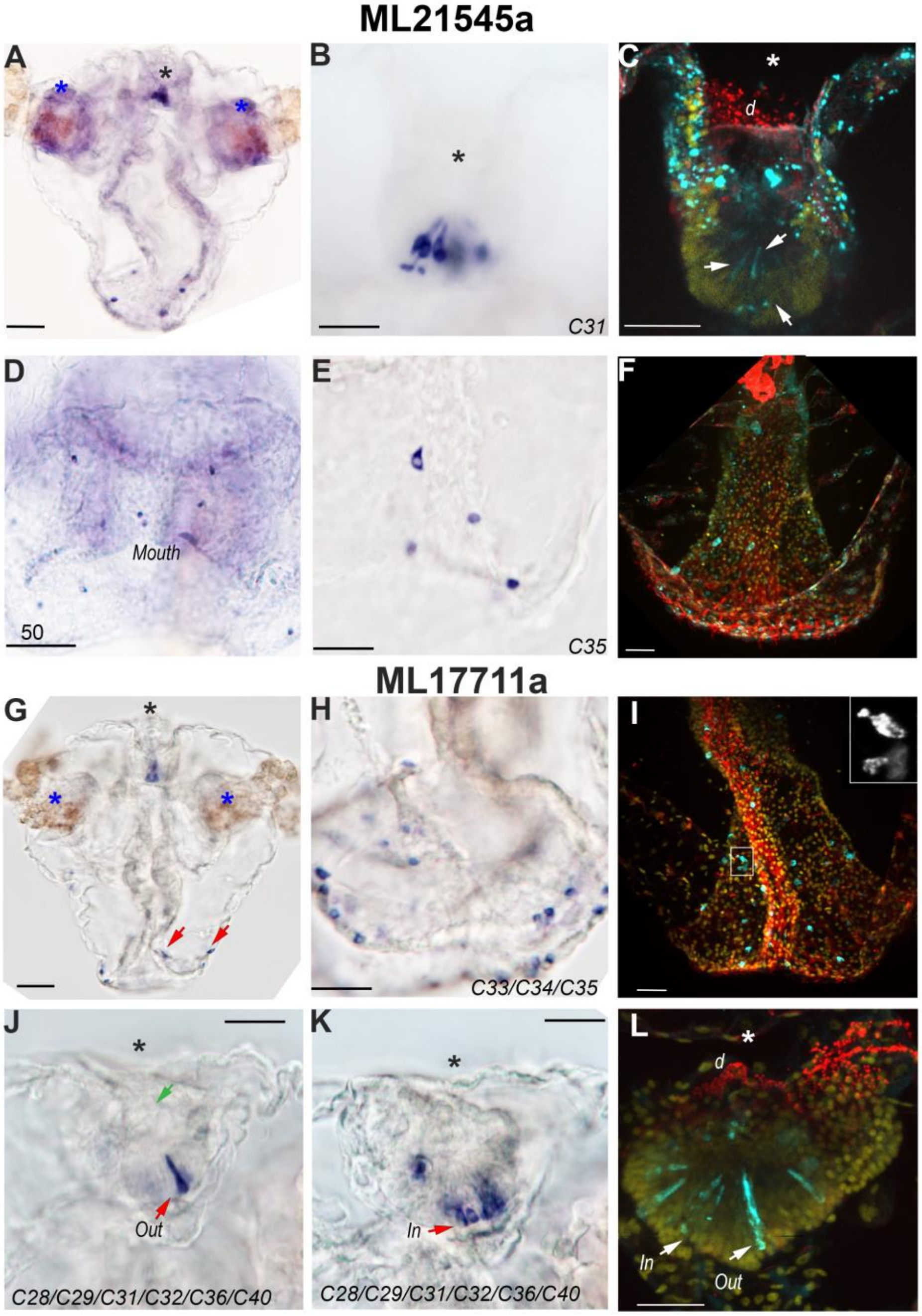


***Supplementary figure 11. Neuropeptides expressed in multiple cell types (ML21545a, A-H; ML17711a, I-P). (A, B, D, E)*** *ISH, (**C, F, I, L**) IHC.* ***(A, I**) whole cydippid, (**B, C, J, K**) aboral organ, (**E, H**) pharynx and lips area. Scale bars: 50 µm for (**A, G**), 20 µm for (**B-F, H-L**).*

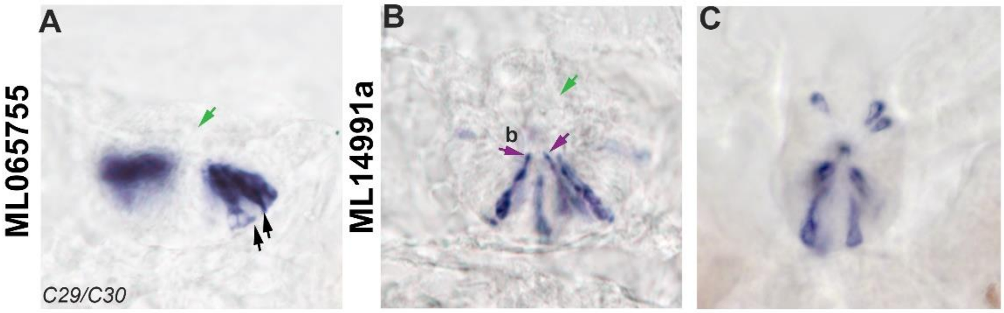


***Supplementary figure 12. ISH with neuropeptide precursors expressed in the AO cells. (A)*** *ML065755a: stained cells appear to have an elongated shape with the cell body close to the base of AO floor and both apical and basal protrusions (the latter is labelled with black arrows). The green arrow indicates the position of the statolith.* ***(B, C)*** *ML14991a is expressed in elongated cells with diverse morphology. Purple arrows indicate the cilium at the apical protrusion, green arrow – statolith, b – balancer.*

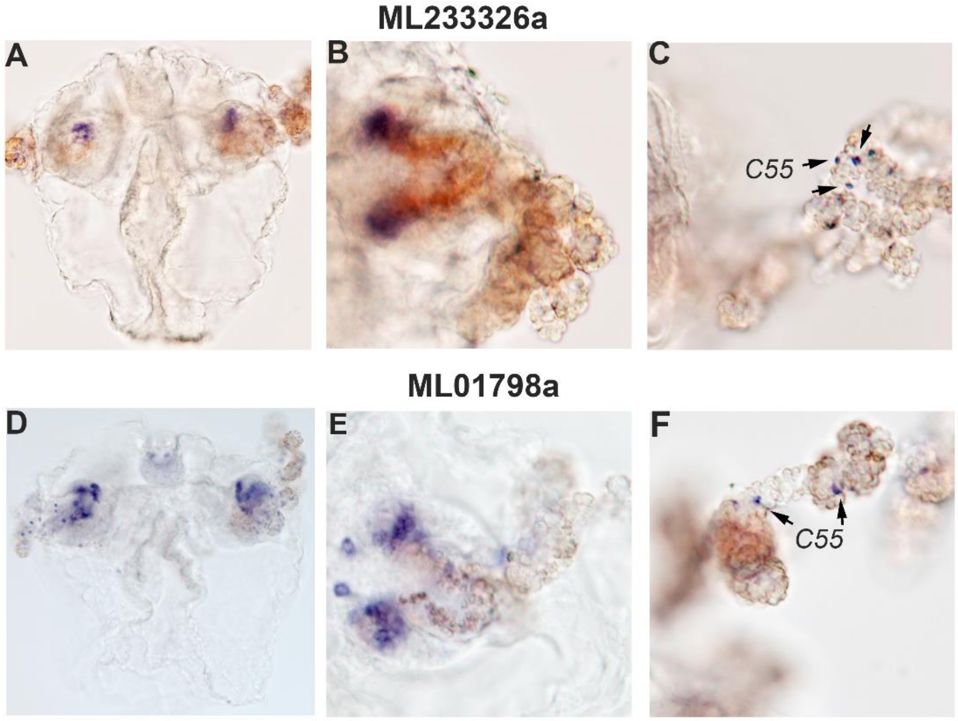


***Supplementary figure 13. ISH of neuropeptide precursor genes expressed in the tentacles.***
***(A-C)*** *ML233326a*, ***(D-F)*** *ML01798a*

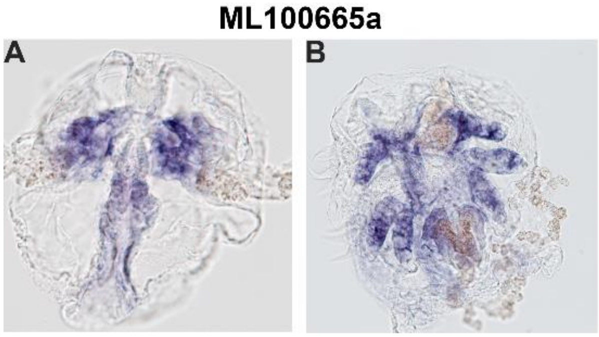


***Supplementary figure 14. ML10665a is expressed in the meridional canals and pharynx.***
***(A****) whole cydippid, front view****; (B)*** *whole cydippid, view from the AO side*

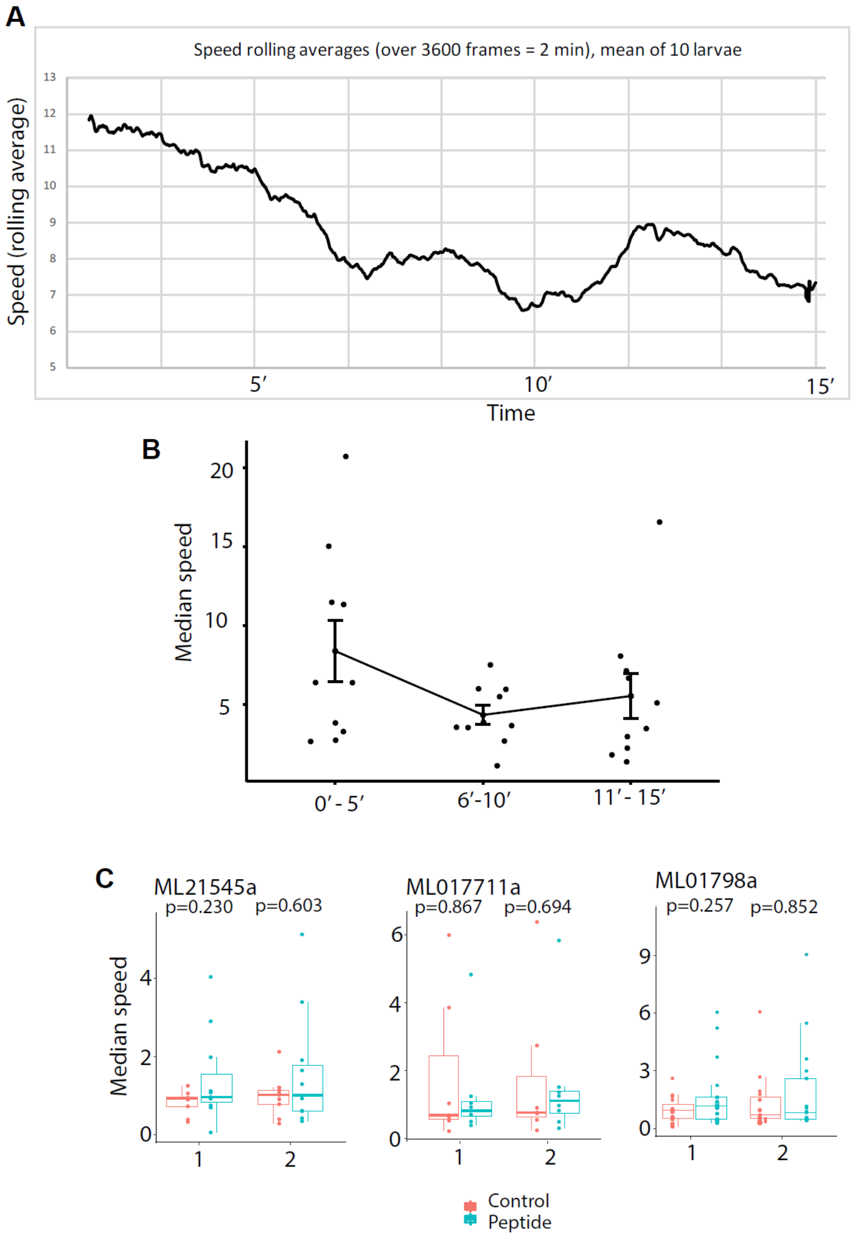


***Supplementary figure 15*. *Behavioural experiments with M. leidyi cydippids.*** *(**A**) Speed rolling averages (over 3600 frames corresponding to 2 min) for a cydippid recorded during 15 minutes after being placed into the arena; the mean for 10 cydippids is shown. (**B**) Median velocity calculated for the three 5 min intervals for the 15 min recordings of cydippids right after placement into arenas. (**C**) Plots showing median velocity of the cydippids after incubation with peptides or control (related to **Fig 3D**).*

**Ferlin (IPR037721)** (Reciprocal Blast eval 0.00)

ML08309a protein model appears truncated since it is missing the transmembrane region (TMR). GFAT01108989.1 cDNA overlaps with ML08309a and has a TMR (highlighted in blue).

```
sp|O75923|DYSF_HUMAN      --------------------------------MLRVFILYAENVHT-PDTDISDAYCSAV 27
GFAT01108989.1            ------------------------------------------------------------ 0
ML08309a                  MFGSIVRRSRSSDDDDHISSYREYKDVMSKEGLVQVLVRNAQNLQNVERFGYSDPHVVLE 60


sp|O75923|DYSF_HUMAN      FAGVKKRTKVIKNSVNPVWNEGFEWDLKGIPLDQGSELHVVVKDHETMGRNRFLGEAKVP 87
GFAT01108989.1            ------------------------------------------------------------ 0
ML08309a                  LEGIKRTTRVIHSELNPEWNETFTWKRY-RPLTEESMLLIKVYDYEKILKNKLLGEAVYP 119


sp|O75923|DYSF_HUMAN      LREVLATPSLSASFNAPLLDTKKQPTGASLVLQVSYTPL----------------P---- 127
GFAT01108989.1            ------------------------------------------------------------ 0
ML08309a                  LKDLVRLGAQEATV--PLRDRDGKVGESRLNLYLEYTKPTLDPDEEMEEEKSGPRPSVAR 177


sp|O75923|DYSF_HUMAN      ----------------GAVPLFPPPT------PLEPSPTLPDLDVVADTGGEEDTEDQGL 165
GFAT01108989.1            ------------------------------------------------------------ 0
ML08309a                  KSTALSHPEIQSLKNAEILPLYHSKRRSTTTSAPVSSPEYPAYHPTGVAGAVEETGNP-- 235


sp|O75923|DYSF_HUMAN      TGDEAEPFLDQSGGPGAPTTPRKLPSRPPPHYPGIKRKRSAPTSRKLLSDKPQDFQIRVQ 225
GFAT01108989.1            ------------------------------------------------------------ 0
ML08309a                  IPDEGEQEVEVAPGDAAGAAKK---------KKSVIAAPRSSIDRSKLSTKKQDFQVRVN 286


sp|O75923|DYSF_HUMAN      VIEGRQLPGVNIKPVVKVTAAGQTKRTRIHK-GNSPLFNETLFFNLFDSPGELFDEPIFI 284
GFAT01108989.1            ------------------------------------------------------------ 0
ML08309a                  IHEGRKLLGGNIHPVCNVHVGKQSKHTRVQKSTNKPLWDEVLFFDFNCSALDLCDEPVTI 346


sp|O75923|DYSF_HUMAN      TVVDSRSLRTDALLGEFRMDVGTIYREPRHAYLRKWLLLSDPDDFSAGARGYLKTSLCVL 344
GFAT01108989.1            ------------------------------------------------------------ 0
ML08309a                  EVLNSRKIRSDSLIGAFKFDLGLVYESQDHQFVHKWVLLTDPEDKESGAKGYVKISISIL 406


sp|O75923|DYSF_HUMAN      GPGDEAPLERKDP-SEDKEDIESNLLRPTGVALRGAHFCLKVFRAEDLPQMDDAVMDNVK 403
GFAT01108989.1            ----------------------------------------------------------MK 2
ML08309a                  GPGDKLKIPPKSSSTDDLVDIESNLLRPAGVQLQPATYTVKIYKAEDVPKMDTDYFEGMK 466
                                                                                    :*

sp|O75923|DYSF_HUMAN      QIFGFESNKKNLVDPFVEVSFAGKMLCSKILEKTANPQWNQNITLPAMFPSMCEKMRIRI 463
GFAT01108989.1            RVLRMQHGDHDLVDPYMIVSFAGKKLKTKVLYKTYTPEWAQELNIGVQMPSMCEQLMLRL 62
ML08309a                  RVLRMQHGDHDLVDPYMIVSFAGKKLKTKVLYKTYTPEWAQELNIGVQMPSMCEQLMLRL 526
                          ::: :: ..::****:: ****** * :*:* ** .*:* *::.: . :*****:: :*:

sp|O75923|DYSF_HUMAN      IDWDRLTHNDIVATTYLSMSKISAPGGEIEEEPAGAVKPSKASDLDDYLGFLPTFGPCYI 523
GFAT01108989.1            MDKDHFNRDDIIATHFLQLTRLSSAD-------------------PDDEGFLPTFGPAYV 103
ML08309a                  MDKDHFNRDDIIATHFLQLTRLSSAD-------------------PDDEGFLPTFGPAYV 567
                          :* *::.::**:** :*.::::*: .                    *  ********.*:

sp|O75923|DYSF_HUMAN      NLYGSPREFTGFPDPYTELNTGKGEGVAYRGRLLLSLETKLVEH--SEQKVEDLPADDIL 581
GFAT01108989.1            NFYGSPRESI-VDDELEPLNRGCGEGCSFRGRALVELTVNIGQEPSKDEMLRDIDGEDWL 162
ML08309a                  NFYGSPRESI-VDDELEPLNRGCGEGCSFRGRALVELTVNIGQEPSKDEMLRDIDGEDWL 626
                          *:******   . *    ** * *** ::*** *:.* .:: :.  .:: :.*: .:* *

sp|O75923|DYSF_HUMAN      RVEKYLRRRKYSLFAAFYSATMLQDVDDAIQFEVSIGNYGNKFDMTCLPLASTTQYSRAV 641
GFAT01108989.1            RVQPFQRRRRYRLFVGFLEGTMIHPVDAPVEFEISIGEYGNKFASTTLPAPSTTQPTNPV 222
ML08309a                  RVQPFQRRRRYRLFVGFLEGTMIHPVDAPVEFEISIGEYGNKFASTTLPAPSTTQPTNPV 686
                          **: : ***:* **..* ..**:: **  ::**:***:*****  * **  **** :. *

sp|O75923|DYSF_HUMAN      FDGCHYYYLPWGNVKPVVVLSSYWEDISHRIETQNQLLGIADRLEAGLEQVHLALKAQCS 701
GFAT01108989.1            YDGSHYYYLPWGQTKPCVMLNSQWEDVMFRIESLNLICRTIDRLEGHLKDVKYLMRAKAP 282
ML08309a                  YDGSHYYYLPWGQTKPCVMLNSQWEDVMFRIESLNLICRTIDRLEGHLKDVKYLMRAKAP 746
                          :**.********:.** *:*.* ***: .***: * :    ****. *::*:  ::*:.

sp|O75923|DYSF_HUMAN      TEDVDSLVAQLTDELIAGCSQPLGDIHETPSATHLDQYLYQLRTHHLSQITEAALALKLG 761
GFAT01108989.1            LAESAGKLIKLLDELIIDLNQPLLELPQK-NITELDKKLYALRDDEMKRILIEAVNLREN 341
ML08309a                  LAESAGKLIKLLDELIIDLNQPLLELPQK-NITELDKKLYALRDDEMKRILIEAVNLREN 805
                            :  . : :* **** . .*** :: :. . *.**: ** ** ..:.:*   *: *: .

sp|O75923|DYSF_HUMAN      HSELPAALEQAEDWLLRLRALAEEPQNSLPDIVIWMLQGDKRVAYQRVPAHQVLFSRRGA 821
GFAT01108989.1            ARDIDQAIVEIDSYIYRLQQIATEPQNSIPDVIIWMLCGNRRVAYHRIPSHQVMFSPK-Q 400
ML08309a                  ARDIDQAIVEIDSYIYRLQQIATEPQNSIPDVIIWMLCGNRRVAYHRIPSHQVMFSPK-Q 864
                            ::  *: : :.:: **: :* *****:**::**** *::****:*:*:***:** :

sp|O75923|DYSF_HUMAN      NYCGKNCGKLQTIFLKYPMEKV-----PGARMPVQIRVKLWFGLSVDEKEFN-QFAEGKL 875
GFAT01108989.1            DCCGKLCGHVFSVFLKRPSVPDKSNAKRQWKLPAKLQVFVWMGLEDHAKGIQHKPLDGEI 460
ML08309a                  DCCGKLCGHVFSVFLKRPSVPDKSNAKRQWKLPAKLQVFVWMGLEDHAKGIQHKPLDGEI 924
                          : *** **:: ::*** *            ::*.:::* :*:**. . * :: :  :*::

sp|O75923|DYSF_HUMAN      SVFAETYENETKLALVGNWGTTGLTYPKFSDVTGKIKLPKDSFRPSAGWTWAGDWFVCPE 935
GFAT01108989.1            SVFAETYENQ--ISLLSKWTTRAMPRPKWSDITGQLKLPKESFTTPGGWRWAGEWFINPN 518
```

```
ML08309a                   SVFAETYENQ--ISLLSKWTTRAMPRPKWSDITGQLKLPKESFTTPGGWRWAGEWFINPN  982
                           *********:  ::*:.:* * .:  **:**:**::****:**   .** ***:**: *:

sp|O75923|DYSF_HUMAN       KTLLHDMDAGHLSFVEEVFENQTRLPGGQWIYMSDNYTDVNGEKVLPKDDIECPLGWKWE  995
GFAT01108989.1             LSLSYDLDSGLSSFQDDVFENQLRVPGSDWPTSKLFWTDVTGEEAQSKEDIMCPAGWEWT  578
ML08309a                   LSLSYDLDSGLSSFQDDVFENQLRVPGSDWPTSKLFWTDVTGEEAQSKEDIMCPAGWEWT  1042
                            :* :*:*:*  ** ::***** *:**.:*    .   :***.**:.  *:** ** **:*

sp|O75923|DYSF_HUMAN       DEEWSTDLNRAVDEQGWEYSITIPPERKPKHWVPAEKMYYTHRRRRWVRLRRRDLSQMEA  1055
GFAT01108989.1             D-IWTVDLNRAVDEEGYEYCLDQ----SVGGFVPVEKTYHLCRRRRWVRTRKRNPDLRQQ  633
ML08309a                   D-IWTVDLNRAVDEEGYEYCLDQ----SVGGFVPVEKTYHLCRRRRWVRTRKRNPDLRQQ  1097
                           *  *:.********:*:**.:      .   :**.** *:  ******* *:*: .  :

sp|O75923|DYSF_HUMAN       LKRHRQAEAEGEGWEYASLFGWKFHLEYRKTDAFRRRRWRRRMEPLEKTGPAAVFALEGA  1115
GFAT01108989.1             AAHQRMVRAAEEGWEYSRLFTTKFHLKQRTMDMVRRRRWHRKMVADNPDA-DAIFIIDPT  692
ML08309a                   AAHQRMVRAAEEGWEYSRLFTTKFHLKQRTMDMVRRRRWHRKMVADNPDA-DAIFIIDPT  1156
                             ::* ..*  *****: **  ****: *. * .*****:*:*   :  .  *:* :: :

sp|O75923|DYSF_HUMAN       LG-----GVMD------DKSEDSMSVSTLSFGVNRPTISCIFDYGNRYHLRCYMYQARDL  1164
GFAT01108989.1             SDAHETVGVSRARSPARGKDDDHLQ----QRTMITPTVFLTYKEPHTYQLRAYIYQARDL  748
ML08309a                   SDAHETVGVSRARSPARGKDDDHLQ----QRTMITPTVFLTYKEPHTYQLRAYIYQARDL  1212
                            .     **        .*.:* :.    .  :  **:   :.  : *:**.*:******

sp|O75923|DYSF_HUMAN       AAMDKDSFSDPYAIVSFLHQSQKTVVVKNTLNPTWDQTLIFYEIEIFGEPATVAEQPPSI  1224
GFAT01108989.1             FSADPSGLSDPYARVVFSRQSQRTKILNETLCPTWDQTLVFEEVEFYGNPTMLAESPPIV  808
ML08309a                   FSADPSGLSDPYARVVFSRQSQRTKILNETLCPTWDQTLVFEEVEFYGNPTMLAESPPIV  1272
                            : * ..:***** * * :***:* ::::** *******:* *:*::*:*: :**.** :

sp|O75923|DYSF_HUMAN       VVELYDHDTYGADEFMGRCICQPSL-----ERMPRLAWFPLTRGSQPSGELLASFELIQR  1279
GFAT01108989.1             VVELFDYDTVGS-DFLGRAIATPIVKLGGEHQMAKLNWHPITRGGEPAGELLGAFELYLN  867
ML08309a                   VVELFDYDTVGS-DFLGRAIAYPDSEAGR-------------------------------  1300
                           ****:*:** *: :*:**.*. *

sp|O75923|DYSF_HUMAN       EKPAIHHIPGFEVQETSRILDESEDTDLPYPPPQREANIYMVPQNIKPALQRTAIEILAW  1339
GFAT01108989.1             EG----------------------AELPFMPPTR-GDVYQVPSGIRPVMQLTRIEVLTW  903
ML08309a                   ------------------------------------------------------------  1300

sp|O75923|DYSF_HUMAN       GLRNMKSYQLANISSPSLVVECGGQTVQSCVIRNLRKNPNFDICTLFMEVMLPREELYCP  1399
GFAT01108989.1             GVRHMKKFQLAAVNSPSIEIECGGVVLSTIKIKNAKKNPNFDTSSMLFDVFLPVEELYTP  963
ML08309a                   ------------------------------------------------------------  1300

sp|O75923|DYSF_HUMAN       PITVKVIDNRQFGRRPVVGQCTIRSLESFLCDPYSAESPSPQ----GGPDDVS--LLSPG  1453
GFAT01108989.1             PLNIRLLDHRSFGVKPLVGTHMIKSLQEYRRDPVAMVQTIREKMLESGFGDAEYAIDMDF  1023
ML08309a                   ------------------------------------------------------------  1300

sp|O75923|DYSF_HUMAN       EDVLIDIDDKEPLI-----PIQEEEFIDWWSKFFASIGEREKCGSYLEKDFDTLKVYDTQ  1508
GFAT01108989.1             ADSAVEAAHQPPATSTSSEEGEFKEDVDWWSKYYSSSGNEQLGRVYREKGYENMVVFEEE  1083
ML08309a                   ------------------------------------------------------------  1300

sp|O75923|DYSF_HUMAN       LENVEAFEGLSDFCNTFKLYRGKTQEE--TEDPSVIGEFKGLFKIYPLPEDPAIPMPPRQ  1566
GFAT01108989.1             LEN--YFDNFTDLAQSFPLFHGKRHEDEDLDEDKAVGYFKGTFRVYPLPADGSDP-PPRM  1140
ML08309a                   ------------------------------------------------------------  1300

sp|O75923|DYSF_HUMAN       FHQLAAQGPQECLVRIYIVRAFGLQPKDPNGKCDPYIKISIGKKSVSDQDNYIPCTLEPV  1626
GFAT01108989.1             LKNVPCNQLVEVLVRVYIVKAFELQPQDPNGLSDPYLALKLGRFKVKDRENYVPKNLSPT  1200
ML08309a                   ------------------------------------------------------------  1300

sp|O75923|DYSF_HUMAN       FGKMFELTCTLPLEKDLKITLYDYDLLSKDEKIGETVVDLENRLLSKFGARCGLPQTYCV  1686
GFAT01108989.1             FGKMFELDGTLPLESELRVQIFDYDLLSGDDLIGETKIDLENRFLSARRGVCGLPKRYYV  1260
ML08309a                   ------------------------------------------------------------  1300

sp|O75923|DYSF_HUMAN       SGPNQWRDQLRPSQLLHLFCQQHRVKAPVYRTDRVMFQ-DKEYSIEEIEAGRIPNPHLGP  1745
GFAT01108989.1             AGPYKWRDAELPRQILEKWCATQALPPPVWRGNSQVYVNGKTCCLRDYETRGQVHKDWGP  1320
ML08309a                   ------------------------------------------------------------  1300

sp|O75923|DYSF_HUMAN       VEERLALHVLQQQGLVPEHVESRPLYSPLQPDIEQGKLQMWVDLFPKALGRPGPPFNITP  1805
GFAT01108989.1             PEERLALYTLLDLGLVPEHIETRTLYNPLRPEIPQGKIQIFVDIFPKSATI-PPPINITP  1379
ML08309a                   ------------------------------------------------------------  1300

sp|O75923|DYSF_HUMAN       RRARRFFLRCIIWNTRDVILDDLSLTGEKMSDIYVKGWMIGFEEHKQKTDVHYRSLGGEG  1865
GFAT01108989.1             RAPQDLQLRVIVYNVQDVVLSDTSFTGEKMSDIYVKGWLKGQD-KKQKTDVHYRSLNGEG  1438
ML08309a                   ------------------------------------------------------------  1300

sp|O75923|DYSF_HUMAN       NFNWRFIFPFDYLPAEQVCTIAKKDAFWRLDKTESKIPARVVFQIWDNDKFSFDDFLGSL  1925
GFAT01108989.1             NFNWRYVFPFKYLPAEEVMVIKKKEHFFSLDKHEEKHPVTFVCQIWDNDIFTPDDFLGLL  1498
ML08309a                   ------------------------------------------------------------  1300
```

```
sp|O75923|DYSF_HUMAN      QLDLNRMPKPAKTAKKCSLDQLDDAF--HPEWFVSLFEQKTVKGWWPCVAEEGE--KKIL 1981
GFAT01108989.1            ELNLNFMPKANKFAKSVSLDDLPDNEKGKPVPMVSLFDQKNVKGWWPMYEEGGPDQPREL 1558
ML08309a                  ------------------------------------------------------------ 1300


sp|O75923|DYSF_HUMAN      AGKLEMTLEIVAESEHEERPAGQGRDEPNMNPKLEDPRRPDTSFLWFTSPYKTMKFILWR 2041
GFAT01108989.1            TGKVEMELEILSKEDAEAKPAGKGQEEPNENPHLDPPNRPATSFLWFTSPWKSLRYIIWN 1618
ML08309a                  ------------------------------------------------------------ 1300


sp|O75923|DYSF_HUMAN      RFRWAIILFIILFILLLFLAIFIYAFPNYAAMKLVKPFS 2080    100%
GFAT01108989.1            NYKWYIIGGLVLLLIIAMVGLFIYSAPGALSTKLIAKL- 1656     46%
ML08309a                  --------------------------------------- 1300     42%
```

#### Secretagogin (Reciprocal Blast eval $9.38 \times 10^{-28}$)

```
sp|O76038|SEGN_HUMAN      -------MDSSREPTLGRL--------DAAGFWQVWQRFDADEKGYIEEKELDAFFLHML 45
ML03617a                  MEISGWLNDPQRVPKVKRKVMERKFRISPATLRNELENNDVYNNNYLETVLLDEFLTNLV 60
                                 * .* *.: *         . * : :  :. *. ::.*:*   ** *: :::

sp|O76038|SEGN_HUMAN      MKLGTDDTVMKANLHKVKQQFMTTQDASKDGRI--------R---MKELAGMFLSEDENF 94
ML03617a                  MKE---------YVWQLNRQDIKCFNTEVIGNIKETLGDKLPTKINKEVLIAAIPAENNY 111
                          **           : ::::* :.  ::.  *.*              **:    :  ::*:

sp|O76038|SEGN_HUMAN      LLLFRRENPLDSSVEFMQIWRKYDADSSGFISAAELRNFLRDLFLHHKKAISEAKLEEYT 154
ML03617a                  LVRIG---HRIPSSDFLKLWRNYDTDHSGYLEIKELQLLVRDFAQLVGEDVSDKDLNTAL 168
                          *: :        * :*:::**:**:* **::.  **: ::**:     : :*: .*:

sp|O76038|SEGN_HUMAN      GTMMKIFDRNKDGRLDLNDLARILALQENFLLQFKMDACSTEERKRDFEKIFAYYDVSKT 214
ML03617a                  EELMSEFDVNKDGRLELEELSHLMSVEDNFMKAFCSRQ---YLTRKDFDRIFAHYDTDST 225
                            :*. ** ******:*::*::::::::**:  *          ::**::***:**...*

sp|O76038|SEGN_HUMAN      GALEGPEVDGFVKDMMELVQPSISGVDLD---KFREILLRHCDVNKDGKIQKSELALCLG 271
ML03617a                  GYLDKEEVMALLNDILKYHESSDAHIPVPVLKEVYKEVMKACDTNNSNTIQKCELALLLT 285
                          * *:  ** .:::*:::  : * : : :    :. : ::: **.*:...***.**** *

sp|O76038|SEGN_HUMAN      LKINP  276  100%
ML03617a                  SV---  287   29%
```

#### VAMP2 = Synaptobrevin (IPR001388) (Reciprocal Blast eval $3.99 \times 10^{-22}$, $1.78 \times 10^{-23}$)

SNARE domain, TMR

Residues important for SNARE complex formation are highlighted in grey and cyan.

```
VAMP2_HUMAN       MSATAATAPPAAPAGEGGPPAPPPNLTSNRRLQQTQAQVDEVVDIMRVNVDKVLERDQKL         60
ML02217a          MAELVAM----------VTLYHPLTSSGGYQTRDTQEQVDQVMGIMRNNIDKVLERDSKI         50
ML214317a         -------------------------MSGGYQVNDTQNQVDEVMGIMRNNIDKVLERDTKI         35
                                           :.. :  .:** ***:*:.*** *:******* *:

VAMP2_HUMAN       SELDDRADALQAGASQFETSAAKLKRKYWWKNLKMMIILGVICAIILIIIIVYFST----         116
ML02217a          QNLNERSDALQVGANQFLQTGTQLKRKMWWKNVKFMIVIGVVVVIVLGVVIVLLDFVTIN         110
ML214317a         QNLNERSDALQVGAHQFQQTGQQLKRKMWWKNVKFMIIIGVVVVVIVGILIGIIVSQTKK         95
                  .:*::*:****.** **  :. :**** ****:*:**::**: .::: ::*  :

|VAMP2_HUMAN       ------------------------------------------------------------        116
ML02217a           RPNVPLSWSLQTRSSMNTELYSVSTYLPHKSWLLEIKPGSLRNCVSKLLLPRLSLRIVSR        170
ML214317a          KKSN--------------------------------------------------------        99


VAMP2_HUMAN       ------------------------------------------------------------         116
ML02217a          EGLTGVLTLSILRSSLSLPRLVAANSSSLTDRPTANQPPPTTSSTNHRRPFLYSTNKERG         230
ML214317a         ------------------------------------------------------------         99


VAMP2_HUMAN       --------------        116     100%
ML02217a          IFITVQSTTPASLS        244      45%
ML214317a         --------------        99       51%
```

#### Syntaxin1 or Syntaxin-2 (IPR028671) (Reciprocal Blast eval $9.57 \times 10^{-45}$)

SNARE domain, TMR

Residues important for SNARE complex formation are highlighted in grey and cyan.

```
STX1B_HUMAN       MKDRTQELRSAKDSDD----EEEVVHVDRDH---FMDEFFEQVEEIRGCIEKLSEDVEQV         53
ML037014a         MRDRINFFRLQLDDQEANEGRATVVDFHSDTGVEEVKKFLKEAQHTREKIVSIEELVNQI         60
```

```
                 *:** : :*   *.::    .  **... *     :.:*:::.:. *  * .:.* *:*:

STX1B_HUMAN      KKQHSAILAAPNPDEKTKQELEDLTADIKKTANKVRSKLKAIEQSI----EQEEGLNRSS       109
ML037014a        REYHGKITGAAARNEEVHKKLNQAMDEIRRLFDTVKEALKKMEQESNALSDKKDAELKRP       120
                 :: *. * .*   :*:.:::*::   :*::  :.*:. ** :**.     ::::.  :

STX1B_HUMAN      ADLRIRKTQHSTLSRKFVEVMTEYNATQSKYRDRCKDRIQRQLEITGRTTTNEELEDMLE       169
ML037014a        ADIRIMSSQYTSLQCWFFETWTEYNECQSDYREKCKEKLTRQIQITEKKVTQEEINTMIE       180
                 **:** .:*:::*.  *.*. ****  **.**::**::: **::** :..*:**:: *:*

STX1B_HUMAN      SGKLAIFTDDIKMDSQMTKQALNEIETRHNEIIKLETSIRELHDMFVDMAMLVESQGEMI       229
ML037014a        SGNFTVFSIN----AQFQPKDIEEMESRHNDITKLAKSLKQLHEMFKDLALMVEQQGEML       236
                 **::::*: :    :*:  : ::*:*:***:* ** .*:::**:** *:*::**.****:

STX1B_HUMAN      DRIEYNVEHSVDYVERAVSDTKKAVKYQSKARRKKIMIIICCVVLGVVLASSIGGTLGL-       288
ML037014a        NNIEKNVDQARDYVADAERECHQAVDLTNKWRKKKLICAVVILI----VVIVGAVIAVV       292
                 :.** **::: ***  *  : ::**.  .* *:**::  :  :::    .  :*..:.:

STX1B_HUMAN      ------      288   100%
ML037014a        AHLQTK      298    33%
```

#### SNAP25 (Reciprocal Blast eval 2.76 × 10-51)

SNARE domain

Residues important for SNARE complex formation are highlighted in grey and cyan.

```
SNP25_HUMAN      MAEDADMRNELEEMQRRADQLADESLESTRRMLQLVEESKDAGIRTLVMLDEQGEQLERI       60
ML26853a         --MSGRTPVQMSDIEIRGNALTDSTLEATREMKRLAAETREIGAATAQNLDRQGEQLNNI       58
                    ..    ::.::: *.: *:*.:**:**.* :*. *::: *  *   **.*****:.*

SNP25_HUMAN      EEGMDQINKDMKEAEKNLTDLGKFCGLCVCPCNKLKSSDA-------YKKAWGNNQDGVV       113
ML26853a         ENNLDEINADLHKAEKHLTNMEKCCGICLCPCARPKGVGGTKTRNDKWKAAQSVNNGGTV       118
                 *:.:*:** *:::***:**:: * **:*:*** : *. ..       :* *  . *:.*.*

SNP25_HUMAN      ASQPARVVDEREQMAISGGFIRRVTNDARENEMDENLEQVSGIIGNLRHMALDMGNEIDT       173
ML26853a         TTQPHGYQK---EEVAKTGYIKRITNDAREDEMDENLAVVADVVGDLKMMAMDMGSELER       175
                 ::**    .   : . . *:*:*:******:******  *:.::*:*: **:***.*::

SNP25_HUMAN      QNRQIDRIMEKADSNKTRIDEANQRATKMLGSG       206    100%
ML26853a         QNAQLDRINVKTDCNVNRVQDAEARTNKLLNS-       207     42%
                 ** *:*** *:*.* .*:::*: *:.*:*.*
```

#### Synaptotagmin-7 (Reciprocal Blast eval 1.05 × $10^{-21}$)

TMR, InerPro C2_dom

```
SYT7_HUMAN     MYRDPEAASPGAPSR-DVL--LVSAIITVSLSVTVVLCGLCHWCQRKLGKRYKNSLETVG 57
ML16217a       -------MSLSLSDELTVLLCVLGVIIFIF---FVFMVFLCFECLRQKEN---------- 40
                       *  .  ..  **  ::..** :      *.:  **. * *:  :

SYT7_HUMAN     TPDSGRGRSEKKAI--KLPAGGKAVNTAPVPGQTPHDESDRRTEPRSSVSD----LVNSL 111
ML16217a       EPEGGAGDNLPFSFLSFLSVNQKGDNSERIGNKNIFSFSKRKEQPPAAVPTETYVLNDSI 100
                *:.* * .   ::   *  .. *. *:  : .:. .. *.*: :* ::*       *  :*:

SYT7_HUMAN     TSEMLMLSPGS---------------------EEDE----AHEGCSRENLGRIQFSVGYN 146
ML16217a       KSSYVDIVPQSKVIYKSVAQDSDSESQFSFTSEPDQRRLSHFAPLEASTCGTIVCSIKFY 160
               .*. : : * *                     * *:      .    . .. * *   *: :

SYT7_HUMAN     FQESTLTVKIMKAQELPAKDFSGTSDPFVKIYLLPDKKHKL--ETKVKRKNLNPHWNETF 204
ML16217a       HFANRLAVRVGEVQLNL--VENLAVNPYVKLHLLPEYKRGNRQTTRVKRNNTLFVCDEDF 218
               .  . *:*:: :.*        . : :*:**::***: *:      *:***:*      :* *

SYT7_HUMAN     LFEGFPYEKVVQRILYLQVLDYDRFSRNDPIGEVSIPLNKVDLTQMQT---FWKDLKPCS 261
ML16217a       LF-GVGSREVRTKSLSLQVYDYKSNTRHLCIGKVNINLGDYDFKDEDKPLSLKRHIIPYR 277
               ** *.  .:*  : * *** **.  :*:  **:*.* *.. *:.: :.   : :.: *

SYT7_HUMAN     DGSGSRGELLLSLCYNPSANSIIVNIIKARNLKAMDIGGTSDPYVKVWLMYKDKRVEKKK 321
ML16217a       ESQEHFGAIQV--AVSIASETLRIGIIKAEGLVPVDMSGTLEPYCRILVHVGSELIHKKK 335
               :..    * : :   . . ::::: :.****..*  :*:.** :** :: :   .: :.***

SYT7_HUMAN     T-VTMKRNLNPIFNESFAFDIPTEKLRET-TIIITVMDKDKLS---RNDVIGKIYLSWKS 376
ML16217a       TCIAEAINRSPIWEEFFTLDLPKNTLLCETSITIEVRDHLRNSRSTSYLLMGRVILSNTA 395
```

```
                 * ::   * .**::* *::*:*.:.*    :* * * *: : *      ::*:: ** .:

SYT7_HUMAN       GPGEVKHWKDMIARPRQPVAQWHQLKA     403   100%
ML16217a         PGHGQAHWEHASAQRGTMITEWQPLLS     422    27%
                        **:.  *:    :::*: * :
```

#### **Dematin** (Reciprocal Blast eval 1.17 × $10^{-09}$)

##### Villin_headpiece domain

```
sp|Q08495|DEMA_HUMAN      ------------------------------------------------------------  0
ML334211a                 MSRSFQDPTTPSRPVVLGGSFIFSNSCDGYISPTVIEKYESLAGDLSLDKENLLNIEKSL  60

sp|Q08495|DEMA_HUMAN      ------------------------------------------------------------  0
ML334211a                 DKGIVTEDVGANQDIISFTTSPPCVESSPAMAEKPANPKKVELVGPYISPHKKDIITFFL  120

sp|Q08495|DEMA_HUMAN      ------------------------------------------------------------  0
ML334211a                 TDIGRQKTMEQDGACYGTSVSVETVGETTPDCVTAERDDKKDENNKDIISFLTAGTNVDA  180

sp|Q08495|DEMA_HUMAN      ------------------------------------------------------------  0
ML334211a                 DNSRKLSYSLSVPPPKSKADLRKPYFSLNFSDIISFYNSEESLLHACSMNSDQEPQGLDS  240

sp|Q08495|DEMA_HUMAN      ------------------------------------------------------------  0
ML334211a                 QKEADDIVQDPKSPDKEKIPANNPDQFVFQNDRSANANKVLTSTPKSTLKKRTTPMMIHQ  300

sp|Q08495|DEMA_HUMAN      ------------------------------------------------------------  0
ML334211a                 GSVISFYSKEQSFLSALEVSSVDGDDINASAEDQDIEEAFKQNKDKCESTVSTANEQEIS  360

sp|Q08495|DEMA_HUMAN      ------------------------------------------------------------  0
ML334211a                 NSDIKVFSNRATEPLPNSFLKQPFKSEPQLRKSNSVLVVTEEFPDIISFYNPGADDTDPT  420

sp|Q08495|DEMA_HUMAN      ------------------------------------------------------------  0
ML334211a                 SPAGATLEVKDSMRKRENSTSLDTAEFGIIELTQLNENMAQVDTEVATAPDEEPFIVIEN  480

sp|Q08495|DEMA_HUMAN      ------------------------------------------------------------  0
ML334211a                 PASEEADEVNPDKFVFRNDRSGFVEKNVISPANQNVRTEEKQTPMMINQGSVITFYRNET  540

sp|Q08495|DEMA_HUMAN      ------------------------------------------------------------  0
ML334211a                 SFLTALNSPNTVQTDESIENDINSPSVDVAEDEADKNVTSESNFDIKVYENTVLEHDTSM  600

sp|Q08495|DEMA_HUMAN      ------------------------------------------------------------  0
ML334211a                 EDDNECLKRSLPDDDEEAPIPEKIEIEDQISKPVTTNIKVDDASIPDVVKETEKEDIERL  660

sp|Q08495|DEMA_HUMAN      ------------------------------------------------------------  0
ML334211a                 VHIVEMNEHNQDSDYDDDMPGLEECPKSIHESDLSENQDSLVVVEDSISEDGIPIRKLQD  720

sp|Q08495|DEMA_HUMAN      ------------------------------------------------------------  0
ML334211a                 IENLPSIEKAIDMVNAIGGALDENLGMSVPEPLDDDYMYDDEEFERDITEEIDSLISEAE  780

sp|Q08495|DEMA_HUMAN      ------------------------------------------------------------  0
ML334211a                 QLMMPTDSADNDALCDDLDRFFDEVDDVDELLLKAEPDEVPDLQIRNVSQNGETEISPRD  840

sp|Q08495|DEMA_HUMAN      ------------------------------------------------------------  0
ML334211a                 SSSESAIINFITQNIDKYMESTSAVDAAVATAPPTNVDPSTDDADDLVARIDELLDVEDY  900

sp|Q08495|DEMA_HUMAN      ------------------------------------------------------------  0
ML334211a                 LDIPGCETPEPPSPLRDTPTPQPHTPTVTRHLIRSTSLGEQALGLSLERTKGSFSAEDDG  960

sp|Q08495|DEMA_HUMAN      ------------------------------------------------------------  0
ML334211a                 GSDEEYHTLFEALNKPILPPIPRVEYDEEDNQIAPAKSLTALQDYVPDPANLSLQPSNNP  1020

sp|Q08495|DEMA_HUMAN      ------------MERLQKQPL---------------------------------------  9
ML334211a                 CDPDSKVLSTGLLSEIVRHPLAGSLKRSASSGDAPGYHRRNPVTRSQKNDLLDEAHLFFL  1080
                                       :..: ::**

sp|Q08495|DEMA_HUMAN      -----------------------TSPGSVSPS----------------RDSSVPGSPSSI  30
ML334211a                 QNNIGVPSRGVGDDPSNREKKPSEDDQELSPISSLAQTLTERLTAHTAGRAGVKMVKPSD  1140
```

```
                                                   .  .:**                   :.*     *

sp|Q08495|DEMA_HUMAN       VAKMDNQVLGYKDLA-AIPKDK---------------------AILDIERPDLMIYEPHF 68
ML334211a                  VVKIDKQLFGELQKKFSKPKEKMSSLRSKYKNRKKTANKWVPVQVVPSQVNHTSLYQVDI 1200
                           *.*:*:*::*  :   : **:*                      ::  :  .  :*: .:

sp|Q08495|DEMA_HUMAN       TYSLLEHVELP----R----SRERSLSPKSTSPPPSPEVWADSRSPGIISQASAPRTTGT 120
ML334211a                  SYDEVNPVDVEVRIPDLTVDSRRRPVSP-----GPESK------YLGSLENVADPEDSHD 1249
                           :*. :: *::          **.* :**      *. :        * :.:.: *. :

sp|Q08495|DEMA_HUMAN       PRTSLPHFHHPET--------SRPDSNIYKKPPIYKQRESVGGSPQTKHLIEDLIIESSK 172
ML334211a                  ----CECKTHPGKLTRQSRNWVSQDSGLYDGTL-----KPVSARK-HTEMFQP------T 1293
                                    ** .           **.:*.        : *..    ..:::       .

sp|Q08495|DEMA_HUMAN       FPAAQPPDPNQPAKIETDYWPCPPSL-A--VVETEW-------------RKRKASRRGAE 216
ML334211a                  WKR-NSPDPVEERPRSTSP-YCQPFSEAAFGVNGSYIDNHVPASVRHPSRARNLSSDSNR 1351
                           :   : *** :    .*.   * *   *   *: .:             * *: *  . .

sp|Q08495|DEMA_HUMAN       EEEEEEDDDSGEEMKALRERQREELSKVTSNLGKMILKEEMEKSLPIR------------ 264
ML334211a                  -ENRIVQNSIGSQTNT----HVDRLARSGS---AASLAVVMASTFPNGGDPNFDLAGTAA 1403
                            *:.  ::. *.: ::     : :.*::  *       *   * .::*

sp|Q08495|DEMA_HUMAN       ---RKTRSLP----DRTPFHTSLHQGTSKSSSLPAYGRT-----TLSRLQSTEFSPSGSE 312
ML334211a                  KISQVCKSLSSSQTDITSCGVNLTRSRSMNSGMDVIHSTEVKRPTRPIYFDRNYSPSSAM 1463
                              :  :**     * *   ..* :. * .*.: .   *     *     . ::***.:

sp|Q08495|DEMA_HUMAN       TGSPGLQNGEGQRGRMDRG-NS-------LPCVLEQKIYPYEMLVVTNKGRTKLPPGVDR 364
ML334211a                  RIRTTPENRVGQPVKAKKATTSYREILKIQEKIKDANIISYQDL--VRCPRNKLPPHVDK 1521
                                 :*  **  : .:. .*          : : :*  *: *  ..  *.**** **:

sp|Q08495|DEMA_HUMAN       MRLERHLSAEDFSRVFAMSPEEFGKLALWKRNELKKKASLF         405
ML334211a                  LQLQKYLSDTEFVDVFKMPREEFDKMAPWKQANLKKNVSLY         1562
                           ::*:::**  :*  ** *  ***.*:* **: :***:.**:
```

##### Two pore domain $K^+$ channel (IPR003280) (Reciprocal Blast eval $2.65 \times 10^{-31}$)

```
sp|Q9NPC2|KCNK9_HUMAN      ---MKRQNVRTLSLIVCTFTYLLVGAAVFDALESDHEMREEEKLKAEEIRIKGKYNISS- 56
ML17562a                   MSKLGGKVIAAFATFIAVLTYLAIGAAIFQAIEQEHEIEQREEYDQQLNLLLDKVDPVFH 60
                              :  : : ::: ::..:*** :***:*:*:*.:**:.:.*: . :   : .* :

sp|Q9NPC2|KCNK9_HUMAN      ED-YRQLELVILQSEPHRAGVQWKFAGSFYFAITVITTIGYGHAAPGTDAGKAFCMFYAV 115
ML17562a                   DELKAPCHMIEILEANPNNTAQWDFVNSVIFCLTIVTTIGYGATYPVTDEGRGFCIFFAL 120
                           ::     .:: : .   .  .**.*..*. *.:*::****** : * ** *:.**:*:*:

sp|Q9NPC2|KCNK9_HUMAN      LGIPLTLVMFQSLGERMNTFVRYLLKRIKKCCGMRNTDVS-MENMVTVGFFSCMGTLCIG 174
ML17562a                   IGIPLFMACLAVYAQHTANLIRWIMKKLGLTNGLEDEQTEKMQQLITAAF--CLTVILTF 178
                           :**** :. :   .::  .::*:::*::    *:.: :.. *::::*..*  *: .:

sp|Q9NPC2|KCNK9_HUMAN      AAAFSQCEEWSFFHAYYYCFITLTTIGFGDYVALQTKGALQKKPLYVAFSFMYILVGLTV 234
ML17562a                   SGYLSYKENWDYSDSVYFTFISFTTIGFGDLYPSSNTTDFH------IIFICFLILGLIS 232
                           :. :*  *:*.: .: *: **::*******    ...  ::       : : ::::**

sp|Q9NPC2|KCNK9_HUMAN      IGAFLNLVVLRFLTMNSEDERRDAEERASLAGNRNSMVIHIPEEPRPSRPRYKADVPDLQ 294
ML17562a                   LGTVIEAQTT----------------------------------------NFGNLLRTIS 252
                           :*:.::  .                                         .:   :  :.

sp|Q9NPC2|KCNK9_HUMAN      SVCSCTCYRSQDYGGRSVAPQNSFSAKLAPHYFHSISYKIEEISPSTLKNSLFPSPISSI 354
ML17562a                   SSCDCLCKNR-CFGGDRVPVDE-------------------------------------- 273
                           * *.* *  .   :**  *  ::

sp|Q9NPC2|KCNK9_HUMAN      SPGLHSFTDHQRLMKRRKSV        374           100%
ML17562a                   --------------------        273       25%
```

##### Innexin (IPR000990) (Reciprocal Blast eval $1.10 \times 10^{-15}$, $1.18 \times 10^{-14}$)

```
sp|P33085|SHAKB_DROME      ML--D-----------IFRGLKNLVKVSHVKTDSIVFRLHYSITVMILMSFSLIITTRQY 47
ML036514a                  MLLLG-----------SLGTIKNLSIFKDLSLDDWLDQMNRTFMFLLLCFMGTIVAVSQY 49
ML32831a                   MRLSEKSTSHDCKACITRSHNEDCARRWGITIDDGWDQLNRSFMFGLLVVMGTTVTVRQY 60
                           *                    ::      :. *.   ::: :: . :*  :.  ::. **

sp|P33085|SHAKB_DROME      VGNPIDCVHTKDIPEDVLNTYCWIQSTYTLKSLFLKKQGVSVPYPGIGNSD--------- 98
ML036514a                  TGKNISCDGFTKFGEDFSQDYCWTQGLYTIKEAYDL-PESQIPYPGIIPENVPACREHAL 108
ML32831a                   TGSVISCDGFKKFGSTFAEDYCWTQGQYTVLEGYDQ-PNQNIPCPVPRPPSRRGSTLNTM 119
                           .*. *.*   ..: . . : *** *. **: . :      .:* *     .

sp|P33085|SHAKB_DROME      -------------GDPADKKHYKYYQWVCFCLFFQAILFYTPRWLWKSWEGGKIHALIMD 145
ML036514a                  KNGGKIVCPPEDQVKPLTRARHLWYQWIPFYFWVIAPVFYLPYMFVKRMGLDRMKPLLKI 168
ML32831a                   SQTQGFLHNPVE------------------------------------------------ 131

sp|P33085|SHAKB_DROME      L-DIGICSE-AEKKQKKKL--------LLDY-----LWENLRYHNWWAYRYYVCELLALI 190
ML036514a                  MSDYYHCTTETPSEEIIVKCADWVYNSIVDRLSEGSSWTSWRNRHGLGLAVLVSKFMYLG 228
```

```
ML32831a                   --------SDQELKKMTDKAATWLFYKFDLYMSEQSLLASLTNKHGLGLSVVFVKILYAA 183
                                        ::            :           .   ::  .    . :::

sp|P33085|SHAKB_DROME      NVIGQ-MFLMNRFFDGEFITFGLKVIDYMETDQEDRMDPMIYIFPRMTKCTFFKYGSSGE 249
ML036514a                  GSVLVMMMTTLMFQVGDFKTYGIEWLRQFPNPENYSTSVKHKLFPKMVACEIKRWGTTGL 288
ML32831a                   VSFGCFLLTADMFSIGDFKTYGSEWINKLKLEDNLATEEKDKLFPKMVACEVKRWGASGI 243
                             .   ::    *  *:* *:* : :  :   ::   .    :**:*. * . ::*::*

sp|P33085|SHAKB_DROME      VEKHDAICILPLNVVNEKIYIFLWFWFILLTFLTLLTLIYRVVIIFSPRMRV-YLFRMRF 308
ML036514a                  -EEENGMCVLAPNVIYQYIFLIMWFALAITICTNFGNIFFYLFKLTATRYTYNKLVATGH 347
ML32831a                   -EEEQGMCVLAPNVINQYLFLILWFCLVFVMFCNIVSIFASLIKLLFTYGSYRRLLST-A 301
                            *:.:.:*:*  **: : :::::** : :    .: .::  :. :         *.

sp|P33085|SHAKB_DROME      RLVRRDAIEIIVRRSKMGDWFLLYLLGENIDTVIFRDVVQDLANRLGHNQHHRVPGLKGE 368
ML036514a                  FSHKHPGWKFMYYRIGTSGRVLLNIVAQNTNPIIFGAIMEKLTPSVIKH--LRIGHVPGE 405
ML32831a                   FLRDDSAIKHMYFNVGSSGRLILHVLANNTAPRVFEDILLTLAPKLIQR--KLRAKDYD- 358
                                 . : :  .   .. .:* ::.:*    :*  ::  *:  : :.         .

sp|P33085|SHAKB_DROME      IQDA--          372    100%
ML036514a                  YLTDPA          411     21%
ML32831a                   ------          358     20%
```

**Munc13 (Protein Unc-13 (IPR027080))** (Reciprocal Blast eval 2.11 × $10^{-174}$)

```
sp|O14795|UN13B_HUMAN      ------------------------------------------------------------ 0
ML24335a                   MGLLYASYAPLNCFGHGNSPQYNKSLSQNLIRLIKHIGSEIALDIIYKKIAKVALDVPQT 60


sp|O14795|UN13B_HUMAN      ------------------------------------------------------------ 0
ML24335a                   ESSLNVYKEMKWLPLHLTRQLHLSNYMFRIIHDDCPTNFMNKFSFIIDNERVQNRRIIAN 120


sp|O14795|UN13B_HUMAN      ------------------------------------------------------------ 0
ML24335a                   EFNKYFASIASNLNEVYSSSDQVRISSLPSFTDYLPKSESSSIYLKESDYDEVSGIIGDL 180


sp|O14795|UN13B_HUMAN      ------------------------------------------------------------ 0
ML24335a                   KNGKSSDIPIHVIKKSSNVIAPFLSKFFNECMSSGHFPDELKTGRISPIYKKENEQLLEN 240


sp|O14795|UN13B_HUMAN      ----------------------MSLLCV-RV---KRAKF--------------------- 13
ML24335a                   YRPVSTLPVFGKILEKLIYTRLYSFLIAKGIIHENQYGFRKGHSTSHALNYSVQHIESMT 300
                                                  *:* .  :   ::  *

sp|O14795|UN13B_HUMAN      ----------QGSPDKFNTYVTLKVQNVKSTTVAVRGDQPSWEQDFMFEI----SRLDLG 59
ML24335a                   KNKQHVLGIFIDLSKAFDTIDHRKL-ITKLNNYGIRGNALKLIKSYLSNRTQFVSVLDIE 359
                                      .  . *:*    *:  .* .. .:**:  .  :.:: :     * **:

sp|O14795|UN13B_HUMAN      LSVEVWNKGLIWDTMVGTVWIALKTIRQSDEEGPGEWSTLEAET---------------- 103
ML24335a                   SGQLPVHFGVPQGSVLGPLLFVLYINDICNITKKGKFVLFADDTNIFVAADSKQKAYNIA 419
                            .    : *:  .:::* : :.*     .:    *::  :  :*

sp|O14795|UN13B_HUMAN      ----------LMKDDEICGTRNPTPHK------------------ILLDTRFELPFDIPE 135
ML24335a                   NEVLLAVISSRFYNKTLCFSEVPMTVTHVIVELWEKKTFHSVNSSVLQLELIDIPFKNKK 479
                                      : :. :* :. *   .                  :*    :::**.  :

sp|O14795|UN13B_HUMAN      -EEARYWTYKWEQINALGADNEYSSQEESQRKPLPTAAAQCSFEDPDSAVDDRDSDYRSE 194
ML24335a                   FRTFGCWAYSISN--TVDSD---------------------------SGCEDNEL--FL- 507
                            .    *:*. .:  ::.:*                           *. :*.:

sp|O14795|UN13B_HUMAN      TSNSFPPP-YHTASQPNASVHQFPVPVRSPQQLLLQGSSRDSCNDSMQSYDLDYPERRAI 253
ML24335a                   QNIVFAPSCYTSYVSRDCENNKLVVL------LVISDVSKE----E-----LALLEQKM- 551
                            .  * *  * :  . :.. ::: *       *::.. *::    .     *   *::

sp|O14795|UN13B_HUMAN      SPTSSSRYGSSCNVSQGSSQLSELDQYHEQDDDHRETDSIHSCHSSHSLSRDGQAGFGEQ 313
ML24335a                   -NVLSKVQEDECLLFNGE----------------------SFCL------KDGN----ST 578
                             . *.   ..* : :*.                        *       :**:    .

sp|O14795|UN13B_HUMAN      EKPLEVTGQAEK----------EAACEPKE-MKEDAT--TH---------PPPDLVLQKD 351
ML24335a                   GDPFPLAGNGDVFTGIDLVIHLGAICIAPQPERNDSFMTIDSGSDFRDAVTNETGMMAQE 638
                            .*: ::*:.:            * *   :  ::*:    .              :: ::

sp|O14795|UN13B_HUMAN      HFLGPQESFPEENA--SSPFTQARAHWIRAVTKVRLQLQEIPDDGDPSLPQWLPEGPAGG 409
ML24335a                   HFKSLIKTFDSSNFKITAPITPITPSPRTASQNGELDMISMS-MSHRSRQSSMSQNSADG 697
                           ** .  ::* ..*   ::*:*        *  : .*:: .:   .. *  . : :. *.*

sp|O14795|UN13B_HUMAN      LYGIDSMPDLRRKKPLPLVSDLSLVQSRKAGITSAMATRTSLKDEELKSHVYKKTLQALI 469
ML24335a                   VYSVSSQPLHPRISNLIH-SDGSVTTRQQSRSTSQ------------------------- 731
                           :*.:.* *   * . *   ** *:.  :::  **

sp|O14795|UN13B_HUMAN      YPISCTTPHNFEVWTATTPTYCYECEGLLWGIARQGMRCS-------------ECGVKCH 516
ML24335a                   FSLSMSDDHEAE-------TEEVSKEALMWMSAVQRIRKNIYMRNVENTAKEGEEGEEGE 784
                           : :* :  *: *       *   . *.*:*  * * :* .             * * : .
```

```
sp|O14795|UN13B_HUMAN      EKCQDLLNADC----------LQRAAEKSC-----KHGAEDRTQNIIMAMKDRMKIRERN 561
ML24335a                   GKGEELKENGAYPTADGEVGGLTRHENNSTVNSDNSSDMFDSDKVKSTAARQRMKVTENA 844
                            * ::* : ..           * *  ::*       . .  *  :    * ::***: *.

sp|O14795|UN13B_HUMAN      KPEIFEVIRDVFTVNKAAHVQQMKTVKQSVLDGTSKWSAKITITVVCAQGLQAKDKTGSS 621
ML24335a                   SKDMFELLRLAFGEESEKLDSVKKEVKHSILTGSSKWSAKIDIEVVEAQGLTGKDKSGTS 904
                           . ::**::* .*  :.    .  * **:*:* *:******* * ** **** .***:*:*

sp|O14795|UN13B_HUMAN      DPYVTVQVSKTKKRTKTIFGNLNPVWEEKFHFECHNSSDRIKVRVWDEDDDIKSRVKQRL 681
ML24335a                   DPYVTVQVGKIKKTTATIQQDLNPKWNENFSFDCNNSSDRIKVRVWDEDDDFKSRVMTHL 964
                           ********.* ** * **  :*** *:*:* *:*:****************:****  :*

sp|O14795|UN13B_HUMAN      KRESDDFLGQTIIEVRTLSGEMDVWYNLEKRTDKSAVSGAIRLQISVEIKGEE-KVAPYH 740
ML24335a                   KREADDFLGQAIIDVKQLCGETDVWLDLKQRTDRSDVSGQVHLRMTIKIEGEEQNMASYH 1024
                           ***:******:**:*: *.** *** :*::***:* *** ::*::::*:*** ::* **

sp|O14795|UN13B_HUMAN      VQYTCLHENLFHYLTDIQGS----GGVRIPEARGDDAWKVYFDETAQEIVDEFAMRYGIE 796
ML24335a                   IQYQLLHETLFNHLCEVNSGVNIPPGTRKLKGDELDEWPTFFEGPAQYIVEEFATRFGIE 1084
                           :**  ***.**::* :::..     *.*  :.   * * .:*:  ** **:*** *:***

sp|O14795|UN13B_HUMAN      SIYQAMTHFACLSSKYMCPG--VPAVMSTLLANINAYYAHTTASTNVSASDRFAASNFGK 854
ML24335a                   SIFMAMTQFSCLTQRFITKKDAPPQDVSKLLALINSHYRRN-------NGQEINVSNFGK 1137
                           **: ***:*:**:.:::      *  :*.*** **::* :.         .:.: .*****

sp|O14795|UN13B_HUMAN      ERFVKLLDQLHNSLRIDLSTYRNNFPAGS-PERLQDLKSTVDLLTSITFFRMKVQELQSP 913
ML24335a                   QKFTTVLDRLQSSLRENLASYKAVYPSNRGQEKLDDLENSIALLTSIVFFRLK------- 1190
                           ::*..:**:*:.*** :*::*:  *:.   *:*:**:.:: *****.***:*

sp|O14795|UN13B_HUMAN      PRASQVVKDCVKACLNSTYEYIFNNCHDLYSRQYQLKQELPPEEQGPSIRNLDFWPKLIT 973
ML24335a                   ------------LCLENTYDFISTKTDEVYAKYA-------NAEENTEHAKIAYWKKLIT 1231
                                       **:.**::* .: .::*::           *:. .  :: :* ****

sp|O14795|UN13B_HUMAN      LIVSIIEEDKNSYTPVLNQFPQELNVGKVSAEVMWHLFAQDMKYALEEHEKDHLCK--SA 1031
ML24335a                   LILVEMKEDKMFYEPIFNRFFPH---AALSAQTFWELLSRDLKKDLTENLDTWLSEIAAS 1288
                           **:  ::***  * *::*:*  .    . :**:.:*.*::*:*  * *: .  *.:  ::

sp|O14795|UN13B_HUMAN      DYMNLHFKVKWLHNEYVRDLPVLQGQVPEYPAWFEQFVLQWLDENED-VSLEFLRGALER 1090
ML24335a                   DVMGFQMTVKFVYDKKISHIPEYENSLPDYPKWFEPFVMKHLQESDSKIKNSFLIKSLDK 1348
                           * *.::.**::::: : .:*  :..:*:** *** **:: *:*.:. :. .**  :*::

sp|O14795|UN13B_HUMAN      DKKDGFQQTSEHALFSCSVVDVFTQLNQSFEIIRKLECPDPSILAHYMRRFAKTIGKVLM 1150
ML24335a                   DTFT--VQPNSSIKHSSSVIDIFSSLRATYDSLDKMKCPIPALRDKYHNRFCETIDAVLI 1406
                           *.     * ..   .*.**:*:*:.*. ::: : *::** *::  :* .**.:**. **:

sp|O14795|UN13B_HUMAN      QYADILSKDFP---AYCTKEKLPCILMNNVQQLRVQLEKMFEAMGGKELDLEAADSLKEL 1207
ML24335a                   EYTKRIIEVFKKGLGTLTSETTSCVILCNIHRVREELESIFKTMGGELLDITAKQTLEGT 1466
                           :*:. : : *     .  *.*.  *::: *:::* :**.:*::***: **: * ::*:

sp|O14795|UN13B_HUMAN      QVKLNTVLDELSMVFGNSFQVRIDECVRQMADILGQVRGTGNASPDARASAAQDADSVLR 1267
ML24335a                   QKRLKDARNSVISDLVKDMKPQIEKCIRDIKSDLRNVRS----------KNGEDYTEIVS 1516
                           * :*: . :.:   : :.:: :*::*:*:: . * :**.           . .:*  .::

sp|O14795|UN13B_HUMAN      PLMDFLDGNLTLFATVCEKTVLKRVLKELWRVVMNTMERMIVLPPLTDQTGTQLIFTAAK 1327
ML24335a                   PLIEYFDVVFKIFVDNLYDDVRKPLLQQTWKRTLKLFEEIIILPDINAIA---------- 1566
                           **:::*  :.:*.     . * * :*:: *: .:: :*.:*:** :.  :

sp|O14795|UN13B_HUMAN      ELSHLSKLKDHMVREETRNLTPKQCAVLDLALDTIKQYFHA-GGNGLKKTFLEKSPDLQS 1386
ML24335a                   -------------HDEVQELNSKQSQVVEAMLDILKVYFTDSSGKCLKAKNLEKTTEMRD 1613
                                        ::*.::*. **. *:: ** :* **   .*: ** . ***: :::.

sp|O14795|UN13B_HUMAN      LRYALSLYTQTTDTLIKTFVRSQTTQG-SGVDDPVGEVSIQVDLFTHPGTGEHKVTVKVV 1445
ML24335a                   LRKVLSQYRLTTDTLIKNFVTQADNQNRFAEEDSQGEIQLQVDLFTPPGQEHHDCTVKGL 1673
                           ** .** *  *******.** .  .*.  . :*  **:.:****** **  .*. *** :

sp|O14795|UN13B_HUMAN      AAN----DLKWQTAGMFRPFVEVTMVGPHQSDKKRKFTTKSKSNNWAPKYNETFHFLLGN 1501
ML24335a                   FSFYYSKFLPWCLELSHS------VSVP-------------------------------- 1695
                            :      * *     .       :  *

sp|O14795|UN13B_HUMAN      EEGPESYELQICVKDYCFAREDRVLGLAVMPLRDVTAKGSCACWCPLGRKIHMDETGLTI 1561
ML24335a                   ------------------------------------------------------------ 1695


sp|O14795|UN13B_HUMAN      LRILSQRSNDEVAREFVKLKSESRSTEEGS          1591   100%
ML24335a                   ------------------------------          1695    33%
```

#### Munc18 (Sec1-like protein (IPR001619)) (Reciprocal Blast eval $2.59 \times 10^{-22}$)

```
sp|P61764|STXB1_HUMAN      -MAPIGLKAVVGEKIMHDVIKKVKKKGEWKVLVVDQLSMRMLSSCCKMTDIMTEGITIVE 59
ML005115a                  MAADVRTRQIDAVKHM-LNLNSPKSESTWKILVFDEYGRDIIAPLLTVKELRECGVTLNL 59
                             * :  : : . * *   ::. *.:. **:**.*: .  :::   .:.::   *:*:

sp|P61764|STXB1_HUMAN      DINKRREPLPSLEAVYLITPSEKSVHSLISDFKDPPTAKYRAAHVFFTDSCPDALFNELV 119
ML005115a                  LIADKRDPIPDVPVVYFVMPTKQNIDLISQDCKNQMYEKFY---INFITAVSRQLLEDLA 116
                            * .:*:*:*.: .**:: *:.:. :*::.*: : .* *:    *:    : *  :    *:::*.

sp|P61764|STXB1_HUMAN      KS----RAAKVIKTLTEINIAFLPYESQVYSLDSADSF-QSFYSPHKAQMKNP----ILE 170
```

```
ML005115a                  QATIESDSVARVEKIFDQYLSFCMLEQNLFTVSAYDAQSCSYYALNRPDAKDADIEKCIN 176
                           ::     :.  ::.: :  ::*   *.::::::.: *:  *:*: :: : *:      ::

sp|P61764|STXB1_HUMAN      RLAEQIATLCATLKEYPAVRYRGEYKDNALLAQLIQDKLDAYKADDPTM-----GEGPDK 225
ML005115a                  AITDSVFSLFVTLRELPVIRCPKGNAAEIVAER-LEKKMRDAR-RDPKSELFKDQAITSF 234
                            :::.: :* .**:* *.:*       : :  : ::.*:   :  **.          .

sp|P61764|STXB1_HUMAN      ARSQLLILDRGFDPSSPVLHELTFQAMSYDLLPIENDVYKYETSGIGEARVKEVLLDEDD 285
ML005115a                  QRPLLILLDRNMDICSTLHHTWTYQALIHDLFNLYLNRCKV----EVDEKVKNYDLGPDD 290
                            *  *::***.:* .* : *  *:**: :**: :  :  *       : :**:  *. **

sp|P61764|STXB1_HUMAN      DLWIALRHKHIAEVSQEVTRSLKDFSSSKRMN---------TGEKTTMRDLSQMLKKMPQ 336
ML005115a                  QFWAKNKGKPFPEVADAVHVELEACRAAEQEIKRLKDVMGESGVENANDQIANTVSSLPE 350
                           ::*   : * : **:: *  .*:   ::::           :* :.:  :::: :..:*:

sp|P61764|STXB1_HUMAN      YQKELSKYSTHLHLA---EDCMKHYQGTVDKLCRVEQDLAMGTDAEGEKIKDPMRAIVPI 393
ML005115a                  LMERKKRVDMHINIATALSDCIRA--RNLDKFFEVEEKIMTKTTLEES--------VLDT 400
                             :. .: . *:::*   .**::    .:**: .**:.:   *  * .        ::

sp|P61764|STXB1_HUMAN      LLDAN-VSTYDKIRIILLYIFLKNGITEENLNKLIQHAQIPPEDSEIITNMAH------L 446
ML005115a                  LKNPSVGTAEDKLRLFIIYYLTTSDLPDNELQQYIEALQEAGCDTSCIEYLAKLKSVSSL 460
                           * : .  :: **:*::::* : ...: :::*:: *:  *    *:. *  :*:      *

sp|P61764|STXB1_HUMAN      GVPIVTDSTLRRRSKPERKERISEQTYQLSRWTPIIKDI-MEDTIEDKLDTKHYPYISTR 505
ML005115a                  NVPAQTKLTSSSGSYS----SMS----NMFKYVKQSSKFMMEGVKSLVVGHRKLPVTRTV 512
                           .**  *. *    *       :*    :: ::.   ..: **.. .  :. :: *   *

sp|P61764|STXB1_HUMAN      ----------SSASFSTTAVSARYGHWHKNKAPGEYRSGPRLIIFILGGVSLNEMRCAYE 555
ML005115a                  DCLSELKQTPETDSFL--ALDPKIYRKTENLLPALKNTFNDVYVFVVGGGSYVEYQNLMT 570
                                     .: **   *:. :  :  :*  *.  .:   : :*::** *  * :

sp|P61764|STXB1_HUMAN      VTQANGKWEVLIGSTHILTPQKLLDTLKKLNKTDEEISS           594   100%
ML005115a                  YSEQHPGKRIVYGSSEMMDSKNFLDQLTKLGSSSS----           605    21%
                            :: :   .:: **:.::  :::** *.**..:..
```

#### Complexin (Synaphin (IPR008849))

Complexin was identified de-novo using O14810|CPLX1_HUMAN as a query (BlastP evalue =$2.64 \times 10^{-04}$ )

```
sp|O14810|CPLX1_HUMAN      MEFVMKQALGGATKDMGKMLGGD--------EEKDPDAAK--KEEERQEALRQAEEERKA 50
ML08643a                   MNFIAKQVLKSKTGALTDKLDEVSKKCKELLESDEPDAAPQAQNLSEEEQMKQEADRRKE 60
                           *:*: **.* . *  : . *.          *..:****   :: ..:* ::*  :.**

sp|O14810|CPLX1_HUMAN      KYAKMEAEREAVRQGIRDKYGIKKKEEREAEAQAAMEANSEGSLTRPKKAIPPGCGDEV- 109
ML08643a                   HYNAFNAEREEQRNKMREKYGLRSDSCPDKSTGNE-----SGADNCVNEASEPESKEDHK 115
                           :*  ::****  *: :*:***::...  : .:        .*: .  ::*  * . ::

sp|O14810|CPLX1_HUMAN      ----EEEDESILDTVIKYLPGPLQDMLKK          134    100%
ML08643a                   DSEKKDGDCCVM-----------------          127     27%
                               :: * .::
```

#### ABLIM3 (reciprocal eval 0.00)

##### Villin_headpiece domain

```
sp|O94929|ABLM3_HUMAN      MNTSIPYQQNPYNPRGSSNVIQCYRCGDTCKGEVVRVHNNHFHIRCFTCQVCGCGLAQSG 60
ML435828a                  ------------------------------------------------------------ 0


sp|O94929|ABLM3_HUMAN      FFFKNQEYICTQDYQQLYGTRCDSCRDFITGEVISALGRTYHPKCFVCSLCRKPFPIGDK 120
ML435828a                  ---------------------------------MSAVE-------------VVPPPLMED 14
                                                            :**:                *  *: :.

sp|O94929|ABLM3_HUMAN      --VTFSGKECVCQTCSQSMASSKPIKIRGPSHCAGCKEE--IKHGQSLLALDKQWHVSCF 176
ML435828a                  CAVPPLGEAC----CDPGLPDP------CPPECQQPPEQASLLAGFYALSLQGAYGTTLI 64
                             *   *: *    *. .: .        * .*    *:  :  *   *:*:  : .: :

sp|O94929|ABLM3_HUMAN      KCQTCSVILTGEYISKDGVPYCESDYH-AQFGIKCE-TCDRYISGRVLEAGGKHYHPTCA 234
ML435828a                  RLSN----------DQGTLKFLVPQLSEQAVQTECEEECEEKKE----EAAEEE---AAA 107
                           : ..          .:. : :   :     .  :**  *:.  .    **. :.   :.*

sp|O94929|ABLM3_HUMAN      RCVRCHQMFTEGEEMYLTGSEVWHPICKQAARAEKKLKHRRTSETSISPPGSSIGSPNRV 294
ML435828a                  EEGKGAESLADNRDLDSISGELSTPLCDDSDL---------------PPP---------A 143
                           .  :  : :::..::   ..*:  *:*.::                  **         .

sp|O94929|ABLM3_HUMAN      ICAKVDNEILNYKDLAALPKVKSIYEVQRPDLISYEPHSRYMSDEMLERCGYGESLGTLS 354
ML435828a                  ECDEERDRM------PDPPQI--------------------------------------- 158
                            * :  :.:         *::

sp|O94929|ABLM3_HUMAN      PYSQDIYENLDLRQRRASSPGYIDSPTYSRQGMSPTFSRSPHHYYRSGPESGRSSPYHSQ 414
ML435828a                  -----------------EEPGDVQKPVGRKESVA----------FD--P----------- 178
                                            ..** ::.*.  ::.::          :   *

sp|O94929|ABLM3_HUMAN      LDVRSSTPTSYQAPKHFHIPAGDSNIYRKPPIYKRHGDLSTATKSKTSEDIS----QTSK 470
ML435828a                  -------ENNPNAVPNFHTPKAV----------NKFGNYTQVQM--LGEDLHAHPSTRRP 219
                                   .. :*  :** * .           ::.*: : .     .**:
```

```
sp|O94929|ABLM3_HUMAN      YSPIYSPDPYYASESEYWTYHGSPKVPRARRFSSGGEEDDFDRSMHKLQSGIGRLILKEE 530
ML435828a                  MSPV-------VSQPIRWECNRSQKGSSPPRITKGTPNKNTSPVRHRTPS---------- 262
                            **:       .*:   *  : * *     *::.*  :.: .   *:  *

sp|O94929|ABLM3_HUMAN      MKARSSSYADPWTPPRSSTSSREALHTAGYE------MSLNGSPRSHYLADSDPLISK-- 582
ML435828a                  ----PQRFKSPDRGPRVRVSNGL-PAGTAYKASGEIDPKIKESPPKDIVNKIRNTIESPS 317
                                . : .*   **  .*.      :.*:       .:: ** .. : .    *..

sp|O94929|ABLM3_HUMAN      -SASLPAYRRNGLHRTPS-ADLFHYDSMNAVNWGMREYKIYPYELLLVT-----TRGRNR 635
ML435828a                  RVKSPPKP------RSPSPVKPFDTDAFRLARKLAQEVADCPPDCIMVRVKDLQRCPKAL 371
                              * *        *:** .. *. *::. ..   :*    * : ::*         :

sp|O94929|ABLM3_HUMAN      LPKDVDRTRLERHLSQEEFYQVFGMTISEFDRLALWKRNELKKQARLF  683
ML435828a                  LPPELDKARLEEYISDTEFRRTLGLSRSEYESFSEWKQQEIKKELGLF  419
                           ** ::*::***.::*: ** :.:*:: **:: :: **::*:**:  **
```

#### **Contactin-5** (reciprocal eval $1.19 \times 10^{-17}$)

```
sp|O94779|CNTN5_HUMAN      ---------------------------MASSWKLM----------------------LFL 11
ML087115a                  MNSTESMPCKEAQSLFSAKNSVDIFQDEDGTFTLLTSIHTANVARRDSQPFCTGDLSLAN 60
                                                        .::.*:                      *

sp|O94779|CNTN5_HUMAN      SVTMCLSEYSKSLPGLSTSYAALLRIKKSSSSSLFGSKTRPRYSSPSLGTLSASS----P 67
ML087115a                  DASICIVTC--SLPDYE-------------CPREFKQRCACPWDKPILKDIDKGICVKSI 105
                           ..::*:     ***. .             .   * .:    :..* *  :. .

sp|O94779|CNTN5_HUMAN      SWLGAAQNYYSPINLYHSSDAFKQDESVDYGPVFVQEPDDIIFPTDSDEKKVALNCEVRG 127
ML087115a                  RDCPPYNTYILPIKGLK-------------QFNFTKQPEDLVK---KRGDKIRMQCTVNG 149
                                 :.*  **:  :                *.::*:*::    .  .*: ::* *.*

sp|O94779|CNTN5_HUMAN      NPVPSYRWLRNGTEI------DLESDYRYSLIDGTFIISNPSEAKDSGHYQCLATNTVGS 181
ML087115a                  DTNYQYFWYKSPSLAMGSHSVSRSSDKYLKQTGNPYFDINSVQPSDSGYYHCIVRTGDTV 209
                           :   .* * :. :        . .**   .  .. ::  *  : .***:*:*:. .

sp|O94779|CNTN5_HUMAN      -----------------ILSREATLQFAYLGN-FSGRTRSAVSVREGQGVVLMCSPPPHS 223
ML087115a                  AGVQRDPKYGSRDAKYSLVSNIGYLKVEYFDVPQLGETASIVTQKGTLTTVLPCDLPKSE 269
                                            ::*. . *:. *:.    *.* * *: :    .** *. *  .

sp|O94779|CNTN5_HUMAN      PEIIYSWVFNEFP-----SFV-AEDSRRFISQETGNLYISKVQTSDVGSYICLVKNTVTN 277
ML087115a                  PPVEAVWYKDDQIVSSKRMFVVPQAQKPDDNVKRGSLYIVMAETGDAGVYTCKTTYGDQT 329
                           * :   *  ::        **  : .:   . : *.***  .:*.*.* * * ..    .

sp|O94779|CNTN5_HUMAN      ARVLSPPTPLTLRNDGVMGEYEPKIEVHFPFTVTAAKGTTVKMECFALGNPVPTITWMKV 337
ML087115a                  YE----VAKVNLQVQGIIGQPTSDPEIVVQVAGRANVGETATFFCAGVGNPVPDVSWSKE 385
                            .     : :.*: :*::*:   . *: . .:  *  * *..: * .:***** ::* *

sp|O94779|CNTN5_HUMAN      NGYIPSK---ARLRKSQAVLEIPNVQLDDAGIYECRAENSRGKNSFRGQLQVYTYPHWVE 394
ML087115a                  DGTSIFESDRVALSDFNRKLTIADVRPEMSRDFACTVSSENSANSRKTILYLSIMPASYE 445
                           :*    :   . * . :  * * :*: : :  : * ...... ** :  * :   *   *

sp|O94779|CNTN5_HUMAN      -KLND------TQLDSGSPLRWECKATGK---PRPTYRWLKNGVPLSP--QSRV------ 436
ML087115a                  INFQETPVSHVIKPNVYTPFTIQCAITVDPVIEAPVMYWIKNGASLDMTPPTRIRTVTDN 505
                            ::::       : :  :*:  :*  * .     *.  *:***. *.    :*:

sp|O94779|CNTN5_HUMAN      -------EMVNGVLMIHNVNQSDAGMYQCLAENKYGAIYASAELKILASAPTFALNQLKK 489
ML087115a                  QKILNKVTIRTSRLIFDTPNESDKGVYQCIVENKEQMRQATAYVDVDKSPSSYDVFYDQE 565
                                   : .. *::.. *:** *:***:.***     *:* :.:  *  :: :   ::

sp|O94779|CNTN5_HUMAN      TIIVTKD---QEVVIECKPQGSPKPT---------------------------------- 512
ML087115a                  YEVFTEEKKIDEAAEHCTKKWGPRGNLVTITNKKESDQVFQLLVQAGVTRAWIGLTSESP 625
                             :.*::   :*.. .*. : .*: .

sp|O94779|CNTN5_HUMAN      ------ISWKKGDRAVRENKR------IAILPDGSLRILNASK-------SDEG--KYVC 551
ML087115a                  IEQTGIWKWQAGSHVRDDFSYWYSGNPDNALGNESYAIMDVSRAGHWLDVNNEGSYPFVC 685
                                  .*: *.:.  : .          * : *  *::.*:       .:**   :**

sp|O94779|CNTN5_HUMAN      RGENVFGSAEIIAS-LSVKEPTRIELTPKRTELTVGESIVLNCKAIHDASLDVTFYWTLK 610
ML087115a                  SRYRA--KCPSVAKFFNNSARPISGLETEGTEYEVGKEITVKCVEKEESNVSQKIVCHPS 743
                              ..  ..  :*. :. .      *  : **  **:.*.::*   .::.:. .:    .

sp|O94779|CNTN5_HUMAN      GQPIDFEEEGGHFESIRAQASSADLMIRNILLMHAGRYGCRVQTTADSVSDEAELLVRGP 670
ML087115a                  G----------TF--------------------MPSSIPCPVKSRAQHVTTSL------- 766
                           *           *                      .   * *:: *: *: .

sp|O94779|CNTN5_HUMAN      PGPPGIVIVEEITESTATLSWSPAADNHSPISSYNLQARSPFSLGWQTVKTVPEIITGDM 730
ML087115a                  ------------------------------------------------------------ 766


sp|O94779|CNTN5_HUMAN      ESAMAVDLNPWVEYEFRVVATNPIGTGDPSTPSRMIRTNEAVPKTAPTNVSGRSGRRHEL 790
ML087115a                  -CHVTVGLFLFLLYR--------------------------------------------- 780
                            . ::*.*  :: *.

sp|O94779|CNTN5_HUMAN      VIAWEPVSEEFQNGEGFGYIVAFRPNGTRGWKEKMVTSSEASKFIYRDESVPPLTPFEVK 850
ML087115a                  ------------------------------------------------------------ 780
```

```
sp|O94779|CNTN5_HUMAN      VGVYNNKGDGPFSQIVVICSAEGEPSAAPTDVKATSVSVSEILVAWKHIKESLGRPQGFE 910
ML087115a                  ------------------------------------------------------------ 780


sp|O94779|CNTN5_HUMAN      VGYWKDMEQEDTAETVKTRGNESFVILTGLEGNTLYHFTVRAYNGAGYGPPSSEVSATTK 970
ML087115a                  ------------------------------------------------------------ 780


sp|O94779|CNTN5_HUMAN      KSPPSQAPSNLRWEQQGSQVSLGWEPVIPLANESEVVGYKVFYRQEGHSNSQVIETQKLQ 1030
ML087115a                  ------------------------------------------------------------ 780


sp|O94779|CNTN5_HUMAN      AVVPLPDAGVYIIEVRAYSEGGDGTASSQIRVPSYSGGKITSAQSTLHSLSTSSSSVTLL 1090
ML087115a                  ------------------------------------------------------------ 780


sp|O94779|CNTN5_HUMAN      LALMIPSTSW      1100  100%
ML087115a                  ----------      780    22%
```

#### **Calcium-activated potassium channel** (Reciprocal Blast eval 0.00, 3.84 × 10-177)

```
sp|Q12791|KCMA1_HUMAN      MANGGGGGGGSSGGGGGGGGSSLRMSSNIHANHLSLDASSSSSSSSSSSSSSSSSSSSSS 60
ML04056a                   ------------------------------------------------------------ 0
ML128229a                  ------------------------------------------------------------ 0


sp|Q12791|KCMA1_HUMAN      VHEPKMDALIIPVTMEVPCDSRGQRMWWAFLASSMVTFFGGLFIILLWRTLKYLWTVCCH 120
ML04056a                   -------------------------------------------------------MSCCD 5
ML128229a                  ------------------------------------------------------------ 0


sp|Q12791|KCMA1_HUMAN      CGGKTKEAQ-----KINNGSSQADGTLKPVDEKEEAVAAEVGWMTSVKD-WAGVMISAQT 174
ML04056a                   EFVNIDISLQFKNRDLYENTSQISRRMFPNHYDRD-KLDEVELHNKLRHYIVNEVISLQT 64
ML128229a                  ---MRKRRRDQDSEEIYFGCIRFIRRVYNIQR-------------RV-VQFAEDISYPKT 43
                                .        .:  .  :    :   .               :    .  :   :*

sp|Q12791|KCMA1_HUMAN      LTGRVLVVLVFALSIGALVIYFIDSSNPIESCQNFYK--DFTLQIDMAFNVFFLLYFGLR 232
ML04056a                   NKGMVLAILYFLFNVISVTLYINQSDQPLEQCEDVLNTCNWTLYADIGLNVFFLFHFLLR 124
ML128229a                  KKGRAMTWVQLLMNLIAIGSFAAQTPEEIERCVELT---EITIIVDLICNIFFLWRFVVM 100
                            .* .:. : : :.: ::  :  :: : :* * :.    : *:  *:  *:***  * :

sp|Q12791|KCMA1_HUMAN      FIAANDKLWFWLEVNSVVDFFTVPPVFVSVYLNRSWLGLRFLRALRLIQFSEILQFLNIL 292
ML04056a                   LIAAPNRLKFWFQMGSIVDFCTIPPCALALILERNWLGMRFMRALRFMQMGEILMTFNST 184
ML128229a                  YIVSKDKLCFWVEIGSLIDFCTIPPALVTFGIRKSWLGLRFVRCLKLLSFGDDIAAVKLI 160
                            *.: ::* **.::.*::** *:**  ::. :.:.***:**:*.*:::.:.: :  .:

sp|Q12791|KCMA1_HUMAN      KTSNSIKLVNLLSIFISTWLTAAGFIHLVENSGDPWENFQ-NNQALTYWECVYLLMVTMS 351
ML04056a                   GTS-IVQLLVLMLDFVSVWLFAAGFVHLVENSGDPWKADSARKDNFTYFDSVWLTMVTMS 243
ML128229a                  SSHKMLTLSRTLCQFVSFWLTAAGLYFLAARMSD-------SGNKIHFFECMYFLLVTIS 213
                            :   : *   :  *:* ** ***: .*. . .*         : : :::.::: :**:*

sp|Q12791|KCMA1_HUMAN      TVGYGDVYAKTTLGRLFMVFFILGGLAMFASYVPEIIELIGNRKKYGGSYSAVSGRKHIV 411
ML04056a                   TVGYGDVYPTTGIGRSFMMVFIVCGLAMFATYTPYLYEFYRSRSPYAGRYERADSKKYVV 303
ML128229a                  TIGYGDLKPETTTTRLVVIVLIAVGLAVFASFGPAMIEMFKKSPPYTLDYDDSK-SSHVV 272
                           *:****:   *   * .::.:*  ***:**:: * : *:  .   *   *.  .  .::*

sp|Q12791|KCMA1_HUMAN      VCGHITLESVSNFLKDFLHKDRDDVNVEIVFLHNISPNLELEALFKRHFTQVEFYQGSVL 471
ML04056a                   VCGHITYNNVKSFLADFLHPDRGDRTSRVLFLDNKRPEPELTGLIKRYFLSVQYFEGSAL 363
ML128229a                  VCGYINTSTVSNFMRDFLHPENEDHETTAVFLNTIPPPEDLQNLMHRNFYRCQYYQGSPL 332
                           ***:*. ..*..*: **** :. *     :**..  *  :*  *::* *   ::::** *

sp|Q12791|KCMA1_HUMAN      NPHDLARVKIESADACLILANKYCADPDAEDASNIMRVISIKNYHPKIRIITQMLQYHNK 531
ML04056a                   NHNDLRRVQLEFSDAVILLCNKDSDDTKSEDAATILRAISVKNYCSKARLIMQLLNYKSK 423
ML128229a                  RASDLARVKLSTAKACLVIANKHSAEPDAEDAANILRVISIKNYKSDVRVIIQILNYHNE 392
                           .  ** **::. :.* :::.** . : .:***:.*:*.**:***  . *:* *:*:*:.:

sp|Q12791|KCMA1_HUMAN      AHLLNIPSWNWK-EGDDAICLAELKLGFIAQSCLAQGLSTMLANLFSMRSFIKI--EEDT 588
ML04056a                   SFLQTIPNWDPINYRDENICISEFMYGLVAQSCLCPGFSTLMINLISTRSYIPGINENFD 483
ML128229a                  EHLFNVPDWNHQLYGDEVICINESKD------------------------------VRSN 422
                            .* .:*.*:     *: **: *                                  .

sp|Q12791|KCMA1_HUMAN      WQKYYLEGVSNEMYTEYLSSAFVGLSFPTVCELCFVKLKLLMIAIEYKSANRESRILINP 648
ML04056a                   WKSDYAYGCDHEIYTVMISSDFNGMTFPEAAEICYKQLNLLMIAVENFDNS---GLAINP 540
ML128229a                  WEYEYGYGASNELYTGSLSSDFLDMTFPEVAEICLIKLHILLIGVETNS--SSTGILFNP 480
                           *:  *  * .:*:**  :** * .::** ..*:*  :*::*:*.:*  .      : :**

sp|Q12791|KCMA1_HUMAN      -GNHLKIQEGTLGFFIASDAKEVKRAFFYCKACHDDITDPKRIKKCGCKRPKMSIYKRMR 707
ML04056a                   LPDQFTIRKECLGYFIASSYKDVCRARFYCALCLNEGIEI-GFKPCKCRLEAVKKKRSNV 599
ML128229a                  SPHVFRFRQGTQGIFIAQNPTVVKSAALFCFKCHEYAIQS-EIVKCTCDKSDSGYTNSRT 539
                             . : :::   * ***.. . *  * ::*  * :   :   :  * *        .

sp|Q12791|KCMA1_HUMAN      RACCFDCGRSERDCSCMSGRVR-GNVDTLERAFPLSSVSVNDCSTSFRAFEDEQPSTLSP 766
ML04056a                   QLIPLSEHKSPN--NNESGLTRVGSIDSKS----STATDVNQLRTVRHLFSQHST----- 648
ML128229a                  ---------------------N------------------SNA------------NTTVP 548
                                                .                  *
```

```
sp|Q12791|KCMA1_HUMAN      KKKQRNGGMRNSPNTSPKLMRHDPLLIPGNDQIDNMDSNVKKYDSTGMFHWCAPKEIEKV 826
ML04056a                   -----------------KSTSVDRGTLA-------GAGMSSTFDPTGNYHWAPEKKFEDV 684
ML128229a                  KKKRKKQS-EYVPYW-KRLSQEER----------AKMDERLTLDGSGTFHWRESCLLEEV 596
                                            :    :              .   . * :* :**     :*.*

sp|Q12791|KCMA1_HUMAN      ILT------------------------RSEAAMTVLSGHVVVCIFGDVSSALIGLRNLVM 862
ML04056a                   DLAGYITKEYQYKPDDAEAINQRVVIKYINEKDHELNNHIVICAFGDDKTPISGLSSLIK 744
ML128229a                  TMT--------------------------KEDVATLTDHIIVCVFGDNNAPTVGLRNLLL 630
                            ::                          :     *..*:::* *** .:   ** .*:

sp|Q12791|KCMA1_HUMAN      PLRASNFHYHELKHIVFVGSIEYLKREWETLHNFPKVSILPGTPLSRADLRAVNINLCDM 922
ML04056a                   PLRASNLPYKKLKVVLLLAEKDFLLKAWKELRHFPKIRVLCGSALDRSKLKRAFINRASM 804
ML128229a                  PLRNSNLPTTEVKKVLLLGKKEYIEKEWKDIDEFQDVYFMEGSPIRREDVISAGVQNCLM 690
                           *** **:   ::* ::::.. ::: : *: : .* .: .: *: : * .:  . :: . *

sp|Q12791|KCMA1_HUMAN      CVILSANQNNI--DDTSLQDKECILASLNIKSMQFDDSIGVLQANSQGFTPPGMDRSSPD 980
ML04056a                   VVILSNYGKRMAYKDPFLLDKEPILACLNLPQVA---------------------GK--- 840
ML128229a                  CLILSSALQSK-ATDPNLVDKETILSFLQIKGCRGTESSD-------------------T 730
                            :***   :     *  * *** **: *::

sp|Q12791|KCMA1_HUMAN      NSPVHGMLRQPSITTGVNIPIITELVNDTNVQFLDQDDDDDPDTELYLTQPFACGTAFAV 1040
ML04056a                   ------------SENSKPVEFIAELVNDSNVHFLDQDNEDEPDIELHLTMKFAQGSCITA 888
ML128229a                  SNTTKYRLSEKPIRNRAQIPIISEIVNDSNVQFMEKSKDS-SVGALYMALPFTRGQAFIV 789
                                         .   : :*:*:***:**:*:::..:.     *:::  *: * .: .

sp|Q12791|KCMA1_HUMAN      SVLDSLMSATYFNDNILTLIRTLVTGGATPELEALI---AEENALRGGYSTPQTLANRDR 1097
ML04056a                   TALDTLMSEAYYNDEVMTLVRALIAGANVPEYDAE------DEEHSGPLQVPANALCGSQ 942
ML128229a                  SVLDAILSLTFYNEKLMSIIRTLITGIGNSHKNETADNYLPENSEDDVMYYQPWLRTRMS 849
                           :.**:::* :::*:::::::*:*::*    . :        ::   .

sp|Q12791|KCMA1_HUMAN      CRVAQLALLDGPFADLGDGGCYGDLFCKALKTYNMLCFGIYRLRDAHLSTPS------QC 1151
ML04056a                   CRVIRIKAEEPSLSEHIKGKAYKDVFVELLQKFNLLCLGLHRAIDWHLDVSV-------E 995
ML128229a                  CRIAQISLRDERFSRF---RVYGELFVYLLHNYSMLALGLYRSYQWANKGKSTKARDNHN 906
                           **: ::   :  ::       * ::*   *:.:.:*.:*::*  :   .

sp|Q12791|KCMA1_HUMAN      TKRYVITNPPYEFELVPTDLIFCLMQFDHNAGQSRASLSHSSHSS----------QSSSK 1201
ML04056a                   HKRFVYINPPGETKIVASDYLYVIVQTSKLQPPTEVMLNKIRDTMLRKGGTPEHEQSADS 1055
ML128229a                  EDRYVTTAPSYDAKLRPNDKVFVLAQSEMELPFPLEMGKNEMD-GLDDDGMSYQTSRETS 965
                            .*:*   *  : ::  .* :: : * .          .:  .            .   .

sp|Q12791|KCMA1_HUMAN      ---------------KSSSVHSIPSTANRQNRPKSRE-----SRDKQKYVQE----ERL- 1236
ML04056a                   KLTRQNKVDNESVRFESVSTRSVSSVAPSPRTPLS--------PHSPPAAGE----ERFY 1103
ML128229a                  PPTVMKPKPDVSVPIAVEEIQPVTQPNNQPSNPTTDTDTDHSSPLKEPVTMETKTDDTLS 1025
                                             . : : .       * :          .   . *    : :

sp|Q12791|KCMA1_HUMAN      -----------------------------------------------   1236  100%
ML04056a                   NPLHA--------------AANHTSSKTTQHILENFRNSYASYSDGH   1136   38%
ML128229a                  HPVDLPAVYTTDPTENLKPPNNTTTSNDIAHLIA-------------   1059   35%
```

***Supplementary figure 16. Alignments of M. leidyi and H. sapiens neuronal proteins shown in Table 1 (https://www.ebi.ac.uk/Tools/msa/clustalo/).***

#### Supplementary videos

***Supplementary video 1. ML02212a localization in the aboral organ of M. leidyi.*** *Neuropeptide: cyan, DAPI: yellow.*

***Supplementary video 2. ML07842a localization around the pharynx of M. leidyi.*** *Neuropeptide: cyan, DAPI: yellow, Tubulin: red.*

***Supplementary video 3. ML07842a localization in the aboral organ of M. leidyi.*** *Neuropetide: cyan, DAPI: yellow.*

***Supplementary video 4. ML07842a localization in the aboral organ of an adult M. leidyi.*** *Neuropeptide: cyan, DAPI: yellow.*

***Supplementary video 5. ML21545a localization around the pharynx of M. leidyi.*** *Neuropeptide: cyan, DAPI: yellow, tubulin: red.*

***Supplementary video 6. ML17711a localization around the pharynx of M.leidyi.*** *Neuropeptide: cyan, DAPI: yellow, tubulin: red.*

***Supplementary video 7. ML17711a localization in the aboral organ of M. leidyi.*** *Neuropeptide: cyan, DAPI: yellow.*

***Supplementary video 8. 3D reconstruction of a part of M. leidyi epidermis.*** *Colours coded as in Fig 6A.*

***Supplementary video 9. 3D reconstruction of a part of M. leidyi epidermis.*** *Neuron is shown half-transparent. Colours coded as in Fig 6B.*

***Supplementary video 10. 3D reconstruction of a part of M. leidyi epidermis.*** *Neuron is not shown. Colours coded as in Fig 6C.*

***Supplementary video 11. 3D reconstruction of a M. leidyi synapse.***

### Supplementary tables

#### *Supplementary table 1. Neuropeptide precursors predicted by NeuroPID.*

| ***Linear precursors*** |
| --- |
| ***-Enriched in uncharacterised metacells*** |
| >ML043317a |
| MKTILLISCLLSAVYSRVLRLNEEDWSRNGAMEDSEDWSRNGAMRDSEDWNRNGGMRDSEDWSRNGRMEDSEDWSRNGAMEDSEDWSRNGAMED |
| SEEWSRNGGLENSEDWSRNGAMRDSEDWSRNGAIKDSEDWSRNGAMRDSEDWSRNGAMKDSEDWSRNGMEDSEDWSRNGA |
| >ML21545a |
| MSKFLFLTLLVGAARAASFASDSDNTLADSDECRTGYNMRYDLECRRRRAARAAARLQLQEDGETEGGEELTKRSAEEQNWNYSARRDRGKRSN |
| NEEQDFSGYRRGSGQKRSAEEDDYDIYKRGESEEQDYSGYRGRGRGE |
| >ML016347a |
| MICLNFRVTAVVLFVILLAGFCPAAPIYSTQENDAFLETGEEKGELDIEAPKTPKLRYFYQKRSEEGKDEDPKLQYFYQIRSLEDKEKKPKLRY |
| FYQKREENKDTKPKLRHFYQKREENKDTKPKLRYFFQKREENKDTKPKLRYFYQEREDNKDTKPKLRYFFQKREENKDTKPKLRNFYRKRGEN |
| >ML07842a |
| MMKLTVVCLLFASLLVRAECGVVERHEEQLFDAEDEQPNFRAGLENKELEKRSPNFRFGSKRSAEDAPNLRGKRSNEDTPNFRGTKRSAEDSPN |
| FRGAKRSAVEAPNFRGAQKSEEDAPHFRGAKRFAEDAPHLRGVKRSTVDAPNFRGAERLEEDVPNLRGAERSEEGSPNFRGAERAVEDAPQFRG |
| >ML14597a |
| MKCFTLLLIGFLIAVETLPLNSHEEDDLAVDIETRKSKGKKHGLVELEERKSNKKHGLIELEERKAKPAPAPAPKKKKGLVELEERKSKKKHGL |
| VELEERKSKKKNGLVELEERKAKPAPVPQKKKGTR |
| >ML06743a |
| MFRVFVSIHLFLHLIAARSTGQVDLEKVDRKAGFFWKPQSKRAVAEDATAPPEDDVGFFWKPKFGKRSVEIPKPNDDKKRAGFFWKPQFKRSVA |
| EEATAAPEDHAGFFWKPNFGGRFGKRSVVKSEPQDGKKRAGFFWKPPIQESRCRRRHRCSRRPFRFFLEA |
| >ML06405a |
| MRKTLCIQLLVLLAIQRFTDGKAFEESENEVLSLEKRAEDLDDEDFFDKRETEDRENEVEIDLD |
| >ML02212a |
| MKLFLFVLLGLVALISCETVESEVDSELSESEDSNAMRVKRAKFSMSNYRGHKQGNRGWTGGAMQEEE |
| >ML01798a |
| MKLLLLTLAVLLACLTAVPVRQEEVPQEEIRLERSAESSGGETARVEKRAAIDTGSDYPGFEGGKRRWYG |
| >ML14991a |
| MSCLSNLSPCKSLLLFLGLVLLTKESVQRTLPESGQALGMKNTIEEDATVMSDSDEEQSRMKRQFFRLKRQSFRPSRGDAEWNLDGYFGNQEKS |
| EAELRDPDTQTRRFKDGRNEQLSQWRAMMNRV |
| >ML11723a |
| MKYFMILAVLILTVGSSPFEKRDAENAELASEKRSEEADPGEEYVLAKRNVDQDAVSLEKRQKYGYKRGE |
| >ML003517a |
| MATMMGFLIVLLIAAVSGHSHSDESPQHNRPDSPPKHHIPKDVIEALQRSGKSRGEHNAHPGPAEHHPEGAHSSHLGKGHNFERHPEGGRPIPP |
| KRVRQRISRL |
| >ML02736a |
| MKCFVVLFALLALSQSASLNSLESVEDVIMADNDNVELEEGALNAEEEARVYKGYNGGNRVWYG |
| >ML017711a |
| MKMFILIGLLITLVLNYVSAGDLARRSLEEENESGINEGAAEEDDESFRGLRESEDESRGKREVADEELFRGKRDFREKRDFRGKRDFRGKREF |
| RGKREFRGKRDFRGMKSSEEELVKRDLSRGQ |
| >ML030511a |
| MALTKILFSLSLILMVSSRAFEESAGDENNQAFGLGNKDGAVGDSLEEITQLGKVDEFKLAEAEDDQHFAVGT |
| >ML206415a |
| MKLHLYIGTILLCLSAMFLDGIAQNDDSADDNAAGQTDQAGAENSEDDAAAAAGEEGAETADGAETNSNDTTGAGAEGEAAEGDSKDPNGGAMS |
| NLKNSYLLLLTIPVVGRIFV |
| >ML00992a |
| MSIMKFWSLLLLTLAITTLSAQEPAQDDSEAVNDTSPDGDAASENVADDAAPADGEAAAADGEAAAADGEAAAADGEAAPADGEDGAAAEDGAA |
| AEDGAAAEDGAAAEDGAVAAEDGAAAEDGAAAEDGAAAEDGAAAEEDGAAAEEEQGEEEAEPAGSGSNVQNKICFSIFVVSILTKVLLC |
| >ML218923a |
| MDKLMLFIIVIEVVIISGRTIISSDDTNLKSSEDFGIKGKGPWNSEEDFGVRGSGGWMSEEDFVVKGNGARMSEEDFGVRGSRKGTNWEAEDLA |
| ESSEEEIYGIPGKGVWNKEDQDLLPGGAVPFSN |
| >ML056913a |
| MFRLTSILLVLVLAIVTFTRTIEDSEEIGHGLRMDKNEEIGHGLTLDKRSADSVDTEAMADEEELVGHGIKGSHAWRK |
| >ML030510a |
| MKFTKLIILMTLLALIASRSLEEDNNADQTIGWGRQRVAEAEDFVAEEKMAGGGNMILEMSEEEQHLGLGF |
| >ML066512a |
| MVFYSKSVLIGILALVCLTYARSLDSEFVDDNEILVLEKREEGDEEDSLDLMDTGSEIEDVVDEGGSEIMIEEKGADSDREQIQPRDGFFFVPD |
| YGRIHWPKSKKRPRKQPSKSVRINSERL |
| >ML233326a |
| MKTTFLVLTLMIICCNYVQSVPMSLEEDLSDEEHRGLQKRAGTKFNKADYKSVGEGTRKWFG |
| >ML199816a |
| MKLFLFLTASLLVLATVTVQTEAREIVAEVAESEAVESAASEEDTFVYRKEEDSAFLFAD |
| >ML31983a |

MTMKIFLVTLSLVALFAVAAADTEEQAKEEFELADEAEDSLEEGDELVEVVKRFADKKKSKKNRMRRW
>ML065755a
MKLAAGIFLVLACLTVVMIPGEAASLGSLDAANSDSNAVLYGADHEILESSLINRKEEAEGALYG
>ML10665a
MRRAALVLLIALTMAILLEFTENSLVEGHSSGSRRRSGSSGSRRRSSSNSGFAGSRRRRYRSNSG
>ML215411a
MKQRITILLTILGVIVLAQSKSILVESEDVLLGDNAELEVDSNSVDLDEELIGSDIKLVPGSGGNPWGRK
>ML124215a
MSRMLFAVTLVIFSVLAITSGASLSEEDEELLTGLEDQMENSEETMMYAASDDADNNEAEMAKRGGKARYRRW
>ML216920a
MVLLLVAVCFTATLSTGKCQTQGTVVTTSTPPFNCSADDLGQITEHNKYRLLHENTAPVCLLHSLVRSAQEYADTLAKLNTEDPQGTARKLPHS
>ML01134a
MAEIRSLFFAVSLIMISGLSFGMRVEQFNGSTNKCSRHQNFTHERNSVRGKDTGPVRHFNKRGTCTELDPEWSGLTTETECLDVNTKTLTLIAK
EVVNRHEQSSTNETEKNGEDL

***-Not enriched in uncharacterised metacells***

>ML21632a
MKLFLLLALFGMVMFAHCEEGAEAAGEEAAGEEGAGEEAAGEKRVLVKRLLVKRMLLVKRLLVKRMLVMMLLPSNWLFSPWPQLLPDSSKSRTV
LRVSDS
>ML093052a
MTKFGGTVLLLGLTVVLTVPATGETDIARKEKTKAWVTIDMTPTRSRGQELTPTPTVEEKPGQVESPDKLDDPEEKEPPRREESGENPQKSKSD
TEYDVESEDKNSAAKLECATILILLTIWLRQ
>ML030512a
MMKFTKTIICLTLVVLVTCRALEEDEQTIGWGKAAVAEEDLDTVGLNMVKEEQSLGLGLEEDSMDRPGMMV
>ML090813a
MKVFLLFLAVAVFYCATADEIEEEDMEFDTEFELEPSAESEMETEEHPGHKRGPGGRRHGM
>ML31164a
MRLLGSTILLVSFLLVVSSVDQLQIGVKKRNQCERKSKKGDQLAMHYTGTLKSDGSKFDSSRDRNSPFEFTLGVGQVIKGWDQGLLNMCPGDQR
KLTIPPHLGYGDRGAGGKIPGGAWLVFEVELLEIKNAKREL
>ML319815a
MSRLWLLTVTIVLCLVVVNSYIVEEYEFGLGSFSKAAENAANRAKRAGQFKSDHKKHHIRHKPHRGPRHGPRHGPHHAKKSMKKHSKKTNKGIK
SKKSNKGKKAKKGKKSKKGKKGKKSKKIKKGKKTKKGKKLKKGKKSKKGKKSKKGKKGKKPKRGKKSKKGKKSIKGKKSKKGKKGKKSKKGKKS
KKGLKSKRGKKAKKGKKSKKGKKSKKGMKSKKGKKSKKGKKPKKGKKSKKGKKAKKGKKSKKSRKSKSGKKSKKGKKAKKGKKSKKGKKAKKGK
KSKNAKKSKKGKKPKKGKKSKKSKKAKKGKKTKNAKKSKKGMKSKKAKKSKKGKKPKKGKKSKKGKKSKKGKKPKKSKKGRKGKKSKKDNKPKK
NKNGKKPKKGETSKKGKKPKKGKKPKKLNKGMKDRKSKKGKKGKESRKGKKPKTNKNGKKSKKGKKSKKGKKPEKGKKPKKSN
>ML07012a
MKLFLYLALIGMIAFVNAQDAEPAGDDAAPADDDAAPADDGAAEGEEDGSEDSSEDGEEGADDGSGAESIQFAIVAALPVVARLF
>ML034332a
MNKVYLVLAVTAVLYCATASEEEASVTDVTSEHPHHHHHHHGPPHHRPHPHPPHHGPHPHHRRGPHPPPHHGPHPPPRHGPHPPPRHGRRGHKH
GGKRSRRVKGRSGKKGRHSKKVRRGKKGGRKGRHGKKAGRKGRRGKKAGRKGRRGKKAGRKGRHGKKGGRKGRHGKKGGRKGRRGKKGGRKGHS
GKKGSRKGRRGMRSGKKGRSGRKGGKKGRSGKRSGGHGGHKQHRGHSRHGGHHHHPRPPPRHHHHHHHHRRPPPRHHHHHHHHHRRPPPRHHHHH
RRGE
>ML07361a
MKVFLLFLAVAVFYCATADEIEEEDFDMEFEVEEGPGHGHGGDGGHGGNWGQGGHGGHGGHGGHGHHH

>ML051421a
MLQTLVVGLSFATSEAIAETFGSVMERYHNTRFFNPGMANDDVRLQKEMFLKLNGPQLGAALPLCQRVADRMNVRLTAAYSALPPIHREKKVGK
VIRRLQSEKPGYFK
>ML174759a
MTGKFLILLVVLLNLAPEPCQGQIGVADWWAKVLKNVASSESQEDGLVDDGEFAKFVRREAAEMEQK
>ML33461a
MLSHLLLLHVLTAGVTLTGALTDLNQVEGLRNLLKGLDGGVIATCNIVFFSELVNYTQADNNCKKFDIGTGRGEDGNL
>ML065715a
MKLHFCIAVLIISAALVLETRAGSRRRPANVRRRPVKQPKYSTSYSPNTSSARYNRRTSDPPSSFSSEARTPPFPWRTSPSSGAETTARSTPPL
PTSQDNTPSTTGSRTSSKSSAITSGSSRVSESDRVTKSSSQSTTPQDYTGPTTAEKWYAFNSDTGTTGTYTSPAPGLSTEEYSGESSTIFPEEP
TTPDIPTTPEEPTTQPSSKGPLTTETPRYSDEASSEQSTETDRPPSESTDTTKLVSASELTDETPGSTEATVSKTSERPTTTSSRTTSESQTPE
LPCGRGKKRGPKGRCDNRGGKNRDDDRKKGKGEARKKDNDRKKGNGKF
>ML067017a
MARSNSAASLLILVLVLLVDRATTLNTTESEQPPGPDSPSTSSTASPIPQELPKPNPTTATPALPDPELENEIEGGTESESSKSESDGNAADSS
NDADSVGGEESATDDTKTENNATNTTSATDSDSKSFAFRPLLLTAIGIAIAVNIVDAL
>ML040029a
MGFKVLGTIFVVLCTEVLAKSKDESDTEPKEPNNTSENWMIYGPIMVAILLIISCVFSCKLRNWYRRNQIKKMSPAQSTEGKKKAESVTKKEDQ
VDGTEYRLLTPDYRSRTPSTHSNNLPLQTSAYRPMTALGGDARYILIRDNGTDSQTTTLTRDERV
>ML46396a
MKVFLLFLAVAVFYCATADEIEEEDMEFDTEFELEPLTDMESEVGSEIETEAGERRSGM
>ML11691a
MKLFVILLAVTLFLTVYGSHSSSSTPYREREPPTEYKGGESDGGESDGGEGWPDEDYMSGSSRCGFTFSLTGVVFVAAIFL
>ML05367a

```
MTIPLFALTLALLALSNSASISDEEVDRMEGIEESAEGGLLIGYPSSRLPSIRSRVSQDSGLSESEQTW
>ML26664a
MRLFILLAAVFAVVCVHAIEEENDSEDVSGEFEEMTMEYEIPAEGKRRKKFANLGGR
>ML039810a
MAIKWLIISLFVLTVFSAPVEIASDEEILSEPPEDGPGGGDRPGGDRGGEDGPPRRDGPPRHDGPPRRDGPPRRDGPPRHDGPPRRDGPPHHDG
PPRRDGPPHHGPPHHGKPHHGPPHHGPPHHGPPHHGPPHHGKPHHGPPHHGPPTMDHHTMDHHTMDHHTMDHLTMDHHLMDHLLMAHRLMAHIH
IPLRMDHVHIPLHMDHILILIIILILIPILILILILILILIPILILILILILNVLNVPIVHAVLERFTVVLEFRVHKRTFD
>ML218823a
MSKVFPALLFAVMAACVFELGSTAAVELEVDYQNNLPEMEVAGLDVAESELSLESEIVAEHNAEEGGYFGQ
>ML083811a
MLIAFGLLITTTTLPSFETVGLGEFDSGFDDFGSHSGLGSGYDDFGSDSGFDDFDSDSGFDDSGFDDFGDYSGFDSGFHDSGSGSDFDDSGFGS
DFDDCDFDSDSGFDDSGFDDFGDYSGFDSGFGSDFDSSASGSDFDDSGFDAGFDDSGFDDDFGGDSGGDSGFDDLGFDDLGFDSGNDDYYDWGD
PGESKQELLFL
>ML005019a
MKFLILLVATVALSRANDDTARLQLAADNPQKLQALFTEFRKENNRQYLNPVEARMRMSIFRFI
>ML05859a
MKVFVAVALLIVVAAYEEDDVIEEENEVYSILGESEPEDYDAYMAAIEQEMPETEEKELSADFEDSPAESSAA
>ML090812a
MKVFLLFLAVAVFYCATADEIEEEDMEFDTEFEMETEEETAPNRGNNNNNGNNRNGRGM
>ML032113a
MSYTAVLLLLITALAFILSVEGAPAAHIGGNQRRHNLALMRRNNQRAENQGLTESQMDSILFENLAENSLSAEKELESKQEEETSEEQASANRS
QNSGYRRGGYRVRGTGRATHSRRHNKQAK
>ML100012a
MKLFVLLALFGLIAFINCQETEETVGGETEGGETEGGETEGGETEGGETEGGDGSGASSVQLALLALAPVAARFF
>ML06328a
MTHKTKLCLLLLLLQFTTYVAGELFGRRRNPGSRRRAGTRNSYTPNYSVSMTIWYVVIGVAVAFVLALIASIVYYFLVVRKRNQKRKEAEMYKP
VGGGYRSKADRDKEMEEYLAHVRVAKGSTPNYDQFVARDQVHT
>ML1541126a
MNSHNVLVVFTICLHATYGLSIVKRSADFAEEEMQTPLRGLSEEMQMPSRGGGLSEEMQMPSRGGGLSEEMQMPSRGGGLSEEMQMPSRGGGRS
EEMMQTPLRGRNAEENFVPLRRMSEELNNDDLALQEMNEAAQDEVPLDMDSPIVPYNRYKIRHLTKTRERDKPKNTRQKYRHRKKRDILADRKV
DLTAIFRRARPWWTPKNV
>ML073260a
MAYLLLFSTLALNLGISLAGKNLFLVPLTSDPDTVTNEQKLDLLERRLLQADHLEYGVAAGIQMAPRSLNLVARKERSVSGDSAPAARQRLNPK
FQSRKRRMLDGTEPPLGWIMY
>ML08268a
MKFFLAFIVALATFSIVNSRSIHSENPRTEKDDTVAAETSPKTIDGSLSSFIEQKLAEAGLTEEVSEEGRQKRSSRRHPARHQKKTRGHNKPKH
NMSAFFYVPPNGK
>ML258215a
MLKLVLVLSVLTFSWCAEEAAAGAAAGATEGAAAGASVEAEAVAKAAEEASQETAEEALDALPDEAMGEGDEDLYDFELEDFDNQDFPEEEGDF
PEEDAEMDELADESSEEETILAEEEPVAEATPTEGATPTEGEKPATENPELK
>ML053618a
MRYVAAYLLAALGGNASPSADDIKNILSAVGVSVDDEKLSFVMGQLKGKDINEVVASGIGKLASVPSGGAVAASSGGAAAAAGGAAAPAEEEKA
PEPESEEESDDDMGFSLFD
>ML053619a
MRYVAAYLLAALGGNASPSADDIKNILSAVGVSVDDEKLSFVMGQLKGKDINEVVASGIGKLASVPSGGAVAASSGGAAAAGGAAAAPAEEEKA
PEPESEEESDDDMGFSLFD
>ML00497a
MKLLRFTLFVLFGILSTTFSLAVAETDEGGAPEDPDASELDTGAAEGGESDEVPVEGGDVSSDTADKEGEDEIPAESNDDAQVESKESEVEEDG
ETDPNGGASAKDLAILVIVPVVARFL
>ML040716a
MGTLIIFAVLSGLAGLATLRLPDTKGVPTPSSAEEVQSRRDRRITQTNTVFSNEAAA
>ML154515a
MFCGVSLLLLFCLIILEIRGAPIPKSSDYLSSSPPWISHSGSLRPRHIVIDLEVILGRTEEPLSSGSYPTAEINYRHIPNVGIEIFPSTAPGST
EGNKFNPYASMKTTNDVLRSRKDFLESNTFAKSKRKVGAFSDDDEESEEDLQQDEKR
```

***Cysteine-rich precursors (>3 Cys residues downstream the signal peptide)***

```
>ML358831a
MQLRAAALTLIVGLALADQCPESYPNVYYNGQYCCSSGREKFYAPQREKCDGSEIQRDSLCCQGGSIKCPTDDCDDYSAPSPDQCPESYPNVYY
NGQYCCSSGREKFYAPQREKCDGSEIQRDSLCCQGGSIKCPTDNCDDYSAPDQCPESHPNVYYNGQYCCSSGREKFYAPQREKCDGSEIQRDSL
CCQGGSIKCPTDNCGDFSG
>ML09126a
MSFRILKAWPSTYFLLVFSTVNIAKYDDIYNYHSGFTCPVCYQGFSQKGNLNRHIRTIHQQIKKFQCDRCKRCFTQKTCLKVHQESAMKRGSCY
YRISSD
>ML375910a
MMSTKVIFATLCLVALVKVSVSGPASGVGTCFVGDKTYRVGDKWAQACNAVCRCRAPGTFECVPMCAPAPPPRACLKTYTSIEDPENSCCTIKT
CPSEQV
>ML093044a
MRLTTFTTIFLGFIALMFADMKCGGTGDIKPADDDIKALCAQVKGAVEDHAKKSYAVFEPHSYKTQVVAGTNYFVKTSPEGEFLHLRIFEHLPC
YGGGVYLHGVLHHHEVGDNEVDYFDQNMNFDEPKWD
```

>ML22307a
MMLNAVAAIALLVTVTSCCSIEEEITRISVREAMRSELEAKKEVDVARQEMQEMKKELAWYSLHVAAAAACRGSTPSGGTGPHANVVLAKENTR
SCDDQCADTYFTECDADVSIQGDFGKATSYTSRPGLVTSTTTAAPQPATPMSNLMKLKLMTTGS
>ML223012a
MLSSLQLALVAALVASTHSIFCSDTHPYCSGGKVCLHNECRRYKPNIDCTTSGGKCPRGYCCMKVASYSAGTCVRKQNFYLQLGDLCQRSGGCD
YCECQPGLVCAKNGFQFFPEVYTCQQIDPEGECDEDDQCASDRCCSSSKCKPKKKQGEACFPKNGYFPEAYYECNGECEDGTSCSRDHTNPNFG
SYVCTVVEPDPVNLCRGS
>ML056970a
MATTSTALLKIVIGVILFIALSDAVALGNRPPWNQWSYAGRALKKRSETETQIEKESGELLLETRGKKGRSKKKGASLIIETESGKTIEAEKSM
KIDEPDAISLNVECGLWEVCSVVDNEDSCIDIDSVCPCRKEASAMQPTPISRVKLLKKRECD
>ML08822a
MKLYFLTASILFLISRISATTYSVSIKETVLKNEGEPHQIREYQVDFGGPKDKRLVECIIETLGSSNEVQTCECDSRKWKNKPTATFLSSLNFR
FQQNRVYLDGVYNCVYQWEKRSRSDFKF
>ML22382a
MKNFLFTVLLAGVNLVLGLADGEDFLCVQGTNSSAAPTPCLDSARSCYGPKFTEHIGLASRVDFGCGRCEPGLHFECEECQESGCNQPQDLGET
FSCYNYKVAGGAVTRHPTLQTCFRYSHAGIICNMPSSSTDYRSTSGCGPCGEQAKQSGSCIECYSDSCNIIQTPEDRFRSQRRAKRSAVDCSAI
CPYR
>ML046720a
MGLILLGYILLTIKLVAAVDGIKDEGDIELIQLLVGREITIRLNLDKIYCVRTFQVFYSEVSSVELSCTSSDCTCGGPSKEFCNSLLFAVDIRR
NELSDDLEKRSDCAYGDTVTLSWLADISDVTVTEIALTEFQARETNNEENNEENNEENNEENNEENNEENNEENNEENNEENNEENNEENNEEN
NEENNEENNEENNEENNEEEDCKQNLESCHTSHGNMRHGSLLGCSRPYLVDVVD
>ML017938a
MKLFLVVFLTCLYLAQSLSVLPQVEYEEFVVEFSQGENLDELLEGCALESESGPQFWNIWEIIRRLHQLKLEIVCGATTTELLDLLLIPRCLRR
LVNIARKIVCRDYKPPTTPPTTTPQPTTTPQPTTTPQPTTTPQPTTTPQPTTTPQPTTTPQPTTTPEPTTTPEPTTTPEPTTTPEPTTTPEPTT
TPEPTTTPQPTTTPQPTTTPEPTTTPEPSTPEPSTPEPSTPEPSTPEPSTPEPTTPEPTTPEPTTPEPNTSAEPTSAEPTSAAPTSAAP
>ML306119a
MIKLLLVTVLIGAAAAFKCFKTAADGAVSSADCEAGTASCHSPTLNILSGMTGQEYGCGDCADEAAAGTCVECATAECNKEMKAVEFECSPYTY
TDSKWTEGETKTKCFKLEGTDGKCN
>ML23459a
MEKWKSHRGYLVVLLVLLTPAPSLAAVSVTVTNDAVLSGETGTITCTVTASSSETLTLYQWKNPGGNVIGAAPGTFTADGVSYTIADLGLSGNQ
IVNSKLEASSVSSDLTPLTCFALTSGGAAQGTGIVDVVSVTPTRNAIVTGGSSTVSCTLAGVSDGANPTAEWSQGGSTISPDGTHTLSTDYTGS
NKIAKLTLSSVSSDQTYTCKFTASAGGSVSSTVDLDHVTITTNGDVVKSGGTADISCVLSGASQAPLSRKWLDGTPADITANNGNPYLYYDNPP
FSGGGQTFLLQINPITTSTTFTCCYEFATGSCINTPAQVDVITITASNYLVVVGDNLDISCVLGNSPSQPNSITWKTSAGSISNGVNLGTFSSN
SQTTTLSISSIAEDTVYTCEFVIPIGGSTATASKSISVDTVTLTPTSGTAQSGGQVTVSCQIVGAGVQPTVVWKDGGTTVGSGINNDPWAPSTG
TLLSKLTVSGLTSSKTYTCEFTVNGEVISKTATANILSITAIGATIATGSAATVTCTIADSPAAPSSVFWKDSSQNTVVMSNGFSKSESAFNAG
GQQATMVLGTNSITADTIYTCVFVVEGASYETTTSVDHITVTAGDTSKLSAGSATVSCVLSGAESTPTVTWLDSSNVPKTSSDSGYTVTPGTFS
SGGLTTTLVSTSDTTDTTYTCKFSFATGVVEKVTTVDVVTITGTNDAVKSGGTATITCSLAGSQFAPSDNAWYLSDVKKSTGGDYVANLGSYDS
GADTQVMTLQKSSVSATETYRCSFSINGDRVDGSAKITSLTITAPDYAVKANEGIAISCVIANAESDPSIVWKTNTGTIGSGIVDGNFVGGSKT
STLTLGSVSSDTTYTCEFTMGADTVSSTIDVDTVTITTTDGTALSGDAVTVTCQVSGAGTEPTVTWEDGGSPVSSGISNDPWVANTGTLTSKLS
VSGLTSNKVYTCKFAVDGSPVSSTVTANMLAITVVGTTVYTGSTSTITCTISNAQQDPSLISWKNGDGNNLVSGSNSYLITNGALSGGSKETKL
TLNTNEISEDKTYTCVFVVASASLQKVVAVDHVSITPSSFAKLSAASATVSCVLSGAGSTPTVTWLDSSNVPKTSSDSGYTVTPGTFSSGGLTT
SLVSTSDTTDTTYTCKFSFTDAELTKVLTVDVVTITPGNDIVKSGGTATISCTITDTQFPTTLAAWTLGQTPISTGGDYNVNVGAYNNVAGNHV
MTLERSNAVADETYTCTFTTSGVTVQNTVTTTVFGVTSASDTVKSGETATLTCGISGLDVEPTVAWLNSAGGVVTTGGDNTVTPGAWSSGSIST
TLTVAGLSADASYTCQFTLSDGSKTSETVKVFFVSVAVTTTTETVVSGSSVTLTCTLSGSSTAPANPVWNDGTNNLSDGGGIAINLGSFAAGTQ
TSTLTVSSVTKDTSYSCSFTISATTVTESKAVDVVVVSVPAVTIAQGEVATLTCTVSDIGSAVTISWTGYSSGITPGSFSSNTQNSVLEVSYIA
SDTTITCSVTRGSFSTTSDGAVTVLDPCAKGSYFQTVSGTTTCTKCALNSYSDNPNLRSCTSCPNSGSTLFTGSSKITQCYTKTSDHSVVMKTG
SSVTLTAAITVSDTVGSATWTSSLGGKLSDAESSSENTLTSTLVVSSWAAQTFTCTFSVTRTGESSPASTAVTITGLVAAIIDQPAATAQSPGA
SYTLSCTAPLPPSGLDYKAGWLKDGEQVDIIMTTAAQTLSDKIKSTLTIGYLTSDMTGNYACFFTYALTANLPTSTYRVTSDTTHLAVEGISLF
PAQSGGLAGSSATITCLGVSSADADSVEWKINGNPVIYSTTNTVTNTYTTVSTSKESKSVLVLGSISSADEGSIDCLMLFDVGWYSSTGVMNVI
SIVASPVDVNGFTGTPAEMHCLVEGPPSPTIEWYKGTTKVDDSLSTISTIGSNQYNSILTISSVSESDEGLYKCKATWATDLGVTGGTTESKAA
DLIVYGISFLTANQEVAQSSNVKLQCRIDDLDTTAIIWKKDGVTVSGDGIQRSTLDSWSDTISADTSKIGTVQYSCQAKYTHTSGDISYSGAQT
STLTIYKTCTALSNPTGGTVVCSGSDSDKTCSVTCNSGYVPRANPRTFTCTDGTWDLQPSNTDCTAKATPSSYRLTYTAVYNYPIPCGTSWGPY
ILAAYPYHMKNAEGSLTNVPAVVQMDATSFVCVNDPTDCTSSSTCQQIKIKFQYVQTTSLSNDDPLQTTRTEIGTYVSSTSLNVLGVLNTAGRL
DSHGNLQGSYGNKKKRSTEEKYSGPGFVMRAAKGEVESSVKCPEFSILVDDHCTMCGQGWFIKETDCAPCPLGTYQAEKDSNKCTPCPEGLSTL
VTAATSPLMCVEVCQVRQPLHGTVHPPEGSLLTSPSDVTVLCDEGFTLETGRRHGEVSCEEEVHCSRIQLIGVPEFIIRDTDLTVDCVIEAEFE
FENCTLIQDGAAVQKSSPFFDNNRQICSFDVFLTSSTELSCSAEGPDLYLKGDPKEITLLRPTAKANMDTFMVSDPINLACSAIVPRGHSVFMR
WKKHGEVIGSPSFSQTTLGLLIHSKKQAQFEDSGKYSCEVEYIGIGVSESESIPVRVVGFLEELSDFSLYKGHSVRISCVLPTGSSISMLEWYR
DGERQNDQGSRIDGEEIRVITSELTVAYPGTYECKGDTSGVPFSSVAVVDLNENYGFVVHPESQYIVEGGYVVFSCRFSQDDVLVEWLLDGSPM
GEGQNNRRTLRLASLTTDKLVQCVAKPPTGALIFSKSAKVIVRRFITEPSDAVLGDEPALFHCSVYGPLVQDIVWETDYGVLQPTSTTQLTAVE
LHATLEVSSSNKVRCAVTFQDGDRLVSRWVNTLTYEVTVSEGFVLLNQGFIARCRIHSDEAPTAVFWKMRADVLYEDSDVIYDGYQGYSESTYY
LTSLTDAQVAELTCTATFRGTTITVTSPQRSLAPLGIISDPRSVSMVRGSGDEVEAWCEFYDYPEVDTVVSWSLDGLEELEVTTIYVRSNIARS
SVRTKVTTEGLFDVKCSVAYSDYGIVDSYTALLSVLEPVTLTVASREVGYGEIAEVEGRIPLLGGADPGYHWSVNGVYINEQEHLHETNHLKSI
LRWPIMQDSTITLTVFSDGEITRESATIQVYGIQSVSVEGAVEIGSTTDLACTVDVRTPPEEVFWYFNTIKHPSTSSTSSDTSSKSVLTLPDIT
PSSFGHYTCVARYADRPSVFQRLALVPVGACTLPHILNGGFPTEFISEGSSATVTCISSYVPVDRDDTLTCRGGVLEGRTPVCVRVLETVDDSS
ASLMVVLATCACVLMTCSIVVSFIFWYKKHNKVTTKLSVEEEGDPYPAKSKPPEIPELEIPHVRGDEREDETLQYDPERKVWLTKHH
>ML460824a
MKAAGRVLLLTFLQVIGLVQSTVDGGYSAWSAWTTCSAACGGGEETRSRTCTNPAPANGGVDCVGERTETRECNTQGCPEENNDSSGDDHDCEK
NETCNSDANSVVAAASMVTVSLLTQLF

>ML008117a
MLRAAILALCLSAAWTLQCYLCDTTDDDVTACSTTVCNGLNAVCYKREYSFDADKDKQYEERGCVENKSYLDGTCKMTLDTVMEDVNSCKIYTC
DGNRCNSSLQISASTTLLAALFAYIFM
>ML23837a
MKSTASILLLLGTLSLATAFTELWCETCVYAYIEGSGRVYNCRSESPNGRICPPSHKCYSLKYTYQKTGTNFDLGLRDPPLGVYTWILQGCIFE
RRDPQHLCDQKKKQFRNLRPTENHTCEISVCEGDKCNQELTDDYVADLD
>ML107912a
MFKYFFFSSLVLSVLSDRPARDCSDGNQADYRGDTRQTWSGIPCQKWTSQRPHNHDYDPAEWPEAGIGFHNNCRNPSGLGRRAWCYTTDPDLRW
DYCDVPRCSTPEVKDPECSSGNQADYRGTMSITNSGFTCQNWLKKEPHEPDYDYIEDSSTGIGDHNYCRNPHNDDTGTWCFTVDDPDEDWDYCA
VPKC
>ML115614a
MKILSVLAVLAVFYRHSEGLKCQTIADFSSGDVGALALTEEDCTEEGVTTCV
>ML10906a
MMGREYFTTMKLFFLLFLFVLVTVSVGNSIEDGSNDPSGSVKWWREDGRCGVNYKLPNGRPAECNPLHPMGHTCCSDKGWCGKTSKHCQCPGCV
DYTARWRKDLRCGEEYALPNGYPAQCDPFHPQGKSCCSDLGWCGKLAKHCSCPDCQDFRALWRKDLRCGPMFKLPNGMPAQCDPRSAKKYSCCS
KKGWCGASVQHCKCRGCVKFTDSGPIEFNQSTKKWNKVKSRGSRKVA
>ML00857a
MERNAILVCTLLLLSISLTIAEDTTTPFTKVTEKPVTWKSEKRWTEKITAAPVRCPAGYYLEGTSTCKQCGRNTYSEEGSSSCTSCPEGLVAPA
GSQSSDACEEATCPGNKEACPEGSGCFDIWQRCDGKPNCPDGSDEAVELCEGEYCPEGTERCPDDSRCCGAKSFCDGTRNCDDGSDEDPLFCGE
YTCPGGMSKCGGGLKCIDNWRFCDGSQDCKDNSDEDPDFCAGYECLGKRTKCGDGLQCVLMQRVCEGKSVCKDNSLTPDEEFCQGDEFSVNNQG
SINSGSSSSSGTSGGWSKFCKAGQHKVGKKCEVCPENTYSLAKATSCTPCPDDKISPAGSKSEKACDYDPCSAGDYMTESGCQQCGENTYSGAG
ASSCTSCPDGKMSAAGSTSEADCQHEQFPTCNCWTPACGYCSNMECTVQQNLPNNQQGFEVEVTTSVTWRKTNTIFLYDANGHVISRLKWNLET
IDLTGCGGCKTPTELRELSKGADTVSWTFSLKDGLFQISSGGEVFYERQLPKECASIYENIDRFSFSDSTCEGSYSFRSNEMERGAKMDSDCGG
VCPQA
>ML020020a
MGRWVTVPILLLLLLDWVEANCKESGFTSGLQCSSCDKLGDHNLGDLEGEAFVNDEKKDKQYANLEIEYARGASPTLILKNEEGVEVESLAIDK
WDTDTVREYLTQHLKP
>ML034338a
MMMRNLCIVGVVLLQLHLARADAEAEADPAACCRKRARFLKGMVGDTGLPGPKGEPGHIGERGDVGPQGPPGAAGPKGTPGISGLPGLKGEQGE
PGETGPIGERGPPGIDGVDGEDGINGIDGCNGTDGINGTKGDTGPTGPTGDQGEAGEDGEKGAKGETGDVGEDTQAPKGEPGPKGATGQTGIAG
LTGAKGDPGSNGLDGEDGAQGPPGEPGDKGVVGAEGPQGSEAGDQGLQGLKGPAGSDGKDGKDGAVGDTGEPGPKGEP
>ML12596a
MLRGLVVVLVVGLAVADNRLSYQTHEACIHNGTKYEHGEKFQDGCKDCWCDKNEVRCIQGSRCGNCVYVKPNGYTATAEPREVFWDGCRWCTCG
NRGTHDCHMNGCPHKCPFKNRDGVDGFAQFFQIWYEGCHRCICRKGWAWQIGMGWAEKDCTFDCDQNWL
>ML18206a
MQILPLLFLSTFLVLGHCGCRGGYYETEEGDCEMCGKGNYCPANSTEPTPCPVGTYGPASTASIERYCYPCMGSYNNQTGAVRCTRCGLGNFCP
HGAVNPTPCPIGSFGPSTSSKRCYPCGSGSYSNVTGASRCTDCPAGHYCESNSTEPIPCPTGSYSRHGRAGRCYTCSAGRYSSVTGATECTACP
AGYYCENNSTEPTPCPEGMTSSPSSRSVRYCRDIRPVEGDCPAGTFKNDIEECEQCSTGHYCPEGSTERTPCPKGTYSSYSNAWKEEFCRECYA
GYYGDEEGLASCKSCPAGSYCEAGSVEPTSCPDGHTSYPYSRSEDHCRPVS
>ML13096a
MSFIWKVIAAVILLFVQIHVQAVESQPDPGCPKSHKRAFNKAENCCKAKSETSILLLPSHAKCLTRRTMKKAFEEGYECCVRNIPCPSVCQEKS
HVWGA
>ML23996a
MRFTLCILLLVLLAVTLAFPDPDGYRKKGKKNKKNKDNTNQEDAEDDETDEEEDEDITKLRCVNGHVSPTKTVANFCETDLSFNCMKVYAADNP
FTLLDTSHFIYTCVKPGLNCKTMYDTYLKNAPEGSYCEECDRKYNGCNAEGQNELYEQAFGGKQVE
>ML214326a
MKLPVLITLLVVAAVQAQDCTEGFTKCADGVQCIKEEYWCDGGDPECDDRSDEVEAVCREYQCTADYAKCGDGTHCTAVEYFCDGAQDCADGSD
EADCKGGYVKDGKCEDADVIMCNDGLQCVLDEFWCSGQKAHCDDKSDEADSTCLRYTCLEGYVKCKDGKQCIKESYLCDEGIPDCHDGSDEENC
>ML03704a
MLMFLLLTACLVVQASGGINCWECRKFGPKRDPFVANVIKNEQLETVESCEETMKECAHNKCYTINYYNATNSASTQVYGCLGRSWFGYYSERV
IKRYLQRKGGSDFSGWSTCGSDGCNAN
>ML26495a
MVFCILTVLSTAFILYAEGTSTGSGISWQEARESTQVKSYLDPSKYRECLESTHIRWFGLKYDDSLERRADSIINTGDNSDCDGGAVTDDDSSS
DDDDMSMCVPLPSSKYCDNYFERSRTDKIKQLYKGEVGCSVKYYRGGYLAACVYEDD
>ML00442a
MKTTAFLLILCVGLSAVSAGKKDDTDKKDPCKKASRAIIKCLKKGYEPTILSKEDLGKCSIKPQELKKKEIKRCAKREKEFSDNGCSQPCKKGS
DDDEETAAPAPPAPESTAAPPAPCIKKTIAQCRHSQQVYRGGIMSPVYHNDIPTFGACVEKCRNIAGCAAVVYIPKVSRCFTKNASHNPLAPEF
IRGRISFSLAMPCLEECSNK
>ML062222a
MLKHVCLILLVVALVTCKENGRGGRAKNKAKNKAKEMGREAAREQVEGRVENMDREAAREKAEGKKNKYKEMFGDGKSLSKKKLLKMTTDHLPT
GIHLPTGMKGVPEACIDAEFGESPAMEKIRRGGKDKFYVCAFVNFQHGVGEDKVSPPYTEETVDMRFALFKQCVDEVNKQNKRFRAGEDTHQAG
INFFCDLTDEERRPYSNGLVMPDE
>ML137713a
MISILLLILPATVATSGTEVDLTSPLPEEMDLRTGRTCNDSIDHYFPDPDRYLQGVAEEESKRVVEDRRFSCHSKYDYQYDPRVKVVCKRLFNS
VDSASTNVKLCELFLTDETPRTRYNVHSKITVQNFGKYGCAFYHHEGISAIACAFLR
>ML22161a
MRNFLTTSLFVVFCLLISVDSLDSNAAVDLMSLANAICRKQKVLYYRDGFEMFCKLSKELICPQTEKTVDAAAATCKPTLNVRIQYQMKKVQKA
VCQYEGSDNTTKEVQARLQDAGRCMYY

>ML279836a
MILLRVFILAILLVCAADAQKGPKGAKGERGSDGADGIPGSKGVAGFCEPKSFCPSNPGEPGEPGQPGLSGAEGEVGVDGYKGDSGPVGPKGPQ
GEPGARGRPGNSGPVGLKGETGVPGDDAEPCTTEECTTTRVNLDMCQDVYSLRGDRQRGYWQCPENHVITGFKGEKVMDMMVTCCHV
>ML03184a
MIITVELNPSLSLSTKMNRRPPCSFSLVSTMTRFSSTLLLLLLISPLLARPTLEKDPLVPVPWPGKIRSCDYYCMALWDPVCTQLGETYSNRCF
MSLASCWDGLERHVVHKGRCGENEDM
>ML05793a
MFVFLSLFALLLSATRPQLIPPETQEEIQECLRKRHINYHADWASNASKSGGSLVSFRRRRRALVERDDGPSTTRQTSMVLRGSCDEYFDVLGS
TLTELSTYTTNLTVGCSVNKQSVISCVYSYRLPSLI
>ML017312a
MTSTICRCILLMLFLGVLSAIAKKPKALRMKMREKRLKRRYRQLEKVDGGYSDWTPFSHCNLETCTETWTRYCNNPRPRNGGKTCVGLSFKKER
CRDDSKCDMILYPDMIFTLLASYLQTERGSLVPQCLGGCIVDTRRSGVY
>ML032413a
MFTILLIAVFFVAGLSEDVSEGISQDIADTIRKNSELLREGRTCMKDCIASAGCDDVDKECKKQCKTECYPGGHKKNGSGKKDRSQAKSKKERK
EKTQNGDRKKKGKGKKTE
>ML00879a
MTSLFLSLFSALLLQVKPDCSNVPDVEDRTGLCKSCRELPEHLLDCEDVLDLSEYNKTYQEDFVEKY
>ML200221a
MRLFLFLVVLLASVALVTSKKTKSSKSCKKLEKKFTKTCLKKGFEPNKLENCIIGEGVLKKKAAKKCMKLENNVLAADCDLKLCAGPEPEPEPE
PEPEPEPEPEPEPEPEPEPVGPNWCRHENSFLYSYAGPSFNSLDEAQEACLKNSRCNGITQEPYNSNRYTQRVGPEIHDWSPTGETSWTVC
>ML23661a
MFIRLILFSWLLFHTQQSDGVCPREEVPCAGTCAISLPQELNHGQAAIQCKIQNKRTYMGSRIDAELENACANAVFGGNDNTHIWSRYKLR
>ML154166a
MTLRLVLLVSVVGLTVGDVYLHGMRGSNNRLNERGRERANANRLFDSQNNNRGGYNQGSLAYFAGSIINLEWTNQHSCGDPNNHCEIILQYMCG
NNTRDGTTTRTIPDKRSQCEKWDCDTDLQ
>ML124231a
MRKLILLACLLQLSTTCSCYRPMGSDPYEEEPRASVLEVKINRVTKRPTASSNPYQISSPWANHGSHYRAFIEVSAVVTRVHRDAGRNFSVGDT
IRLSSRMSDAQCGVGGFLRVGETLVFRTYKKRSKELQLCEYLEFNLERLMSSG
>ML05977a
MELKFFLLVFCFLLAAESDDADRFGKARMGNEVATGHCQDRWHSQMDADCLSYMWDFYHLGNRFWKPCRDFTESFDRGWRKPKNRDCHRISGKC
FGSNNEQCCKSIYCFSRSIGRIPT
>ML329517a
MKLAIFTLLAAVLLHAALSNDMCSRKNLKCGTTLKGNYMRCLIHGYSSGYKCQATTVLNLPYDPTCCHTIEKRLFNCVKKEKINFDCMQYEDEE
DEILH
>ML25762a
MNTANVFAFIFVIFGLFESALSHECQPGYVAHNIEDSEDYECVRVEQPPPVPVALAPPRKQSSVAIEHAESEGCTTVDNPTVSNRVFSLDCPEG
KAVRTVEMRLGASPEIECCDIGVSRS
>ML33986a
MSLRIYLVLSGYLLGMATPIFKGKDHEGKESNEIEDWEYTRQQGCKTYTSNIKHMSCECLEHFSAHNNCESYVNFKDLKSLPRCKKLEFCLPQD
MGCHCRRNFCAIRGRCYVLWRSILSMKRVDHLLQWLANCNTFGVNDTSKRAVP
>ML087211a
MPKLAFLSVFLLLCHYAQSESRTSLQFQECVKKCTSPDTRDRIKCKIGCSEDMKKDYVREMEDDSRRRREEEGIEVKKCSNQCPSSEEDPINGV
ERQMAEEEVYMDNEEYHTVTGLDSEQFSSFKEEL
>ML26493a
MNGLRLMLLTAALLGATSGADLTSAERNEIDCLRSTHRHYYSIHWDTDLQKQARKTLMKKQGRVARQALEEDFMLDSTWDVAPPKGASVSESNT
LMVRGLCTRYFKVKGSQIKQYQRGQVGCTVEQQNGKRMKIVCHYKRN
>ML01292a
MRGKLWVLVITISVAIVIQMAGKCLISASEGRSGGSRSRKRSVEDAEKSQKLNLKEYGDDGNYGNDGFNGYSRRRSYDYSRRRSYYTSDYDRRR
SYDYSRRRSSYSSGSSTAATLKYVGIGLTVLGVLIGMCKYCAKTCADEDNPVGPGVGATVEPVHTAGESDLQHVQHTENFVEAHVQQNVSYRDE
TPHFSAPPAEFLFAYKVERHDSEDIPAMPPPSYEDVVSPNYTPQ
>ML079714a
MRYLLWTILLVTAVSAKKKKQKCEWSDRPGIYLKNYDRKSKAYPSVEEAKEACEKDSSCGAVGLDLIPYREVPGGPVIHDNTVYLLEGDGPESL
LLGARPNRRYKSYIKVECTEEKKGKRNKEKKNKKDKKDKKEPKTKRSQKKKMI
>ML22358a
MKKTLYTILLLSLFLGIFSKTELALDDATVGHTDCGEPHQVSLETTSCLCLKTLYEGNKCTYSDALKKPFGKECRVGKLEYCCNYIYRLAEWKD
CTSFPY
>ML22359a
MKKIVLLMSVVIASLGSDLENGSEDLQLSQREEDRNVGFTHCGEPYQVSWETSCSCLKTLYEGNRCTYSDAIENPFGKKCKAGKLEYCCNYMYR
LAKKKACTSFPY
>ML05961a
MKFTRVVFVLLLATFVNCELECYTCHADEDPLKDRFISEVCNVKKKCGNYKYCLSMTYKYNGLSYQKRGCMESDQ
>ML059710a
MKFSLLFLLTLLLLAQLSTCRKKKNLGQTEEEETCVIYGKTLDVGESQYMEHICGYCHCRSPQNVICDDTVCEANEVNNSEDEENTCKNRSQWE
IFDSQDGCCRFFCDNTLEKHVLRKLKPECNACS
>ML063320a
MTLRTFSLLLLVSLSLSVSSLPTRSTEEDTPTPPYFSTTDQTNTETNCVSTDGELVAEGDLLYEDCEVRCYCVAGSESCLDRCRRPFPPSNCRL
PRIVPDPRDSCCRVYRCFD
>ML20733a

```
MFAKRVSVTIITMLLSLTSYIEETTFPVDTGTELKVTCKDGFLLKGSDTVTCTDDTTFNSDVPPACVRPACNQTASRLKKLREGMGTTGLTQRH
EDN
>ML049619a
MRTVVVLTLLLGVVYSYCPSHVESRLDQVITKLEMMINSLENEDFSAISFDGDDSQDGTEEPPTPDTSEGDSRTNFLFCVPIRLQNSCPAGYVA
RSGSRDSNGRRSFMCCK
>ML12367a
MKLTLMVLVGLALVVAVHGVCVHNGVEYQTGDSFIRECNKCRCRRNGTSACTRKKCLAEMAVCEHNGNSYFLGEVFKDSCNTCSCSRSLCTQRC
RIPNRRQFHKGMQQVPVSQEWNVRLYQKEMPCRDGGLRAQR
>ML12368a
MKILVLLLMLTLVLVSDGAKKGTKKTCEHNGVTYRPGQKFKDDCNKCKCKKSGTVSCTEKVCDVNICEHDGVIYYAGDSFKADCNTCTCTGRGI
SVCTEMACLGDF
>ML021119a
MNCVSFLFVVTLCSLAFVQGGRLRRGADLKSAEQLLKRHKSRDCGRRDAPECEKWASFGLCSDELYNVHCARTCGFCKKELENEVQLEERHKSR
DCGRRDAPECEKWASFGLCSDELYNVHCARTCGFCKKELENEVQLEESMYRFYLIMVNTLNVNNVIAFGANLSSLRLYQRQFTLCFEYYSY
>ML32998a
MRILEIIPIFVILWEVGIPRAQPDEDTTQDSFACYRCETKADEDICEEKDDTCAYPKCAKPRYAASPNCSRTNLVTPGCVTSPSVTAATASSRR
RSA
```

***Supplementary table 2. NeuroPID scoring for the linear neuropeptide precursors.***

| Protein Name | Extra Trees Prediction | MinEx Tree Prediction | Linear SVM Prediction | Gradient Boosting Prediction | Internal Score |
|---|---|---|---|---|---|
| ML319815a | 0,80 | 0,86 | 0,96 | 1,00 | 25,12 |
| ML14597a | 0,83 | 0,89 | 0,83 | 1,00 | 15,38 |
| ML10665a | 0,81 | 0,88 | 0,96 | 1,00 | 15 |
| ML034332a | 0,87 | 0,88 | 0,99 | 1,00 | 13,03 |
| ML31983a | 0,78 | 0,82 | 0,91 | 1,00 | 11,63 |
| ML11723a | 0,92 | 0,90 | 0,99 | 1,00 | 8,89 |
| ML06743a | 0,88 | 0,82 | 0,96 | 1,00 | 8,63 |
| ML017711a | 0,87 | 0,78 | 1,00 | 1,00 | 8 |
| ML06328a | 0,81 | 0,85 | 0,77 | 1,00 | 7,14 |
| ML21545a | 0,93 | 0,93 | 0,99 | 1,00 | 6,9 |
| ML21632a | 0,88 | 0,83 | 0,96 | 1,00 | 6,67 |
| ML01798a | 0,92 | 0,90 | 0,86 | 1,00 | 6,67 |
| ML040716a | 0,83 | 0,84 | 0,63 | 1,00 | 6,25 |
| ML26664a | 0,80 | 0,77 | 0,97 | 1,00 | 6,25 |
| ML090813a | 0,87 | 0,84 | 1,00 | 1,00 | 5,56 |
| ML31164a | 0,87 | 0,92 | 0,87 | 1,00 | 5,45 |
| ML233326a | 0,84 | 0,80 | 0,82 | 1,00 | 5,41 |
| ML06405a | 0,91 | 0,85 | 0,99 | 1,00 | 5,13 |
| ML016347a | 0,81 | 0,82 | 0,91 | 1,00 | 4,94 |
| ML073260a | 0,91 | 0,93 | 0,67 | 1,00 | 4,44 |
| ML07842a | 0,97 | 0,88 | 0,97 | 1,00 | 4,29 |
| ML124215a | 0,85 | 0,89 | 0,93 | 1,00 | 4,17 |
| ML066512a | 0,86 | 0,92 | 0,32 | 1,00 | 4,12 |
| ML032113a | 0,92 | 0,90 | 0,79 | 1,00 | 4,08 |
| ML065715a | 0,71 | 0,62 | 0,98 | 1,00 | 3,93 |
| ML056913a | 0,85 | 0,87 | 0,76 | 1,00 | 3,77 |
| ML040029a | 0,75 | 0,81 | 0,33 | 1,00 | 3,73 |
| ML08268a | 0,81 | 0,77 | 0,81 | 1,00 | 3,66 |
| ML154515a | 0,85 | 0,94 | 0,43 | 1,00 | 3,17 |
| ML14991a | 0,93 | 0,89 | 0,90 | 1,00 | 2,97 |
| ML46396a | 0,80 | 0,77 | 0,95 | 1,00 | 2,94 |
| ML199816a | 0,83 | 0,78 | 0,93 | 1,00 | 2,86 |
| ML1541126a | 0,84 | 0,93 | 0,68 | 1,00 | 2,76 |
| ML005019a | 0,88 | 0,88 | 0,75 | 1,00 | 2,56 |
| ML065755a | 0,72 | 0,83 | 0,84 | 1,00 | 2,5 |
| ML039810a | 0,73 | 0,79 | 0,90 | 1,00 | 2,47 |
| ML051421a | 0,87 | 0,85 | 0,83 | 1,00 | 2,41 |
| ML174759a | 0,89 | 0,90 | 0,96 | 1,00 | 2,38 |
| ML02212a | 0,83 | 0,89 | 0,95 | 1,00 | 2,33 |
| ML215411a | 0,76 | 0,84 | 0,76 | 1,00 | 2,22 |
| ML093052a | 0,85 | 0,85 | 0,85 | 1,00 | 2 |
| ML33461a | 0,79 | 0,77 | 0,84 | 1,00 | 1,89 |

| | | | | | |
|---|---|---|---|---|---|
| ML216920a | 0,86 | 0,88 | 0,87 | 1,00 | 1,45 |
| ML003517a | 0,88 | 0,88 | 0,88 | 1,00 | 1,27 |
| ML01134a | 0,84 | 0,78 | 0,91 | 1,00 | 1,11 |
| ML218923a | 0,81 | 0,85 | 0,89 | 0,99 | 0,98 |
| ML07361a | 0,81 | 0,85 | 0,97 | 1,00 | 0 |
| ML07012a | 0,80 | 0,85 | 0,95 | 1,00 | 0 |
| ML030512a | 0,85 | 0,85 | 0,90 | 1,00 | 0 |
| ML02736a | 0,82 | 0,85 | 0,82 | 1,00 | 0 |
| ML043317a | 0,83 | 0,81 | 0,93 | 1,00 | 0 |
| ML00497a | 0,79 | 0,81 | 0,85 | 1,00 | 0 |
| ML053619a | 0,77 | 0,81 | 0,89 | 1,00 | 0 |
| ML053618a | 0,77 | 0,81 | 0,89 | 1,00 | 0 |
| ML258215a | 0,80 | 0,80 | 0,98 | 1,00 | 0 |
| ML100012a | 0,80 | 0,77 | 0,99 | 1,00 | 0 |
| ML090812a | 0,83 | 0,78 | 0,96 | 1,00 | 0 |
| ML05859a | 0,76 | 0,85 | 0,87 | 1,00 | 0 |
| ML030510a | 0,81 | 0,79 | 0,90 | 1,00 | 0 |
| ML05367a | 0,76 | 0,80 | 0,85 | 1,00 | 0 |
| ML11691a | 0,77 | 0,71 | 0,93 | 1,00 | 0 |
| ML083811a | 0,59 | 0,60 | 0,87 | 1,00 | 0 |
| ML218823a | 0,76 | 0,78 | 0,91 | 1,00 | 0 |
| ML00992a | 0,71 | 0,79 | 0,96 | 1,00 | 0 |
| ML206415a | 0,76 | 0,80 | 0,87 | 1,00 | 0 |
| ML030511a | 0,76 | 0,78 | 0,84 | 1,00 | 0 |
| ML067017a | 0,65 | 0,73 | 0,87 | 1,00 | 0 |

***Supplementary table 3. Neuropeptide homologs uncovered in other ctenophore species***

| | |
|---|---|
| ML01798a | >Pukia_falcata_sb\|13085131\|<br>>Pukia_falcata_sb\|13047163\|<br>>Dryodora_glandiformis_sb\|284310\|<br>>GHXS01075231.1 TSA: Hormiphora californensis isolate 20161213-T1 tx_DN13442_c0_g3_i1, transcribed RNA sequence<br>>GGLO01069197.1 TSA: Hormiphora californensis TR28038_c0_g1_i2 transcribed RNA sequence |
| ML02212a | >Pleur_sb\|12660472\|<br>>Pukia_falcata_sb\|13062132<br>>Dryodora_glandiformis_sb\|292161<br>>Bolinopsis_infundibulum_sb\|12215413<br>>Bolinopsis_ashleyi_sb\|12973892\|<br>>GHXS01044202.1 TSA: Hormiphora californensis isolate 20161213-T1 tx_DN17776_c0_g1_i1, transcribed RNA sequence<br>>GHXY01365093.1 TSA: Beroe forskalii isolate Bf201507 CTE_B_forskalii.D785-D11.30471c0g1i3, transcribed RNA sequence |
| ML02736a | >Pleur_sb\|12672458\|<br>>Pukia_falcata_sb\|13054685\|<br>>Euplokamis_dunlapae_sb\|10633353\|<br>>Bolinopsis_infundibulum_sb\|12166874\|<br>>Bolinopsis_ashleyi_sb\|12966060\|<br>>GGLO01055400.1 TSA: Hormiphora californensis TR22397_c1_g1_i1 transcribed RNA sequence<br>>GHXY01172047.1 TSA: Beroe forskalii isolate Bf201507 CTE_B_forskalii.D785-D11.25303c0g5i3, transcribed RNA sequence<br>>GHXY01128478.1 TSA: Beroe forskalii isolate Bf201507 CTE_B_forskalii.D785-D11.35947c8g1i4, transcribed RNA sequence<br>>GHXY01128479.1 TSA: Beroe forskalii isolate Bf201507 CTE_B_forskalii.D785-D11.35947c8g1i5, transcribed RNA sequence<br>>GHXY01128476.1 TSA: Beroe forskalii isolate Bf201507 CTE_B_forskalii.D785-D11.35947c8g1i8, transcribed RNA sequence<br>>GGLO01055401.1 TSA: Hormiphora californensis TR22397_c1_g1_i2 transcribed RNA sequence<br>>Beroe_abyssicola_sb\|12131309\|<br>>Beroe_sp_pink_sb\|12859312\| |
| ML07842a | >Pukia_falcata_sb\|13030337\|<br>>Bolinopsis_infundibulum_sb\|12232314\|<br>>Bolinopsis_ashleyi_sb\|12925108\| |
| ML017711a | >Pukia_falcata_sb\|13023494\|<br>>Bolinopsis_infundibulum_sb\|12232315\|<br>>Bolinopsis_ashleyi_sb\|12948034\|<br>>GHXY01131763.1_3 TSA: Beroe forskalii isolate Bf201507 CTE_B_forskalii.D785-D11.29326c5g3i1, transcribed RNA sequence<br>>GGLO01068950.1_2 TSA: Hormiphora californensis TR27976_c4_g1_i2 transcribed RNA sequence<br>>GHXS01104377.1_3 TSA: Hormiphora californensis isolate 20161213-T1 tx_DN17364_c0_g3_i4, transcribed RNA sequence<br>>GGLO01068949.1_2 TSA: Hormiphora californensis TR27976_c4_g1_i1 transcribed RNA sequence |
| ML06743a | >Pleur_sb\|12652162\|<br>>Pukia_falcata_sb\|13031051\|<br>>Euplokamis_dunlapae_sb\|10678971\| |
| ML21545a | >Dryodora_glandiformis_sb\|290240\| |

| | |
|---|---|
| | >Bolinopsis_infundibulum_sb\|12203994\|<br>>Bolinopsis_infundibulum_sb\|12284934\|<br>>Beroe_abyssicola_sb\|12126456\|<br>>Beroe_sp_pink_sb\|12870588\| |
| ML030510a | >Bolinopsis_ashleyi_sb\|12953705\|<br>>Bolinopsis_infundibulum_sb\|12247816\|<br>>Bolinopsis_infundibulum_sb\|12202816\| |
| ML31983a | >Bolinopsis_ashleyi_sb\|12936299\| |
| ML043317 | >P-bachei_GeneM_sb\|11611576\|<br>>Bolinopsis_infundibulum_sb\|12203929<br>>GHXY01322137.1_1 TSA: Beroe forskalii isolate Bf201507 CTE_B_forskalii.D785-D11.36135c9g1i2 transcribed RNA sequence<br>>Beroe_sp_pink_sb\|12908207\| |
| ML056913a | >Pleur_sb\|12812941\|<br>>Pukia_falcata_sb\|13012837\|<br>>Dryodora_glandiformis_sb\|290245\|<br>>Bolinopsis_ashleyi_sb\|12952707\|<br>>GGLO01074549.1_2 TSA: Hormiphora californensis TR29740_c0_g1_i2 transcribed RNA sequence<br>>GHXS01080954.1_1 TSA: Hormiphora californensis isolate 20161213-T1 tx_DN13025_c0_g1_i1 transcribed RNA sequence<br>>GHXY01201788.1_2 TSA: Beroe forskalii isolate Bf201507 CTE_B_forskalii.D785-D11.35563c0g1i1 transcribed RNA sequence<br>>Beroe_abyssicola_sb\|12126338\|<br>>Beroe_sp_pink_sb\|12885168\| |
| ML065755a | >Pleur_sb\|12752694\|<br>>Pukia_falcata_sb\|13044247\|<br>>Bolinopsis_infundibulum_sb\|12202363\|<br>>GHXY01305395.1 TSA: Beroe forskalii isolate Bf201507 CTE_B_forskalii.D785-D11.26122c4g1i3 transcribed RNA sequence<br>>GHXS01080165.1 TSA: Hormiphora californensis isolate 20161213-T1 tx_DN12392_c1_g1_i2 transcribed RNA sequence<br>>Beroe_sp_pink_sb\|12842138\| |
| ML066512a | >Bolinopsis_ashleyi_sb\|12949501\| |
| ML199816a | >Pleur_sb\|12660689\|<br>>Pukia_falcata_sb\|13058854\|<br>>Euplokamis_dunlapae_sb\|10617555\|<br>>Bolinopsis_infundibulum_sb\|12214463\|<br>>Bolinopsis_ashleyi_sb\|12935571\|<br>>GGLO01031268.1 TSA: Hormiphora californensis TR13199_c3_g1_i1 transcribed RNA sequence<br>>Beroe_abyssicola_sb\|12118498\|<br>>Beroe_sp_pink_sb\|12869105\| |
| ML233326a | >Pleur_sb\|12660472\|<br>>Pukia_falcata_sb\|13042259\|<br>>Bolinopsis_infundibulum_sb\|12225922\|<br>>Bolinopsis_ashleyi_sb\|12973892\|<br>>GHXS01067440.1 TSA: Hormiphora californensis isolate 20161213-T1 tx_DN14248_c0_g1_i1 transcribed RNA sequence<br>>GHXY01353421.1 TSA: Beroe forskalii isolate Bf201507 CTE_B_forskalii.D785-D11.23317c0g1i1 transcribed RNA sequence<br>>Beroe_sp_pink_sb\|12857913\| |
| ML218923a | >P-bachei_GeneM_sb\|11609489\|<br>>Pukia_falcata_sb\|13091370\|<br>>GHXS01041156.1 TSA: Hormiphora californensis isolate 20161213-T1 tx_DN18013_c2_g5_i1 transcribed RNA sequence |

***Supplementary table 4. Summary of neuropeptide expression patterns and metacell annotation.***
*The first match between a neuropeptide expression site and a metacell is highlighted in red; such matches correspond to specific expression in particular metacells and were key for the annotation of other metacells.*

| Probe | Localisation | Metacell |
|---|---|---|
| ML065755a | AO | C29, C30 |
| ML206415a | AO | C29, C30 |
| ML14991a | AO | C28, C29, C32, C36, (C39) |
| ML003517a | AO | C32, C36 |
| | AO rim | C27 |
| ML07842a | Pharyngeal neural net | C34 |
| | AO | C40 |
| ML43317a | Sensory neurons gut/mouth | C35 |
| ML030510a, ML030511a | Sensory neurons gut/mouth | C35 |
| | AO rim | C27 |
| ML233326a | Tentacles | C55 |
| ML01798a | Tentacles | C55 |
| ML02212a | Epithelial neural net | C33 |
| | Pharyngeal neural net | C34 |
| | Sensory neurons gut/mouth | C35 |
| | AO | C30 |
| | Tentacle bulb | C33 |
| ML02736a | Epithelial neural net | C33 |
| | Pharyngeal neural net | C34 |
| | Sensory neurons gut/mouth | C35 |
| | AO | C33, C40 |
| | Tentacle bulb | C33 |
| | Tentacles | ? |
| ML199816a | Epithelial neural net | C33 |
| | Pharyngeal neural net | C34 |
| | Sensory neurons gut/mouth | C35 |
| | Tentacle bulb | C33 |
| | Mesogleal neurons inside the tentacle | ? |
| | AO | C33, C28, C32, C36, C40 |
| ML056913a | Epithelial neural net | C33 |
| | Pharyngeal neural net | C34 |
| | Sensory neurons gut/mouth | C35 |
| | Tentacles | C55 |
| | AO | C28, C29, C32, C36 |
| ML21545a | Sensory neurons gut/mouth | C35 |
| | AO | C31 |
| ML17711a | AO | C28, C29, C31, C32, C36, C40 |
| | Epithelial neural net | C33 |
| | Pharyngeal neural net | C34 |
| | Sensory neurons gut/mouth | C35 |

***Supplementary table 5. ISH probes.***

| Gene | Sequence |
|---|---|
| ML00218a | atgaagcttatcgtgttgtttgcgtgtgtctctctggccctggccggaagtaagctgaagaagc<br>tgaagaacgagatggcagatctagaagacaagtttgctactgtgatgaatgacactagcagctc<br>tcttgaggaactccaagcagatgtggatggactcattaccgcgtttaacgagtacaaagaagct<br>caagctaagaaagacatcgacaaccagtggcaagctctgcacatggcagctgctgctgcctgca<br>gaggaagcaccccacaaggaggcaagggaccctggtccaactcggtcgttcctaaagaaaacgg<br>agtgaaatgcgatgaccagtgcagccgtaacgacgacggaaggacactctgcaaggccgaggtc<br>tctctgaccgggtaccctgggaaagccacctcctacacagagcgagtcggacacttctacaact<br>acgagtgtgacggaggctggaacgccgacatcactcacaacgaggttgcggctgaggaggacgc<br>tattcaggttactggtactgcttactacaggttctgctgtt |
| ML016347a | tttgaatttcagggttacagctgtggttctttttgtaattctacttgccgggttttgccctgca<br>gcgccgatatacagtactcaagaaaacgatgcattccttgaaaccggagaagaaaaaggagagc<br>tggatattgaggcaccgaaaacgccaaaacttcgatatttttaccaaaaacgatcagaagaagg<br>caaagatgaggaccctaaacttcaatatttctaccaaatacgatcactagaggacaaagagaag<br>aaacccaaacttcgatacttctaccaaaagcgagaagagaataaagatacgaaacccaaacttc<br>gacacttctaccaaaagcgagaagagaataaagatacaaaacccaaacttcgatacttcttcca<br>aaaacgtgaagagaacaaagatacaaaacccaaacttcgatacttctaccaagagcgagaagat<br>aacaaagatacgaaacccaaacttcgatacttcttccaaaaacgtgaagagaacaaagatacga<br>aacccaaacttcgaaacttctaccgaaaacgaggagaaaact |
| ML07842a | gatgaaactaacggtagtttgcctcctcttcgctagtttgctggtgagggctgaatgcggggtt<br>gttgaaagacacgaggagcagctgtttgatgcggaagatgaacagcccaatttcagagctgggt<br>tagagaataaagaactcgagaaaagaagtccaaactttagatttggatcaaaaaggtcagctga<br>agatgctccaaatctaagaggaaagaggtcaaacgaagataccccccaattttagaggaactaaa<br>agatctgccgaggatagccccaattttagaggagcaaaaagatccgcagtggaagccccaaatt<br>ttagaggagcccaaaaatcagaagaagatgctcctcattttagaggagctaaaagatttgctga<br>ggatgcccccccatttaagaggagtaaagagatccaccgtagatgccccaaattttagaggagca<br>gaaagattagaagaagatgtccctaatttaagaggagcagaaagatccgaagaaggttcccccca<br>attttagaggagcagaaagagcagttgaagatg |
| ML21545a | taagccgggatttaaccgaaggatgtcgaagttcctatttcttactttgctggtgggcgc<br>cgctcgggctgcaagcttcgcttcggactctgacaacactcttgcagactcagacgagtg<br>tagaactggatacaacatgagatacgatctggagtgtaggagaaggagggcagccagggc<br>ggcagccaggttacagctccaggaggatggagagactgaaggaggggaggagctgactaa<br>gagatcggccgaggagcagaactggaattacagcgcaagacgagaccgcggcaaaagatc<br>gaacaacgaggagcaggactttagtggttataggcgtggaagtggacaaaaaaggtctgc<br>tgaagaggacgattacgacatttacaaaagaggggaatcagaagagcaagactactccgg<br>ttatagaggaagggggcgaggagagtagagataacaacaataaattctctaactacacgt<br>gctgggatctcatctctaactacacgtgctggatgtcgtctctagctacacgtactgga<br>tctcatcgctagctacacgtgctggatgtcatcgctaactacaagtgctgggatgtcat<br>cgctaactacacagtacacgtgct |
| ML017711a | aacatttcaagactcacataacctcataaaagatgaagatgtttatattgattggattgc<br>tgatcacactggttctcaactacgtatcagccggagatcttgctagaagatcgcttgagg<br>aggaaaacgagtctggaataaatgaaggagctgctgaggaagatgatgagtctttcaggg<br>gcctccgagaaagcgaggatgagagtaggggtaaaagggaggtcgcagatgaagagctgt<br>tcaggggcaagcgagatttcagggaaaagagagatttcagaggaaaaagggacttcagag<br>gcaagagggaatttagaggcaagagagagttcaggggcaagagagatttccgaggaatga<br>agagcagtgaagaagaattagtaaagagagatctgagtagaggacagtagtaaggatagt<br>attagaagttgaagtgagattatcaggatacatgagctcatttaaagctcgacgttgaa<br>gttttataatgcgatactggtaggaccctgataatgtcctgagtagatgacatatttg<br>atattcaacagctttttttaccttattttagttatagttggtgcagctccgtgcttttc<br>gggatgtacatcatcgcagtcgtgatggcaacaaggccacacctttttgagagaggca |
| ML02212a | ctatcaagtctccaaatctcaacatgaaactcttcctgtttgtccttcttggtctggtggccct<br>gatcagctgcgaaactgtggagtctgaagtggacagcgaacttagcgagagtgaagattctaac<br>gcgatgagggtgaagcgagctaagttcagcatgagcaactacagaggacacaagcagggaaaca<br>gaggctggactggaggtgccatgcaggaggaggagtaggttttatgaaggagcagccacctggt<br>gacgtcatggttacgtcacttagtgacgtcatcgttacgtcacctgaccggcgggaaggaagga<br>ctctttacgaaggagcatgtcttaagtttttaacttattgttaggctaccctggttgattgctg<br>tcttagctggacagattttatcaagtctttactctctcaagatgattttagggattgtcaaata<br>ctttattgcaataatctggacaaatttccaatctagctaattaacatccttcattataaatcca<br>tcacaaaatgtcttcagatt |

| | |
|---|---|
| ML02736a | taagagacagaaagttccagtactttccgagacgaacgaagagaaaaccatgaagtgtttcgtt<br>gtgttgtttgccctcctcgccctatctcagtcagcctcactcaactccctagaatcagtagaag<br>atgtgataatggctgataacgacaacgttgaactggaagaaggtgccctgaacgctgaggagga<br>agcccgagtttacaaaggctacaacggaggtaaccgagtctggtacggataagaagacagacaa<br>ccaacaaggacaagactcaccatgacaacaatcactatggcaacgtcaccatggtaacgtcacc<br>atggcaacgtcaccatggtgacctcatcatggtgccgtgaaatcttgtcgacatggcatcacct<br>tggttacatcttctgtcttcccatcaccacctcacgacgatactgcactcgtgagcttgtccat<br>tttgtcaccgtgttttgtgatttaagaacccggacagtttccacgtttacatgatgagttaatc<br>cattttgaatcttgttaccgtttaaaactgttttcgtaattatatttacaattaatttaaggat<br>tgtgtcactgttattctcgaataataagacaagaaggtg |
| ML056913a | tccatcctccttgtcctcgtcctcgccatcgtgactttcaccagaacaattgaagattctgaag<br>aaatcggacatggactaaggatggataagaacgaggagattggacacggtctgactttagacaa<br>acgaagtgcagacagcgtggacacggaggcaatggctgacgaagaagaactggttggtcacgga<br>atcaaaggatcccatgcgtggaggaagtaaggactgttacggtttgtgacgtcacccaacatca<br>ccatgacgtcactgggtgttgcccatgcgtgttgccgtggtaacgcatgtcctgcaaccttacc<br>attgactttaacaacacgggtcttaaaaagtatgtttggcacgacctgaggatcgttatttaga<br>ttgacaattcttagcaaattttaattagttagcacctcgatgttcatatttggagggtctttgc<br>agagaggataattttattggatgcgattctaagttcaaaatacagttaagattggtatgctctt<br>a |
| ML199816a | tttatagtttgacattcgtatcgatattaattcgaacaacaacaaaatgaagctgttcctcttt<br>ctgaccgcctctctgctcgtgctcgctactgttactgtccagacagaggctagagaaatagtcg<br>cggaggtagccgagtcggaagcagtggagtcggcagcaagtgaagaagacactttcgtgtacag<br>aaaagaggaggactccgcctttctgtttgctgactaatctcacatctgctagagcaacgtcacg<br>gttccactgagagagagacgacgagacagagagataggtctctatttgagacaagcctcttttt<br>gggacaagcctcttttttgagacaagcctctttttgaggcaggtctcttttttgagattaagtctc<br>gctatgaaagccggtactggaaaggactgtgtgtgcttgacttttcatttaatcgcatgggcag<br>tcaattcgcaatgaaatgatacgatacttt |
| ML233326a | aggagctttacagctacacacacacaatgaagaccaccttccttgttctgacccttatgattat<br>ctgctgtaactacgtgcagtctgtaccaatgagtcttgaagaggatctcagcgatgaagagcac<br>cgtggtctacagaagcgagcgggcaccaagtttaacaaggctgactacaaatcagtcggagaag<br>gcaccaggaaatggttcggataatctgccctacctcacctcttttatcagccctcggcgtcagc<br>gtcagaataacctaaactttaaaacagaatgatgaagagcagctttaggactacgcttttaaaa<br>aagacaaaactgtatcattttttaagttcgtgttgatccatgtggcttgctatttaaatttttac<br>gaaatattgaaaatcgcctcgttcaaatgactattttaagtcaattaacagcaaaatacttggt<br>tacgtcttcaagaagt |
| ML065755a | aggatgaaactcgctgctggaatcttcctggttctcgcctgtttaacggttgttatgataccag<br>gcgaagcagcctcattggggagtttagacgctgcaaattcagattctaacgcagtcctgtacgg<br>agctgatcacgaaattttggaatcctcactcatcaaccggaaagaggaagctgagggtgctctg<br>tacggttaaagcgtcagttctactcccacaataaacagattccggacccctctaaccatggacc<br>acaacaacaccacctcccgccctgcagggaacatcttcaaatctgtgacgtcatcattcccgag<br>acttttctcatttgcagctgagtttcggtcgtttaacggaacttatcctacattatatcgcgat<br>atgttcggttatcagcaactctcgttcttagattccgtcgagggcttttaattattttcttttc<br>cgttttgagttttgaagattttagtggtcacatgactggggaaaatgcagggtatttccccaga<br>ggcgagtttcatttagcatccatcatgtgatcacgatggactgggagtgttgctatcaa |
| ML01798a | atgaagctcctactcctaaccctggcagttctcctggcctgcttgactgcagtgcctgtcagac<br>aagaggaggtcccacaagaagagatccgattagaaagatcagctgagagttccggaggagagac<br>agcgagagtggaaaaaagagccgcaatcgatactggttctgattatcccggattcgagggcgga<br>aaacggaggtggtacggttaagaagttctgttcttcttctagtgacgtcctttcctaaagagtg<br>cccaacgaccttcaccgtgatctttctcgaagtagatccctcagtattgctgattggacgcaga<br>cttggtgaaatcagactttattgttcaaaatgataatcaggcgctctttatagttgtttatctt<br>ccatttgcaattggttggctctataatagtattgcaaacatacaattctgattgaaatcttatg<br>catatgttaaaatacccgtatgttgctaacttaccttaatcgatttaaaggagcatctttgatc<br>taagattc |
| ML030510a | ttctctagacggaagttcagacaaacacaaaaatcaaactcaacatgaagttcacaaaactgat<br>catcctgatgactctattggccctcatcgcctcacgatcactagaagaagacaacaacgcagac<br>caaacaattgggtggggccgccagcgagtcgctgaagcggaagacttcgtcgccgaggagaaga<br>tggctgggggaggaaacatgatcctggagatgtcggaggaggagcagcatctcggactgggctt<br>ctaactgaaccggaacatttggaaagataatcttggagagtgatattagtaaaattgtaatcat<br>aatttggcggttaggtttattagttagatacacatttttcatctgaagctagagtaagggtgcg<br>tttaacccaaaaatatattaaaattttctgggtaaaatccatgctaattactgcactagagcta |

| | |
|---|---|
| | tcgcaggtttgaagtccatagaacggtatttgctcgccattttaattttagaatttagaaagtt<br>aagaaagatatttgagtgatctaacgttgattcttt |
| ML14991a | caaaaaggctcagttcccaccagcactactgatttgtctgtctctcaaagttacctctacaaat<br>gtcctgtctcagtaatctctctccttgcaaatcgctactgctgttcctcggtcttgtattattg<br>acgaaggaaagcgttcagcgaacattgccagaatcgggtcaagcattggggatgaaaaacacga<br>tcgaagaggatgctacagtgatgtcagacagcgatgaagagcagtcgaggatgaaacgacagtt<br>tttcagactcaaaagacaatcttttcgtccatcaagaggggatgctgaatggaatcttgacggc<br>tactttggaaaccaggagaaaagtgaagctgagttgagggaccctgacacacaaactagacggt<br>ttaaggacggaaggaacgaacagttaagtcaatggagagcgatgatgaatagggtttgataata<br>gggacttttaacatgtttcatgggagcttagttggattttgctttg |
| ML030511a | aggtcctatgaacgcataaagacaagtaaacaagttattcctgtcatggcgctcactaagattc<br>ttttctctctcagcctcattctgatggtctcttcaagagctttcgaggagtcggctggagatga<br>gaataatcaggcatttgggttgggaaacaaagatggtgcagtaggagacagcctagaggagatt<br>acacagcttgggaaagtagatgagtttaaactggctgaagctgaagatgaccaacatttcgctg<br>tcggaacgtagtgagatctcaaggaaaaacgatggttttaaaagaagattgagccaaaatccta<br>tcgtgtgatccgaataactggcaaatcaggaatattgtattttagccacggacattaatatgat<br>atatgtatgaacgattgtgacaggaaa |
| ML003517a | ttccagctgaacatggctactatgatgggctttttgatagtgctgcttatagccgcagtgtcag<br>ggcacagtcacagtgacgagagtcctcaacacaacagacctgacagtccaccaaaacatcacat<br>ccctaaagatgtcattgaggcacttcagagaagcggaaaaagtcgtggagaacataatgctcac<br>cctggaccagcagaacatcatcctgaaggggctcattcttcacatcttggcaagggacacaatt<br>tcgaacgacaccccgaaggagggaggccaataccccccaaagagggttcgccagagaatttccag<br>attgtagtcatttcactgattcatttaacagatttggccagcaaagtaaaggtgttaactgtgg<br>gctgggatgcttgtgttcgttagccagttattcaactacatgtaactgtattctataaactata<br>gtatgtaatcattaattgtactaagtagtatgtgattatgtagatcgaaagaaggcattttgtt<br>atttaacaataaataatgttttgcactgtattttgttctttttccctgtcttgaatttgtaaaa<br>ttgttaatagtaagcaatggaacagaaagatagtga |
| ML206415a | aattacaatgaagctgcacctctacatcgggacgatcctcctttgtctgagcgccatgttcctt<br>gacggcatcgcccaaaatgatgacagtgcggacgataacgccgccggccagacagaccaggctg<br>gagcggagaacagcgaggatgacgccgccgccgccgccggggaggagggagcagagaccgccga<br>cggagctgagaccaatagtaacgacactactggagcaggtgcggagggagaggctgccgagggc<br>gatagtaaagatccaaacggcggggccatgtcaaacttaaagaattcctatcttcttcttctca<br>cgatccctgtcgttggacgaatattcgtgtagaattgcagtagaagttcatcagttacgtcgtc<br>aacgatgtacaacgacaatcgttcattgtatttgtaaagaggtagtttagactgtaggactcat<br>agtttctttttcttcacaaatatttgtatgttcttgtatttgtattttcaaaatttagtgacgc<br>aattagtacaattttcatattttgacttaactgttgaggagctgt |
| ML43317a | ATGAAGACGATTTTGCTGATATCTTGCCTTCTGTCCGCTGTATACAGTCGAGTGCTTCG<br>TTTAAATGAAGAAGATTGGTCCAGAAATGGAGCTATGGAAGACAGCGAAGATTGGTCCA<br>GAAATGGAGCTATGAGGGATAGCGAAGATTGGAACAGAAATGGTGGTATGAGAGACAGC<br>GAAGATTGGTCTAGAAATGGTCGTATGGAAGACAGTGAAGATTGGTCCAGAAATGGAGC<br>TATGGAAGACAGTGAAGATTGGTCCAGAAATGGAGCTATGGAAGACAGTGAAGAATGGT<br>CTAGAAATGGGGGTTTGGAAAACAGTGAAGATTGGTCCAGAAATGGGGCTATGAGGGAT<br>AGCGAAGATTGGTCAAGAAATGGTGCTATAAAGGATAGTGAAGATTGGTCCAGAAATGG<br>TGCTATGAGGGATAGCGAAGATTGGTCAAGAAATGGTGCTATGAAGGATAGTGAAGATT<br>GGTCCAGAAATGGTATGGAAGACAGTGAAGATTGGTCAAGAAATGGTGCATAGTGAAAT<br>GCATCTGTG |

***Supplementary tables 6-9. Provided as separate Excel sheets.***
